# Genome-wide rare variant analysis for thousands of phenotypes in 54,000 exomes

**DOI:** 10.1101/692368

**Authors:** Elizabeth T. Cirulli, Simon White, Robert W. Read, Gai Elhanan, William J Metcalf, Karen A. Schlauch, Joseph J. Grzymski, James Lu, Nicole L. Washington

## Abstract

Defining the effects that rare variants can have on human phenotypes is essential to advancing our understanding of human health and disease. Large-scale human genetic analyses have thus far focused on common variants, but the development of large cohorts of deeply phenotyped individuals with exome sequence data has now made comprehensive analyses of rare variants possible. We analyzed the effects of rare (MAF<0.1%) variants on 3,166 phenotypes in 40,468 exome-sequenced individuals from the UK Biobank and performed replication as well as meta-analyses with 1,067 phenotypes in 13,470 members of the Healthy Nevada Project (HNP) cohort who underwent Exome+ sequencing at Helix. Our analyses of non-benign coding and loss of function (LoF) variants identified 78 gene-based associations that passed our statistical significance threshold (p<5×10-9). These are associations in which carrying any rare coding or LoF variant in the gene is associated with an enrichment for a specific phenotype, as opposed to GWAS-based associations of strictly single variants. Importantly, our results do not suffer from the test statistic inflation that is often seen with rare variant analyses of biobank-scale data because of our rare variant-tailored methodology, which includes a step that optimizes the carrier frequency threshold for each phenotype based on prevalence. Of the 47 discovery associations whose phenotypes were represented in the replication cohort, 98% showed effects in the expected direction, and 45% attained formal replication significance (p<0.001). Six additional significant associations were identified in our meta-analysis of both cohorts. Among the results, we confirm known associations of *PCSK9* and *APOB* variation with LDL levels; we extend knowledge of variation in the *TYRP1* gene, previously associated with blonde hair color only in Solomon Islanders to blonde hair color in individuals of European ancestry; we show that *PAPPA*, a gene in which common variants had previously associated with height via GWAS, contains rare variants that decrease height; and we make the novel discovery that *STAB1* variation is associated with blood flow in the brain. Our results are available for download and interactive browsing in an app (https://ukb.research.helix.com). This comprehensive analysis of the effects of rare variants on human phenotypes marks one of the first steps in the next big phase of human genetics, where large, deeply phenotyped cohorts with next generation sequence data will elucidate the effects of rare variants.

## Introduction

Over the past decade, we have witnessed the growing depth and breadth of genome-wide association studies (GWAS) leveraging genotyped common variants. We have seen that the most useful and predictive insights about the genetic effects of common variants only begin to appear as sample sizes reach into the hundreds of thousands. Modern resources like the UK Biobank (UKB, www.ukbiobank.ac.uk) that make thousands of phenotypes available to match these genetic data are proving a boon to our understanding of human genetics. In addition to identifying specific variants associated with traits, modern GWAS have shown that polygenic scores utilizing thousands of common variants together can explain a sizeable portion of phenotypic variation and that genetic risk for one phenotype can help explain variation in another^1–3^.

Until now, the insights stemming from these large sample sizes have only been available for common and low frequency variants, with comprehensive studies only available to a minor allele frequency (MAF) of about 0.1%. It has been repeatedly shown that as allele frequencies drop, the effect sizes of these variants increase beyond the limits imposed by natural selection on more common variants^4–6^. In rare diseases and family-based studies, rare variant studies that aggregate phenotypically similar probands have been crucial to our understanding of disease; exome and genome-based approaches are now standard of care for evaluating these patients. However, the impact of rare variants on common traits and sub-clinical phenotypes has only been examined for selected phenotypes as large exome and phenotypic datasets have not been available.

The release by the UKB of 49,960 exomes matched to thousands of phenotypes finally changes this status quo and marks the beginning of the next era in the study of human genetics^7^. While the sample size remains modest compared to modern chip-based GWAS, this is the first time that rare variants and corresponding phenotypes can be analyzed by researchers around the world on such a scale. In addition to this incredible public resource, we have sequenced the exomes of 18,102 participants in the Healthy Nevada Project (HNP, Renown Health, Reno, Nevada) who consented to research involving their electronic medical records. We therefore can not only perform analyses of rare variants against thousands of phenotypes in the UKB dataset but can also perform independent replication analyses to confirm associations and meta-analysis to discover new signals.

Population-based analyses that aim to identify statistically significant associations between traits and rare variants require a different methodology from the common variant methods to which the field has grown accustomed ^8,9^. The power to identify rare variants as statistically significant associations decreases as the minor allele frequency decreases. This is why the methodology for rare disease and private mutations require aggregation of similar probands and aggregation of variants at the gene level. For population-based genetic analyses of rare variants, it is therefore essential to group the signals of multiple rare variants together, such as by co-occurrence in the same gene, to increase statistical power (Figure 1). This method has been used in exome and genome sequencing studies to successfully identify genes associated with many traits, including myocardial infarction, amyotrophic lateral sclerosis, and blood pressure^10–12^.

**Figure 1.**
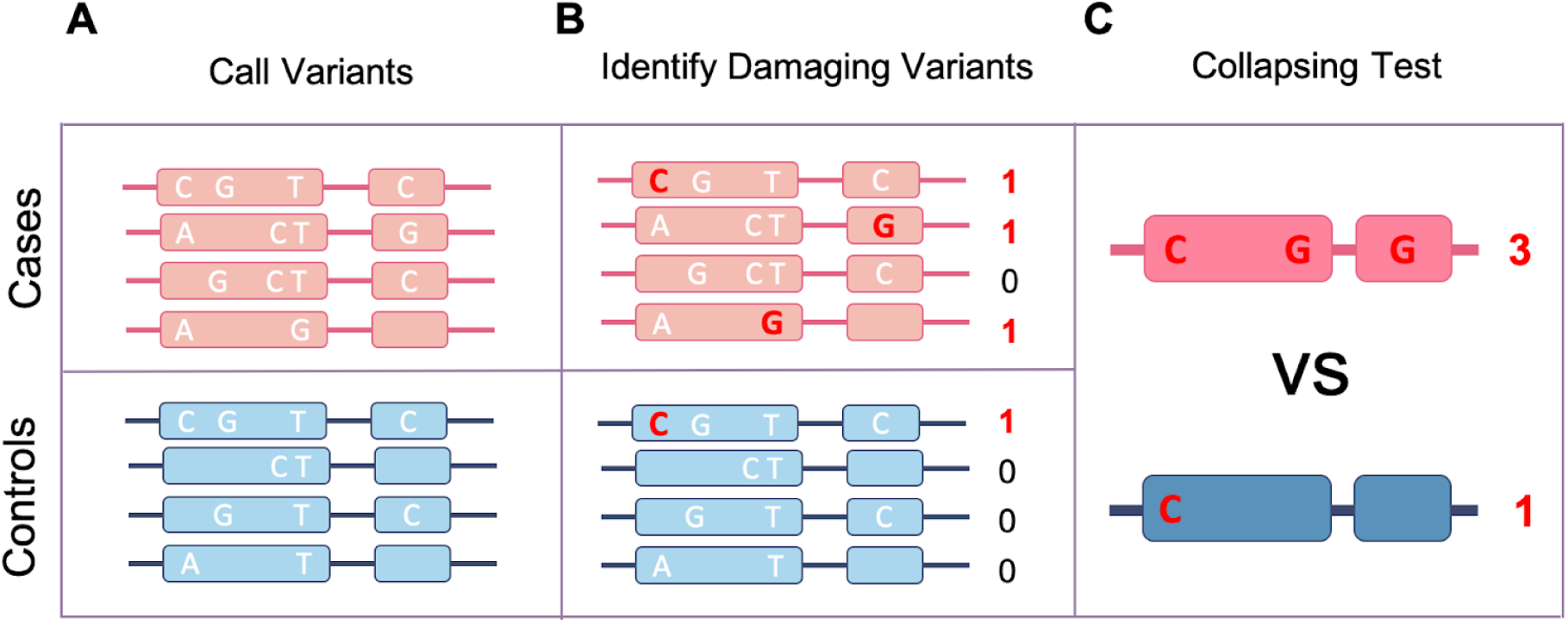
Gene-based collapsing analysis. A) First, variants in each gene are identified by sequencing in cases and controls. B) Variants that are predicted to be damaging—those that are rare and annotated as likely to affect the functionality of the gene, such as coding variants—are then selected for analysis, while variants that are common or do not have damaging annotations are excluded from further analysis. The exact parameters used to select variants can be flexible and tailored to each study. C) Finally, the number of cases with a qualifying variant in each gene is compared to the number of controls with a qualifying variant. This comparison of cases to controls produces one statistical result per gene instead of one per variant.

Here, we apply a gene-based collapsing analysis method to 40,468 participants over 3,166 phenotypes measured by the UKB. We perform replication and meta-analyses for 1,067 traits in an additional 13,470 participants from the Healthy Nevada Project (HNP) cohort. This analysis is the first to make novel rare-variant discoveries by combining tens of thousands of exomes with thousands of phenotypes across multiple cohorts.

## Results

### Gene-based collapsing discovery analysis

We performed a gene-based collapsing analysis to identify genes in which rare variants were in aggregate associated with a phenotype. In brief, we identified qualifying variants that met specific annotation criteria (see Methods) and were below a MAF of 0.1%. We explored two gene-based collapsing models: 1) coding and 2) loss of function (LoF). The LoF model was used to identify associations where only LoF variants had an effect. For the coding model, we included 747,865 qualifying variants across 15,474 genes (see Methods). In the LoF model, we included 115,628 qualifying variants across 8,307 genes. The mean number of qualifying variants per gene in the discovery population was 46; the mean percentage of carriers for each gene was 0.38%.

We analyzed 3,166 phenotypes in the 40,468 individuals who were classified by the UKB as genetically Caucasian (field 22006) (Table 1, see Supplement for list of phenotypes). As described previously, the exome-sequenced set of UKB samples is enriched for individuals with MRI data, enhanced baseline measurements, hospital episode statistics, and linked primary care records (described for Category 170 at http://biobank.ctsu.ox.ac.uk/crystal/label.cgi?id=170). We included relatives in the analysis, accounting for their relatedness with a linear mixed model. To reduce test statistic inflation, genes were only included in analyses of quantitative traits if they had at least 10 phenotyped people carrying a qualifying variant. For binary traits, the expected number of carriers in the case group was required to be at least 10, based on the overall carrier and phenotype frequency^13^. We identified 78 associations that were statistically significant (p<5×10^-9^) after applying the Bonferroni multiple testing correction (Tables 2-3). The vast majority of these associations were expected given the current knowledge in the field. For example, rare variants in *PCSK9* and *APOB* were associated with low density lipoprotein (LDL) levels, and rare loss of function variants in *TUBB1* were associated with platelet distribution width.

**Table 1.** Study and cohort information. Except for total individuals, counts shown are for the European ancestry subset.

|  | UK Biobank (UKB) | Healthy Nevada Project (HNP) |
| --- | --- | --- |
| N individuals: total / European ancestry | 49,960 / 40,468 | 18,102 / 13,470 |
| N phenotypes: binary / quantitative | 1,917 / 1,249 | 916 / 151 |
| N phenotypes unique to cohort: binary / quantitative | 1,189 / 1,198 | 188 / 100 |
| Median N cases for binary traits (range) | 257 (50-37,983) | 181 (24-6,414) |
| Median N phenotyped for quantitative traits (range) | 9,287 (516 - 40,428) | 1,440 (190-10,244) |

**Table 2.** Statistically significant associations with binary traits in the UKB gene-based collapsing analysis.

| Gene | Model | Phenotype | Case carrier n (%) | Ctrl carrier n (%) | p-value | Direction | Replication |
| --- | --- | --- | --- | --- | --- | --- | --- |
| *SLC45A2* | coding | Hair colour: Blonde | 62 (1.33%) | 117 (0.33%) | 3.70E-27 | + | NA |
| *LDLR* | coding | High cholesterol | 75 (1.46%) | 170 (0.48%) | 7.50E-17 | + | NA |
| *COL4A4* | coding | R31 Unspecified haematuria | 40 (2.7%) | 328 (0.84%) | 5.50E-14 | + | consistent |
| *MC1R* | coding | Hair colour: Red | 29 (1.63%) | 213 (0.55%) | 4.40E-11 | + | NA |
| *TYRP1* | coding | Hair colour: Blonde | 55 (1.18%) | 175 (0.49%) | 4.60E-11 | + | NA |
| *LDLR* | coding | Cholesterol lowering medication | 45 (1.61%) | 91 (0.48%) | 1.20E-10 | + | NA |
| *TTN* | LoF | I48 Atrial fibrillation and flutter | 38 (2.2%) | 291 (0.75%) | 1.20E-10 | + | consistent |
| *SLC45A2* | coding | Hair colour: Dark brown | 38 (0.25%) | 141 (0.56%) | 4.60E-10 | - | NA |

Direction of effect is + for betas above 0 and - for betas below 0. "Consistent" means that the beta was in the same direction for both UKB and HNP but with $p > 0.001$ . Consistent betas with $p < 0.001$ would be labeled ' $p < 0.001$ '.

**Table 3.** Statistically significant associations with quantitative traits in the UKB gene-based collapsing analysis.

| Gene | Model | Phenotype | Carrier n (%) | p-value | Direction | Replication |
| --- | --- | --- | --- | --- | --- | --- |
| *ALPL* | coding | Alkaline phosphatase | 249 (0.66%) | 2.30E-187 | - | p<0.001 |
| *SLC22A12* | coding | Urate | 313 (0.83%) | 9.90E-119 | - | consistent |
| *GOT1* | coding | Aspartate aminotransferase | 89 (0.24%) | 1.50E-68 | - | p<0.001 |
| *CST3* | LoF | Cystatin C | 23 (0.06%) | 7.70E-53 | - | NA |
| *CST3* | coding | Cystatin C | 56 (0.15%) | 8.00E-53 | - | NA |
| *TUBB1* | coding | Platelet distribution width | 229 (0.58%) | 7.40E-45 | + | NA |
| *ABCA1* | coding | Apolipoprotein A | 429 (1.21%) | 6.80E-41 | - | NA |
| *GPLD1* | coding | Alkaline phosphatase | 317 (0.84%) | 4.10E-36 | - | p<0.001 |
| *ABCA1* | coding | HDL cholesterol | 430 (1.2%) | 4.40E-36 | - | p<0.001 |
| *SHBG* | coding | SHBG | 148 (0.42%) | 5.70E-33 | - | NA |
| *SHBG* | LoF | SHBG | 82 (0.23%) | 7.50E-32 | - | NA |
| *APOB* | LoF | LDL direct | 90 (0.24%) | 2.10E-31 | - | p<0.001 |
| *GPT* | coding | Alanine aminotransferase | 124 (0.33%) | 2.00E-28 | - | consistent |
| *PCSK9* | coding | LDL direct | 205 (0.54%) | 2.00E-27 | - | p<0.001 |
| *PCSK9* | coding | Apolipoprotein B | 206 (0.55%) | 1.70E-25 | - | NA |
| *APOB* | LoF | Cholesterol | 90 (0.24%) | 3.40E-25 | - | p<0.001 |
| *TUBB1* | LoF | Platelet distribution width | 24 (0.06%) | 3.90E-24 | + | NA |
| *SLC2A9* | coding | Urate | 139 (0.37%) | 8.00E-23 | - | consistent |
| *PCSK9* | coding | Cholesterol | 207 (0.55%) | 1.60E-22 | - | consistent |
| *APOB* | LoF | Apolipoprotein B | 71 (0.19%) | 2.70E-21 | - | NA |
| *JAK2* | coding | Platelet count | 271 (0.69%) | 2.40E-20 | + | consistent |
| *PCSK9* | LoF | LDL direct | 81 (0.21%) | 7.70E-20 | - | consistent |
| *PCSK9* | LoF | Apolipoprotein B | 81 (0.21%) | 1.30E-19 | - | NA |
| *ALB* | LoF | Albumin | 11 (0.03%) | 1.80E-19 | - | consistent |
| *JAK2* | coding | Platelet crit | 271 (0.69%) | 1.90E-19 | + | NA |
| *ASGR1* | coding | Alkaline phosphatase | 123 (0.32%) | 5.60E-19 | + | consistent |
| *ALPL* | LoF | Alkaline phosphatase | 68 (0.18%) | 7.70E-17 | - | consistent |
| *TUBB1* | coding | Platelet count | 229 (0.58%) | 2.30E-16 | - | p<0.001 |
| *ABCA1* | LoF | Apolipoprotein A | 72 (0.2%) | 5.30E-16 | - | NA |
| *PCSK9* | LoF | Cholesterol | 81 (0.21%) | 8.10E-16 | - | consistent |
| *ALPL* | coding | Phosphate | 236 (0.66%) | 1.10E-15 | + | NA |
| *LCAT* | coding | HDL cholesterol | 88 (0.25%) | 1.80E-15 | - | p<0.001 |
| *KLF1* | LoF | Red blood cell (erythrocyte) distribution width | 27 (0.07%) | 1.10E-14 | + | NA |
| *KLF1* | LoF | Mean corpuscular haemoglobin | 27 (0.07%) | 1.30E-14 | - | p<0.001 |
| *STAB1* | LoF | Median T2star in putamen (right) | 35 (0.37%) | 4.00E-14 | + | NA |
| *ABCB11* | coding | Alkaline phosphatase | 175 (0.46%) | 5.40E-14 | + | consistent |
| *APOB* | LoF | Triglycerides | 89 (0.23%) | 6.60E-14 | - | p<0.001 |
| *GP9* | coding | Mean platelet volume | 84 (0.21%) | 9.00E-14 | + | p<0.001 |
| *ABCA1* | LoF | HDL cholesterol | 72 (0.2%) | 1.30E-13 | - | p<0.001 |
| *IQGAP2* | LoF | Mean platelet volume | 160 (0.41%) | 1.30E-13 | + | No effect |
| *STAB1* | coding | Median T2star in putamen (right) | 223 (2.33%) | 1.40E-13 | + | NA |
| *KLF1* | LoF | Mean corpuscular volume | 27 (0.07%) | 1.50E-13 | - | p<0.001 |
| *MPL* | coding | Platelet count | 242 (0.62%) | 2.80E-13 | + | consistent |
| *GCK* | coding | Glycated haemoglobin | 54 (0.14%) | 4.60E-13 | + | consistent |
| *ANGPTL3* | coding | Apolipoprotein A | 158 (0.45%) | 6.10E-13 | - | NA |
| *GPLD1* | LoF | Alkaline phosphatase | 75 (0.2%) | 3.80E-12 | - | p<0.001 |
| *STAB1* | coding | Median T2star in putamen (left) | 223 (2.33%) | 5.00E-12 | + | NA |
| *LCAT* | coding | Apolipoprotein A | 88 (0.25%) | 6.70E-12 | - | NA |
| *APOA5* | LoF | Triglycerides | 42 (0.11%) | 1.00E-11 | + | consistent |
| *STAB1* | LoF | Median T2star in putamen (left) | 35 (0.37%) | 1.30E-11 | + | NA |
| *CETP* | LoF | HDL cholesterol | 36 (0.1%) | 1.50E-11 | + | p<0.001 |
| *GOT1* | LoF | Aspartate aminotransferase | 10 (0.03%) | 1.50E-11 | - | NA |
| *GP9* | coding | Platelet count | 84 (0.21%) | 2.60E-11 | - | consistent |
| *GP1BB* | coding | Mean platelet volume | 31 (0.08%) | 2.70E-11 | + | NA |
| *GCK* | coding | Glucose | 48 (0.13%) | 4.40E-11 | + | consistent |
| *ITGA2B* | coding | Platelet count | 321 (0.82%) | 4.60E-11 | - | consistent |
| *ASGR1* | LoF | Alkaline phosphatase | 26 (0.07%) | 6.60E-11 | + | consistent |
| *ANGPTL3* | coding | Triglycerides | 168 (0.44%) | 8.40E-11 | - | consistent |
| *CRP* | coding | C-reactive protein | 65 (0.17%) | 9.90E-11 | - | consistent |
| *TUBB1* | coding | Mean platelet volume | 229 (0.58%) | 1.30E-10 | + | consistent |
| *IQGAP2* | coding | Mean platelet volume | 586 (1.49%) | 2.10E-10 | + | consistent |
| *LIPC* | coding | Apolipoprotein A | 248 (0.7%) | 3.40E-10 | + | NA |
| *MPL* | coding | Platelet crit | 242 (0.62%) | 4.20E-10 | + | NA |
| *SCARB1* | coding | HDL cholesterol | 146 (0.41%) | 1.60E-09 | + | p<0.001 |
| *PAPPA* | LoF | Standing height | 11 (0.03%) | 1.90E-09 | - | consistent |
| *CETP* | coding | HDL cholesterol | 169 (0.47%) | 2.00E-09 | + | p<0.001 |
| *TUBB1* | LoF | Platelet count | 24 (0.06%) | 3.00E-09 | - | p<0.001 |
| *GFI1B* | coding | Mean platelet volume | 109 (0.28%) | 3.00E-09 | + | p<0.001 |
| *GP1BA* | LoF | Mean platelet volume | 85 (0.22%) | 3.50E-09 | + | p<0.001 |
| *TUBB1* | coding | Platelet crit | 229 (0.58%) | 3.90E-09 | - | NA |

Direction of effect is + for betas above 0 and - for betas below 0. "Consistent" means that the beta was in the same direction for both UKB and HNP but with $p>0.001$ . Consistent betas with $p<0.001$ are labeled 'p<0.001'.

We observed a number of expected associations that could reasonably be expected given the knowledge in the field but had not been previously identified using these analysis techniques. For example, we found that rare coding variants in *GP1BB* were associated with higher mean platelet volumes in the general population, consistent with their previous association with some bleeding and platelet disorders^14^. As another example, we identified associations between rare coding variants in *TYRP1* and blonde hair. A variant in this gene had previously been shown to cause blonde hair in dark-skinned individuals of Melanesian ancestry from the Solomon Islands, but until now there has not been evidence of a role for this gene in those of European ancestry^15,16^.

Additional discoveries were novel. For example, we found that rare coding variants in *STAB1* were associated with median T2star MRI measures in several brain structures, with the strongest association in the putamen. As *STAB1* is a transmembrane receptor that is thought to play a role in angiogenesis, this finding provides novel hypotheses for further study. As another example, LoF variants in the gene *PAPPA* were associated with decreased height. While this gene was implicated in a previous GWAS of height, this is the first time that rare variants in this gene have been found to be significantly associated with height in a human population^17^.

Of the 3,166 phenotypes chosen for analysis, 298 were unable to be analyzed in the same way because their trait heritability, based on the common variants used in the linear mixed model, fell below the required threshold for the algorithm (see Methods). These phenotypes were analyzed using logistic regression in the subset of unrelated European ancestry individuals and produced no statistically significant associations.

### Replication study and meta analysis

We analyzed 1,067 phenotypes in 13,470 individuals from the HNP cohort who had been sequenced at Helix and volunteered their medical records and blood biomarkers for analysis. For our replication study, 47 of the phenotype/gene combinations of the 78 that were statistically significant were able to be assessed in the HNP cohort. Of these 47, 98% showed directions of effect that were consistent with the discovery signal, and 45% of the 47 achieved statistically significant replication (p<0.001). The only hit that did not replicate was the association between LoF variants in *IQGAP2* and mean platelet volume (though the coding model that included the LoF variants showed a consistent though weak effect in HNP).

We next performed a meta-analysis to identify associations that only achieved statistical significance when combining the signals from the two separately analyzed cohorts. We identified six new associations (p<5×10^-9^), each of which also achieved nominally significant associations (p<0.01) in the analysis of individual cohorts (Table 4).

**Table 4.**
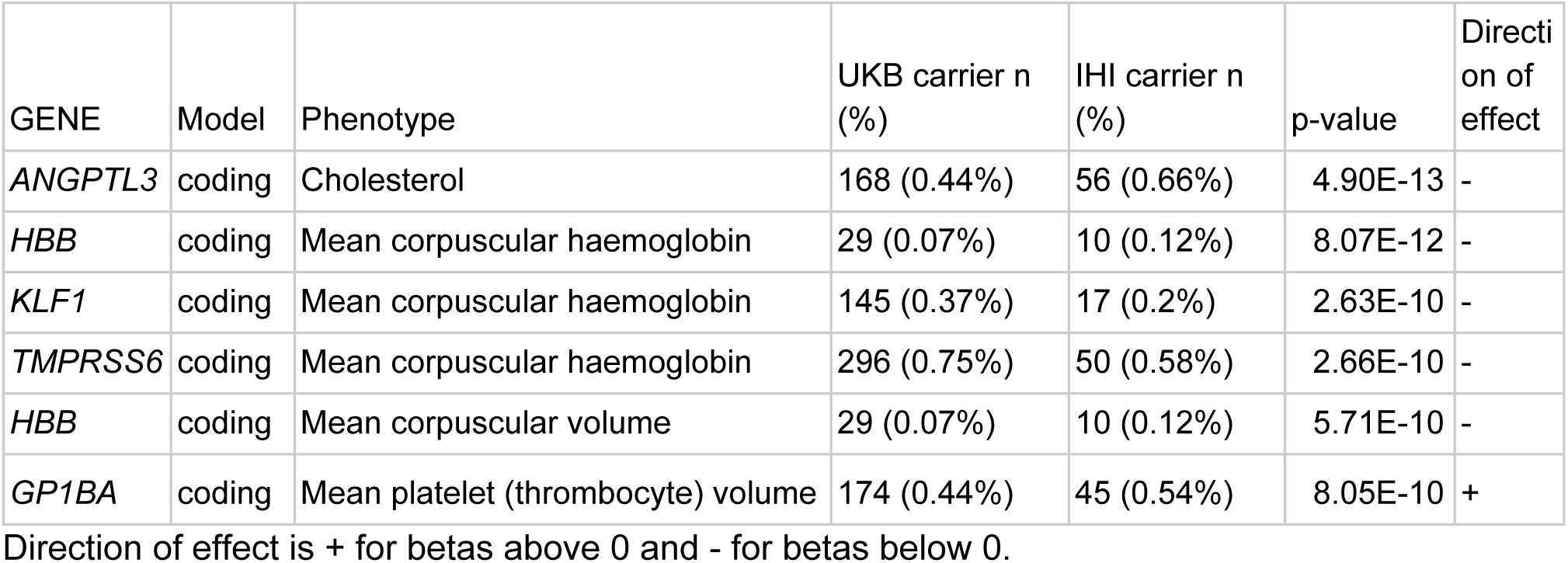
Statistically significant associations from the meta analysis of UKB and HNP.

### Individual variant analysis

We additionally analyzed the individual variants that went into the collapsing analysis, with the intent to identify which genetic variants were driving the signals. Of the 84 significant gene-based associations, only 5% could be explained by a single variant in the gene. Of the 27 associations that formally replicated in the HNP cohort or were significant from the meta analysis of both cohorts together, only one was explained by a single variant. The remaining associations had contributions toward significance from multiple variants, highlighting the utility of grouping together rare variants to improve power for discovery. In fact, the signal for these associations was sufficiently dispersed over multiple variants per gene such that had the analysis been done by traditional GWAS of analyzing one variant at a time, 78% of the gene-based associations found here would not have had a single variant whose individual p-value was sufficient to pass the threshold for correction for multiple tests.

Mapping the precise effects of each contributing variant can elucidate the underlying biology of an association. For example, variants in *SLC2A9* are associated with low urate levels (Figure 3A, B). The protein encoded by this gene reabsorbs urate in the proximal tubules of the kidneys, and variants that disrupt the transmembrane regions or lower gene expression are known to be associated with hypouricemia^18^. We find that the association signal in this gene is most heavily concentrated in missense variants in the transmembrane regions of the protein, especially in the first half of the protein (Figure 3A,B). Of the >30 variants associated with increased urate levels, 89% are in or directly adjacent to a predicted transmembrane region.

**Figure 2.**
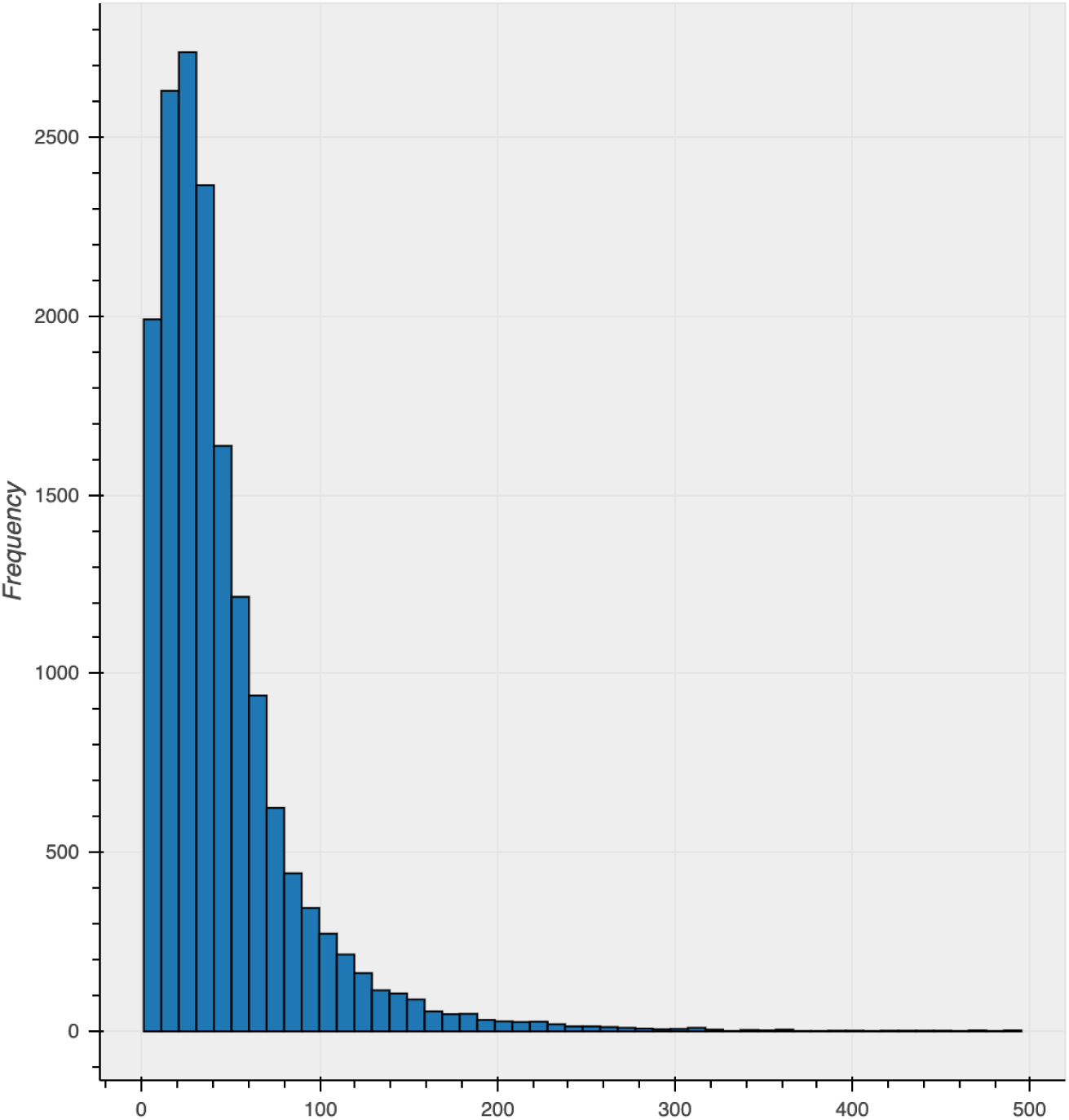
Histogram of number of qualifying variants per gene. Eleven genes with >500 variants excluded from plot. Mean of 45.7 variants per gene (standard deviation 52.2).

**Figure 3.**
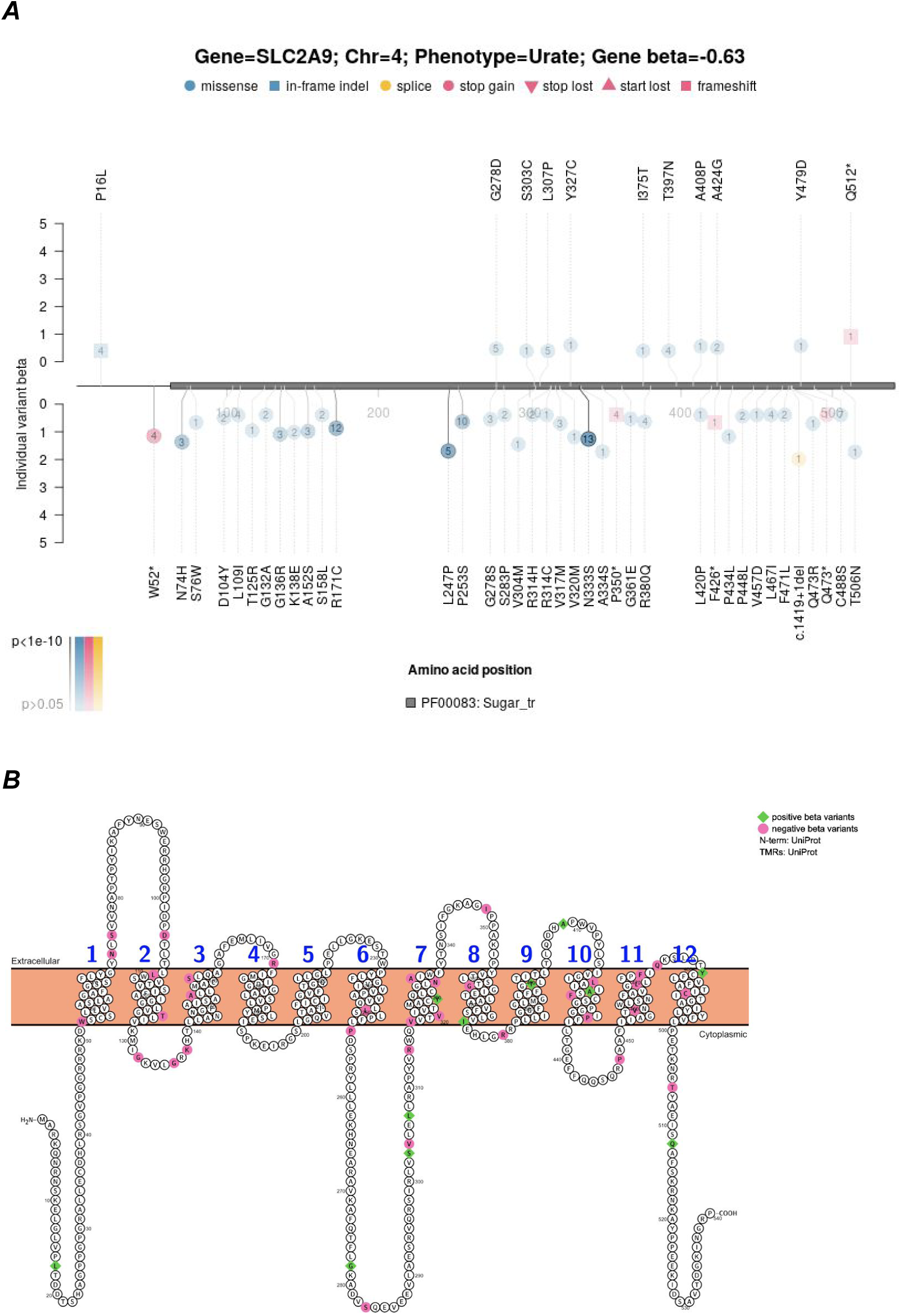
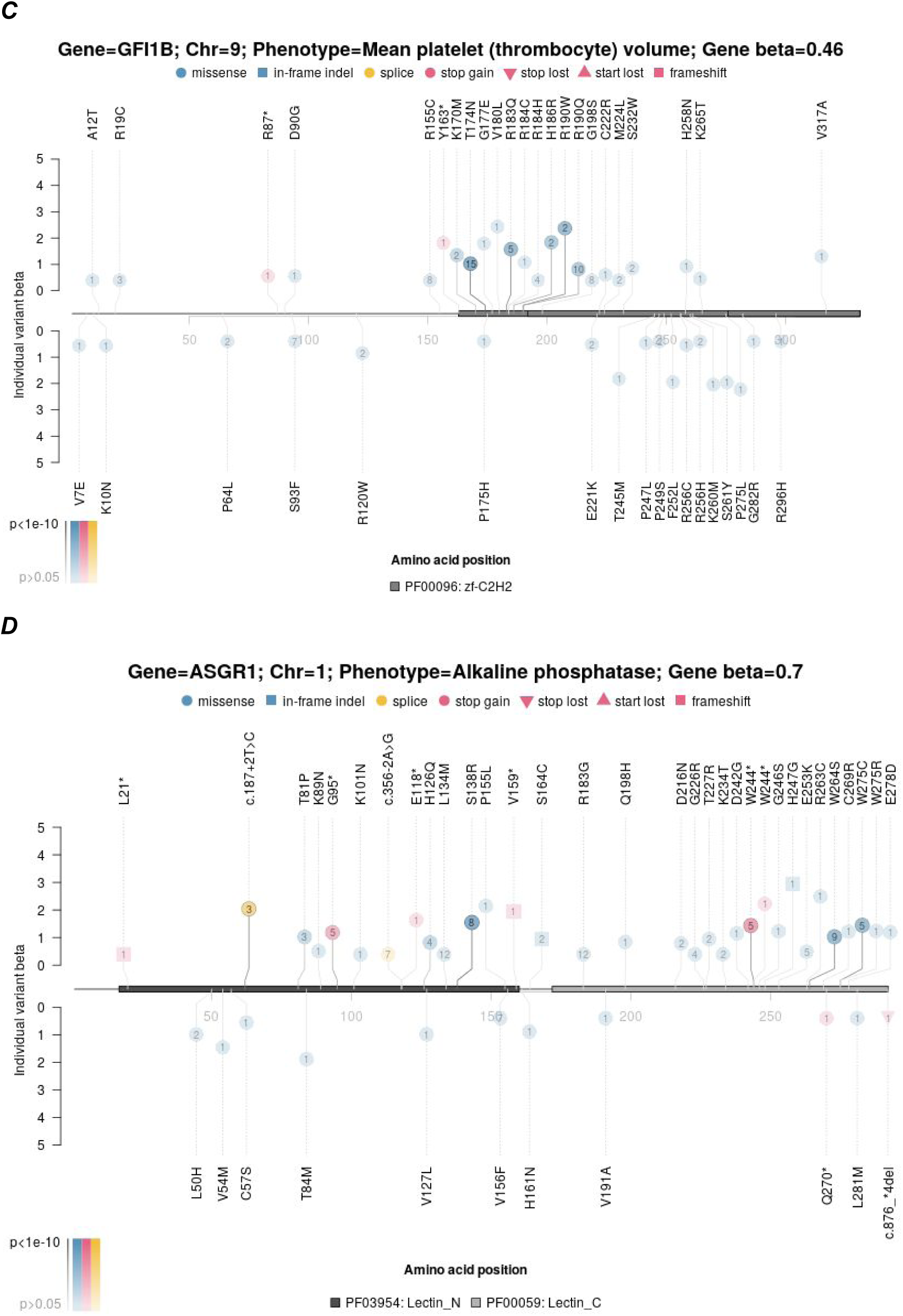
Distribution of effects of rare variants on phenotypes in select genes. A) Plot of the SLC2A9 protein showing the effect sizes of rare damaging variants on urate levels. The legend shows the gene, its associated phenotype, and the Effect Size (β). The effect size is from the gene-based collapsing model, where individuals were coded as either having or not having a qualifying variant. A positive value indicates that variant carriers have, on average, higher values for the phenotype, while a negative value indicates that variant carriers have lower values. The amino acid positions are shown on the x-axis, with the PFAM domain highlighted. The y-axis displays the beta of each individual variant, with negative values shown below and positive values above. Variants are indicated according to their consequence as shown and labeled according to their amino acid change or splice site variation. The number inside the circle is the number of people carrying that variant. Darker lines connecting the variants to the gene and darker-filled shapes indicate more significant p-values for the association. B) Membrane topology plot of SLC2A9 showing variants with positive effect size (green) on urate levels and variants with negative effect size (pink). SLC2A9 (Glut9) reabsorbs urate in the proximal tubules of the kidneys. Variants that disrupt the transmembrane regions or lower gene expression are known to be associated with hypouricemia^18^. Here, 89% of the variants with negative betas, associated with lowered urate levels, are in or directly adjacent to a predicted transmembrane region, as opposed to only 50% of the variants with positive effect size. C) Plot of the GFI1B protein with the effect sizes of rare damaging variants on mean platelet volume shown. Consistent with the literature, variants at the beginning of the zinc finger domain are associated with increased platelet volumes, but we make the novel observation that variants closer to the C terminus may be having an effect in the opposite direction^19,20^. D) Plot of the ASGR1 protein with the effect sizes of rare damaging variants on alkaline phosphatase levels shown. In addition to the known effects of LoF variants, we show that the missense variants that influence alkaline phosphatase levels are more heavily concentrated in the C-type lectin domain (p<0.05 from a Fisher’s exact test comparing the proportion of positively-associated missense variants in this domain to outside of this domain)^21^.

Likewise, variants in different portions of *GFI1B* have distinct effects on mean platelet volume (Figure 3C). Consistent with the literature, variants at the beginning of the zinc finger domain of this gene are associated with increased platelet volumes, but we make the novel observation that variants closer to the C-terminus may be having an effect in the opposite direction^19,20^.

As another example, a significant association is observed between variants in *ASGR1* and alkaline phosphatase levels, with the greatest incidence of mutations that increase alkaline phosphatase levels occurring in the C-type lectin domain, involved in carbohydrate binding. Previously, two loss of function variants in this gene were found to be associated with alkaline phosphatase levels, coronary artery disease, and non-HDL cholesterol^21^. In our analysis, the association is most strongly influenced by LoF variants, but many missense variants in the C-type lectin domain contribute to the signal as well (Figure 3D).

In this way, mapping individual missense variants to their sequence context after a gene-based discovery can refine the classification of missense variants as truly loss of function. Corresponding images detailing the locations for all of the rare variants in each significant gene-phenotype association can be found in the supplement.

### Web resource

We have made the results of our study downloadable via cloud storage (https://s3.amazonaws.com/helix-research-public/ukbb_exome_analysis_results/README.txt) and also browseable with an interactive web tool at https://ukb.research.helix.com.

## Discussion

Here we present the first analysis to catalog the effects of rare and unique coding variants on thousands of phenotypes across two large cohorts. Until now, rare variant analyses using next generation sequence data have been performed on a small number of phenotypes at a time. For example, studies with thousands of sequenced cases have now been undertaken for conditions like schizophrenia, developmental delay, and diabetes ^22–24^. Each of these studies were designed around specific phenotypes and collected targeted, disease-specific samples. Simultaneously analyzing thousands of traits in a biobank population presents additional challenges as well as those inherent to examining rare variants. Due to the rarity of the variants, association tests are prone to false positives. Best practices to produce reliable results include restricting to high quality regions of the genome, setting a very low MAF cutoff, and requiring that at least a minimum threshold of individuals carry qualifying variants in the gene to be analyzed. Now that rare variant information is available to researchers for large numbers of phenotypes and samples, we can expect that new studies will be increasingly successful at utilizing gene-based analyses and other new techniques to characterize the impacts of specific rare variants on different human traits.

Our analysis found that the vast majority of statistically significant gene-based associations were not driven by single explanatory variants. Not only were 95% of the gene-based associations not explained by a single variant, but 78% of the associated genes had no variants with p-values that would pass the multiple testing threshold if analyzed individually.

In genes where multiple rare variants contribute to the signal, we find that mapping the precise contributions of each variant in the context of the secondary and tertiary structures can reveal the most functional parts of the gene for the given phenotype and provide additional support for a statistical association. Formal statistical tests of domain enrichment and discovery analyses that focus on different gene regions will doubtless uncover novel associations but will also often have less power due to the small number of people who will be carrying rare variants in each domain.

Importantly, the associations identified in this analysis can only be obtained using sequencing techniques, as opposed to chip-based methods. All of the variants used in our analysis have a MAF below 0.1%, which is below the range of frequencies that can currently be comfortably imputed. Furthermore, 35% of the 747,865 variants included in our analysis were singletons--only observed once in our dataset and never reported in gnomAD^25^. Such unique variants will never be accessible by chip and were vital to our study’s success. In fact, 87% of our statistically significant associations received worse p-values--on average by threeorders of magnitude--when the singletons were removed from the analysis.

Our analysis differs from the one presented by Van Hout et al., who performed a gene-based analysis on this same UKB dataset and with some of the same phenotypes^7^. Some of our analysis differences included our use of a replication dataset, our more stringent MAF cutoff (1% vs. 0.1%), and collapsing model differences (LoF vs. both LoF and coding models). Our analysis led to the discovery of 84 statistically significant associations, among which we identified 6 of the 17 gene-based associations reported by Van Hout et al. (between *TUBB1, GP1BA, ASXL1, IQGAP2, HBB* and *KLF1* and blood cell phenotypes)^7^. The associations that we did not confirm were largely driven by variants that did not pass the stringent parameters used in our analysis, especially the requirement for MAF<0.1% and at least 10 variant carriers per gene.

Our analysis has a number of limitations. The analysis was restricted to European ancestry individuals. The analysis included rigid criteria for variant qualification and grouped variants at the most basic level, the gene. Future studies can utilize more complex weighting algorithms as opposed to rigid cutoffs and can explore different ways of grouping rare variants, such as by gene family or by exon^26^. Our study used a simple dominant model of inheritance, while recessive models and models that include gene-gene or gene-variant interactions will doubtless provide novel insights as well. Finally, our requirement for at least 10 phenotyped carriers of a qualifying variant in a gene to be included in the analysis, while removing problems with test statistic inflation, also reduced the number of genes that we could investigate. Of the 3,166 phenotypes analyzed, 407 phenotypes resulted in no genes passing this filter.

This analysis presents one of the first forays into a new standard for human genetics research. As the sample sizes of cohorts with extensive phenotypic data and next generation sequencing grows, both through publicly available cohorts such as the UKB or population-based screening efforts such as the Healthy Nevada Project, we are now able to investigate the biological impact of rare variants with the same fine-tuned precision with which we currently assess the effects of common variants. A wealth of discoveries await us as we embark on this next phase of incorporating rare genome sequencing information into truly personalized medicine. We provide an interactive browser of our results as a resource to the human genetics community (https://ukb.research.helix.com/) to facilitate these discoveries.

## Methods

### Samples, phenotypes and variant annotation

We utilized the FE version^27^ of the UKB plink-formatted exome files (field 23160) as well as the imputed genotypes from GWAS genotyping (field 22801-22823). The HNP study was reviewed and approved by the University of Nevada, Reno Institutional Review Board (IRB, project 956068-12). The HNP samples were sequenced at Helix using the Exome+ assay, a proprietary exome that combines a highly performant medical exome with a microarray-equivalent SNP backbone into a single sequencing assay (www.helix.com)^28^. Data were processed using a custom version of Sentieon and aligned to GRCh38, with variant calling and phasing algorithms following GATK best practices^29^. Imputation of common variants in the HNP data was performed by pre-phasing samples and then imputing. Pre-phasing was performed using reference databases, which include the 1000 Genomes Phase 3 data. This was followed by genotype imputation for all 1000 Genomes Phase 3 sites that have genotype quality (GQ) values less than 20. Imputation results were then filtered for quality so that only high precision imputed variant calls were reported.

Variant annotation was performed with VEP^30^. Coding regions were defined according to Gencode version GENCODE 26, and the Ensembl canonical transcript was used to determine variant consequence^31,32^. Genotype processing was performed in Hail^33^.

For the collapsing analysis, samples were coded as a 1 for each gene if they had a qualifying variant and a 0 otherwise. We defined “qualifying” as coding (stop_lost, missense_variant, start_lost, splice_donor_variant, inframe_deletion, frameshift_variant, splice_acceptor_variant, stop_gained, or inframe_insertion) and not Polyphen or SIFT benign (Polyphen benign is <0.15, SIFT benign is >0.05)^34,35^. We also ran a loss of function (LoF) model that only included LoF variants (stop_lost, start_lost, splice_donor_variant, frameshift_variant, splice_acceptor_variant, or stop_gained). We used a MAF cutoff of 0.1%. To pass the MAF filter, the variant must be below that frequency cutoff in all gnomAD populations^25^ as well as the European ancestry UKB exomes (defined by UKB as Genetic Ethnicity = Caucasian in field 22006). This was the sample set to which we restricted in our analysis. Only PASS calls were used in the analysis, with an average depth of 62.9x. Qualifying variants were also restricted to the high-confidence regions of the genome as defined by the Genome in a Bottle resource for NA12878^36^.

Most UKB phenotypes were processed using the Neale lab modified version of PHESANT, which transforms quantitative traits to normally distributed data and breaks up categorical traits into binary sets^37,38^. ICD-10 diagnosis code phenotypes were coded with 1 if participants had the ICD-10 code recorded at least once in their series of Electronic Health Records (EHR), and otherwise a 0, with controls restricted to one sex when appropriate. HNP phenotypes were processed in the same fashion, with the additional step that the pre-transformed median of quantitative traits was taken when multiple measurements were available.

### Analysis

We used BOLT-LMM for statistical analysis^39^. Briefly, this method builds a linear mixed model using common variants to account for the effects of relatedness and population stratification. The covariates included were age and sex. In the HNP analysis, the Helix bioinformatics pipeline version was also included as a covariate in the model to account for batch effects.

A representative set of LD-pruned, high-quality common variants were identified for both the creation of principal components and for the random effects and trait heritability in the BOLT-LMM mixed model. Inclusion in this set required MAF>1%, imputation with reasonable accuracy in UKB (INFO>0.7) and high coverage or imputation in Helix samples (>99% of samples with a sequence-based call or an imputed call with GP>0.95), LD-pruned (r^2^<0.6) to a set of 184,445 variants. This set of variants was genome-wide, including both coding and noncoding regions. The set of unrelated European ancestry individuals from UKB was used for LD pruning and MAF cutoffs. In the interest of saving time and compute power, the random effects in the BOLT-LMM mixed model for individual variants, as opposed to the faster gene-based analyses, were further restricted to a set of 13,036 LD-pruned variants (r^2^<0.01).

Our gene-based analyses required at least 10 carriers of qualifying variants in analyzed genes for quantitative traits and at least 10 carriers of qualifying variants to be expected in the smaller sample group for analyzed genes for each binary trait, similar to previously suggested guidelines^13^.

Meta-analysis was performed on the summary stats from each separate analysis using *PLINK*, and we required statistically significant hits to have at least one variant carrier from both the UKB and the HNP groups in addition to 10 variant carriers overall^40,41^.

BOLT-LMM determined that 298 phenotypes had 0 heritability based on the 184,445 common variants. These phenotypes were analyzed by logistic regression of unrelated individuals using *PLINK* 2.0 with age, sex, and the first 10 European-specific principal components (calculated on these 184,445 variants) included as covariates^40,42^.

Gene plots were made using trackViewer and Protter and annotated with Pfam domains v. 32.0^43–45^.

## Supporting information

Gene plots for coding model hits

Gene plots for LoF model hits

## Acknowledgements

This research has been conducted using the UK Biobank Resource under Application Number 40436. We acknowledge A. Buckley and J. Ou for assistance with figures, O. Mendelevitch, G. Sayfan, and T. Michaud for assistance with creating the web resource, and to W. Lee and the entire Helix Bioinformatics team for their contributions to the production exome sequencing pipeline. We thank M. Henderson, T. Curreri and all the ambassadors of the Healthy Nevada Project (HNP). We thank Renown Health and DRI marketing for helping to launch the HNP project.

## References

1. Richardson, T. G., Harrison, S., Hemani, G. & Davey Smith, G. An atlas of polygenic risk score associations to highlight putative causal relationships across the human phenome. Elife 8, (2019).

2. Khera, A. V. et al. Genome-wide polygenic scores for common diseases identify individuals with risk equivalent to monogenic mutations. Nat. Genet. 50, 1219–1224 (2018).

3. Krapohl, E. et al. Phenome-wide analysis of genome-wide polygenic scores. Mol. Psychiatry 21, 1188–1193 (2016).

4. Long, T. et al. Whole-genome sequencing identifies common-to-rare variants associated with human blood metabolites. Nat. Genet. 49, 568–578 (2017).

5. Zhu, Q. et al. A genome-wide comparison of the functional properties of rare and common genetic variants in humans. Am. J. Hum. Genet. 88, 458–468 (2011).

6. Nejentsev, S., Walker, N., Riches, D., Egholm, M. & Todd, J. A. Rare variants of IFIH1, a gene implicated in antiviral responses, protect against type 1 diabetes. Science 324, 387–389 (2009).

7. Van Hout, C. V. et al. Whole exome sequencing and characterization of coding variation in 49,960 individuals in the UK Biobank. bioRxiv 572347 (2019). doi:10.1101/572347

8. Li, B. & Leal, S. M. Methods for detecting associations with rare variants for common diseases: application to analysis of sequence data. Am. J. Hum. Genet. 83, 311–321 (2008).

9. Lee, S., Abecasis, G. R., Boehnke, M. & Lin, X. Rare-Variant Association Analysis: Study Designs and Statistical Tests. The American Journal of Human Genetics 95, 5–23 (2014).

10. Do, R. et al. Exome sequencing identifies rare LDLR and APOA5 alleles conferring risk for myocardial infarction. Nature 518, 102–106 (2015).

11. Cirulli, E. T. et al. Exome sequencing in amyotrophic lateral sclerosis identifies risk genes and pathways. Science 347, 1436–1441 (2015).

12. Liu, C. et al. Meta-analysis identifies common and rare variants influencing blood pressure and overlapping with metabolic trait loci. Nat. Genet. 48, 1162–1170 (2016).

13. Churchhouse, C. Details and considerations of the UK Biobank GWAS. Neale lab (2017). Available at:http://www.nealelab.is/blog/2017/9/11/details-and-considerations-of-the-uk-biobank-gwas. (Accessed: 19th April 2019)

14. Wylie, L. A., Mouillesseaux, K. P., Chong, D. C. & Bautch, V. L. Developmental SMAD6 loss leads to blood vessel hemorrhage and disrupted endothelial cell junctions. Dev. Biol. 442, 199–209 (2018).

15. Kenny, E. E. et al. Melanesian blond hair is caused by an amino acid change in TYRP1. Science 336, 554 (2012).

16. Corbyn, Z. Blonde hair evolved more than once. Nature (2012). doi:10.1038/nature.2012.10587

17. Marouli, E. et al. Rare and low-frequency coding variants alter human adult height. Nature 542, 186–190 (2017).

18. Ruiz, A., Gautschi, I., Schild, L. & Bonny, O. Human Mutations in SLC2A9 (Glut9) Affect Transport Capacity for Urate. Front. Physiol. 9, 476 (2018).

19. Möröy, T., Vassen, L., Wilkes, B. & Khandanpour, C. From cytopenia to leukemia: the role of Gfi1 and Gfi1b in blood formation. Blood 126, 2561–2569 (2015).

20. Polfus, L. M. et al. Whole-Exome Sequencing Identifies Loci Associated with Blood Cell Traits and Reveals a Role for Alternative GFI1B Splice Variants in Human Hematopoiesis. Am. J. Hum. Genet. 99, 785 (2016).

21. Nioi, P. et al. Variant ASGR1 Associated with a Reduced Risk of Coronary Artery Disease. N. Engl. J. Med. 374, 2131–2141 (2016).

22. Genovese, G. et al. Increased burden of ultra-rare protein-altering variants among 4,877 individuals with schizophrenia. Nat. Neurosci. 19, 1433–1441 (2016).

23. Deciphering Developmental Disorders Study. Large-scale discovery of novel genetic causes of developmental disorders. Nature 519, 223–228 (2015).

24. Fuchsberger, C. et al. The genetic architecture of type 2 diabetes. Nature 536, 41–47 (2016).

25. Lek, M. et al. Analysis of protein-coding genetic variation in 60,706 humans. Nature 536, 285–291 (2016).

26. Wu, M. C. et al. Rare-variant association testing for sequencing data with the sequence kernel association test. Am. J. Hum. Genet. 89, 82–93 (2011).

27. Regier, A. A. et al. Functional equivalence of genome sequencing analysis pipelines enables harmonized variant calling across human genetics projects. Nat. Commun. 9, 4038 (2018).

28. Helix’s Exome+ Performance White Paper. Available at: https://cdn.helix.com/wp-content/uploads/2017/07/Helix-PersonalGenomics-Platform-White-Paper.pdf.

29. Kendig, K. I. et al. Computational performance and accuracy of Sentieon DNASeq variant calling workflow. bioRxiv 396325 (2018). doi:10.1101/396325

30. McLaren, W. et al. The Ensembl Variant Effect Predictor. Genome Biol. 17, 122 (2016).

31. Frankish, A. et al. GENCODE reference annotation for the human and mouse genomes. Nucleic Acids Res. 47, D766–D773 (2019).

32. Zerbino, D. R. et al. Ensembl 2018. Nucleic Acids Res. 46, D754–D761 (2018).

33. hail-is. hail-is/hail. GitHub Available at: https://github.com/hail-is/hail. (Accessed: 19th April 2019)

34. Adzhubei, I. A. et al. A method and server for predicting damaging missense mutations. Nat. Methods 7, 248–249 (2010).

35. Sim, N.-L. et al. SIFT web server: predicting effects of amino acid substitutions on proteins. Nucleic Acids Res. 40, W452–7 (2012).

36. Genome in a Bottle. Available at: ftp://ftp-trace.ncbi.nlm.nih.gov/giab/ftp/release/NA12878_HG001/NISTv3.3.2/GRCh38/.

37. astheeggeggs. astheeggeggs/PHESANT. GitHub Available at: https://github.com/astheeggeggs/PHESANT. (Accessed: 19th April 2019)

38. Millard, L., Davies, N. M., Gaunt, T., Smith, G. D. & Tilling, K. PHESANT: a tool for performing automated phenome scans in UK Biobank. doi:10.1101/111500

39. Loh, P.-R., Kichaev, G., Gazal, S., Schoech, A. P. & Price, A. L. Mixed model association for biobank-scale data sets. doi:10.1101/194944

40. Purcell, S. et al. PLINK: A Tool Set for Whole-Genome Association and Population-Based Linkage Analyses. The American Journal of Human Genetics 81, 559–575 (2007).

41. Purcell, S. plinkv1.07. Available at: http://zzz.bwh.harvard.edu/plink/.

42. Yang, J., Zaitlen, N. A., Goddard, M. E., Visscher, P. M. & Price, A. L. Advantages and pitfalls in the application of mixed-model association methods. Nat. Genet. 46, 100–106 (2014).

43. Omasits, U., Ahrens, C. H., Müller, S. & Wollscheid, B. Protter: interactive protein feature visualization and integration with experimental proteomic data. Bioinformatics 30, 884–886 (2014).

44. Ou, J. & Zhu, L. J. trackViewer: a Bioconductor package for interactive and integrative visualization of multi-omics data. Nat. Methods 16, 453–454 (2019).

45. El-Gebali, S. et al. The Pfam protein families database in 2019. Nucleic Acids Res. 47, D427–D432 (2019).

