## Supplementary figures and images for "Genome-wide rare variant analysis for thousands of phenotypes in 54,000 exomes"

### ABCA1_x30630.pdf

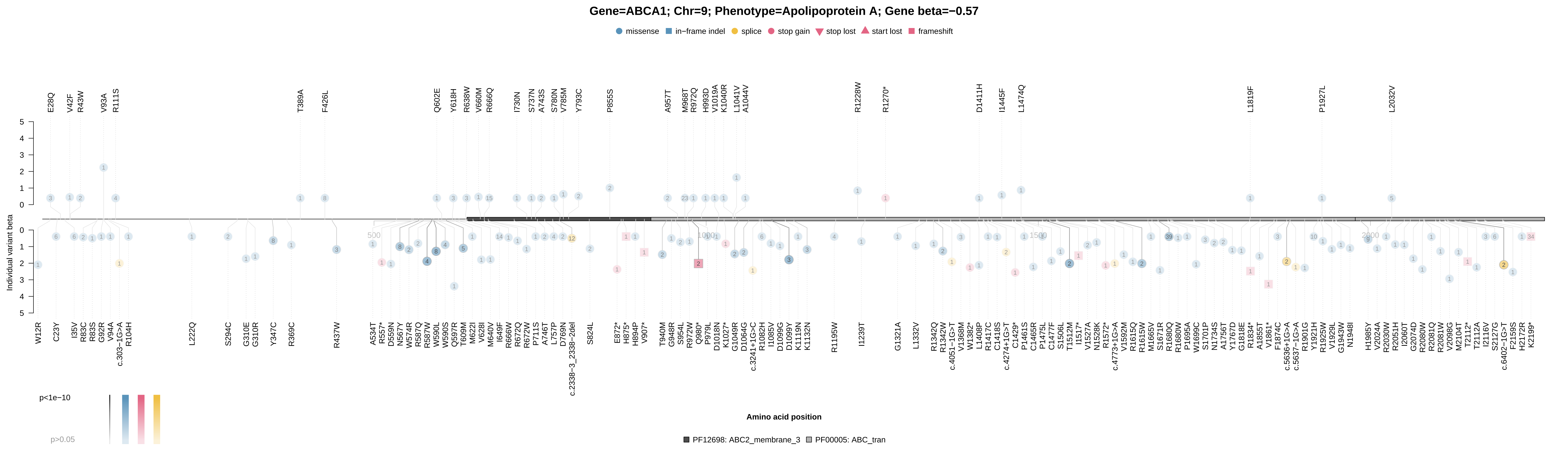

### ABCA1_x30630.pdf

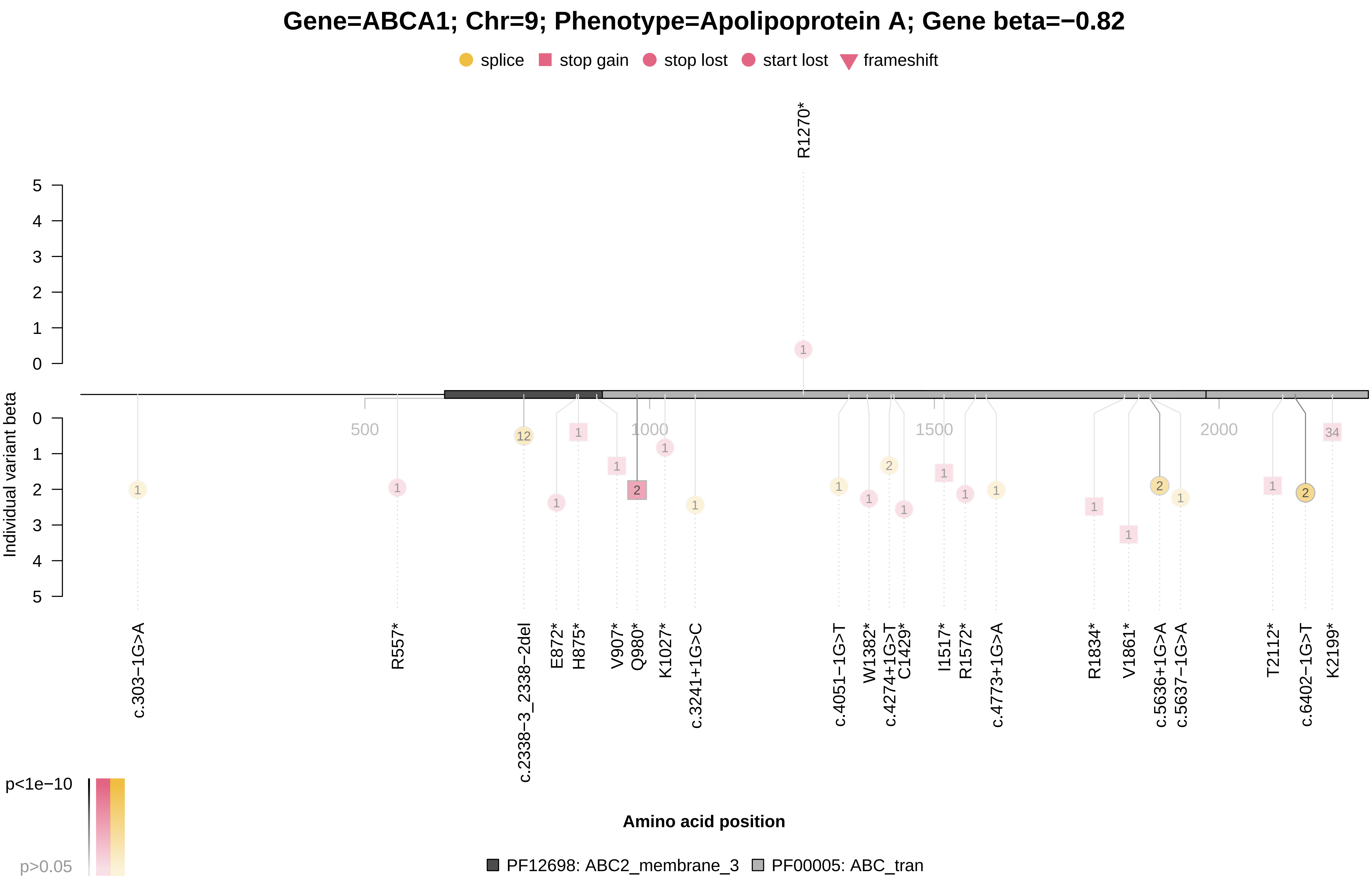

### ABCA1_x30760.pdf

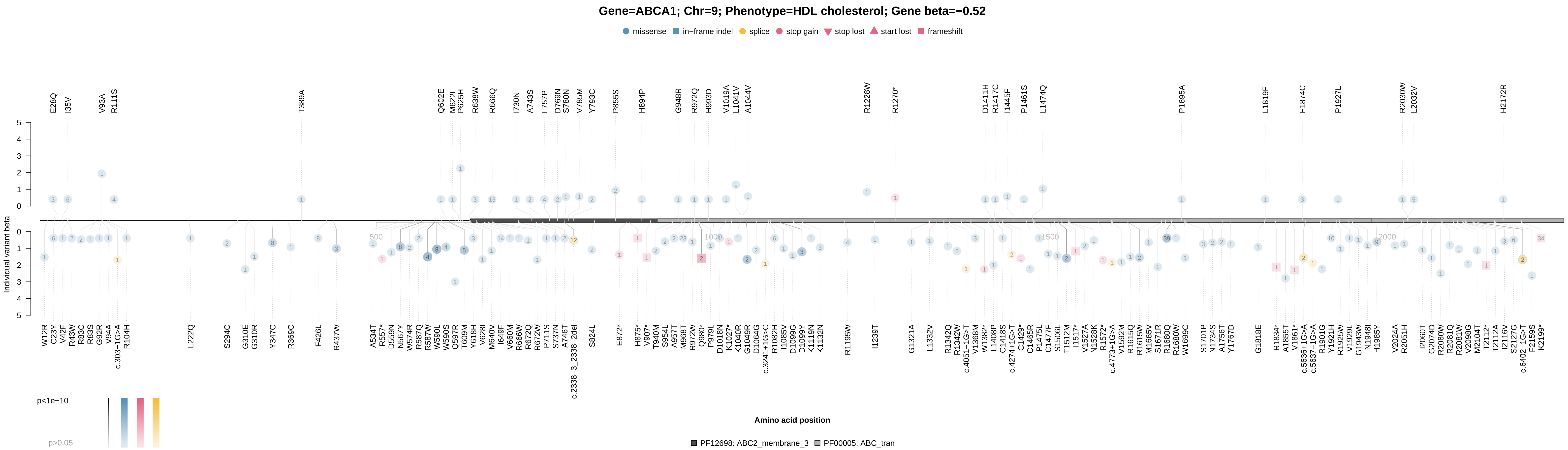

### ABCA1_x30760.pdf

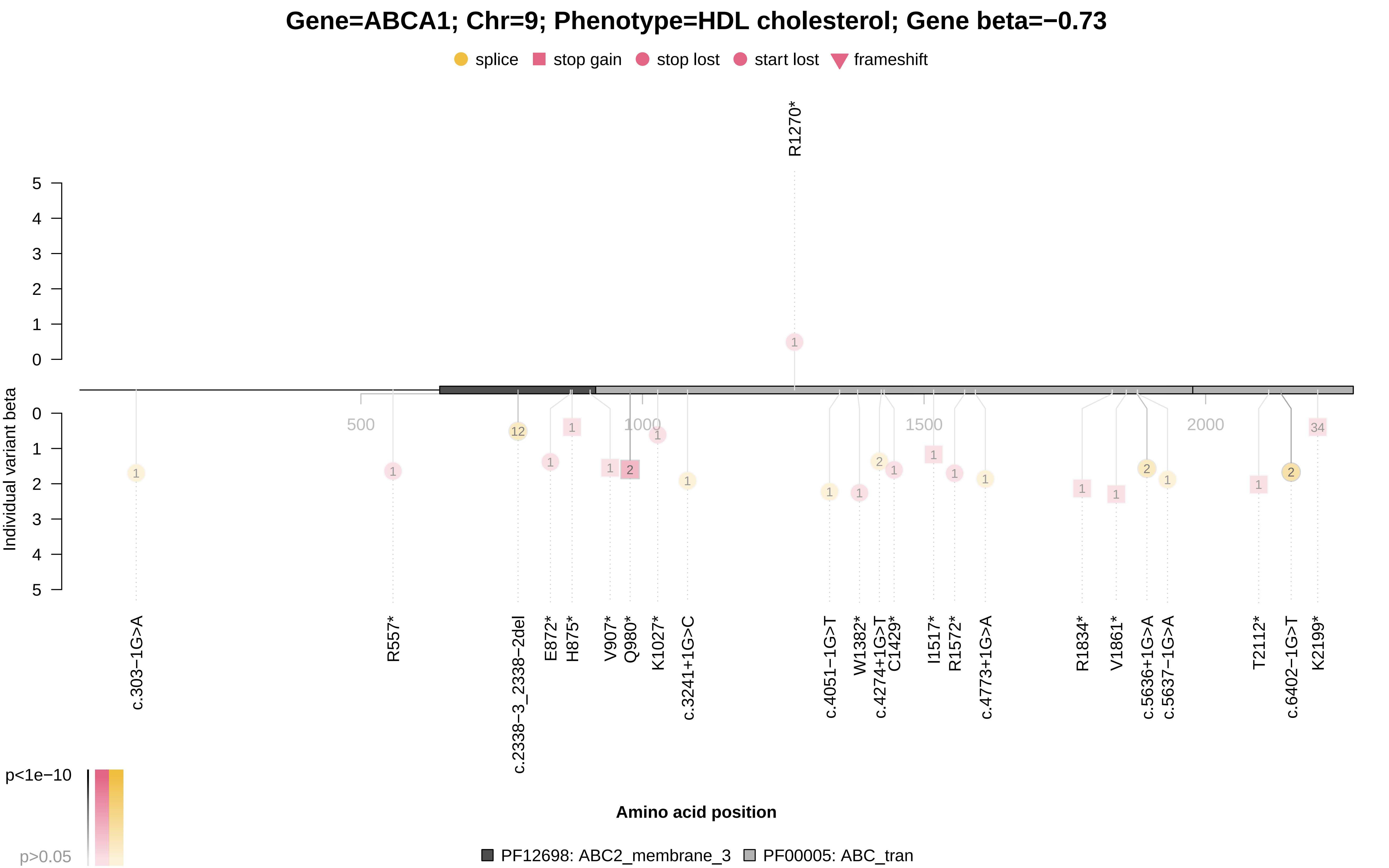

### ABCB11_x30610.pdf

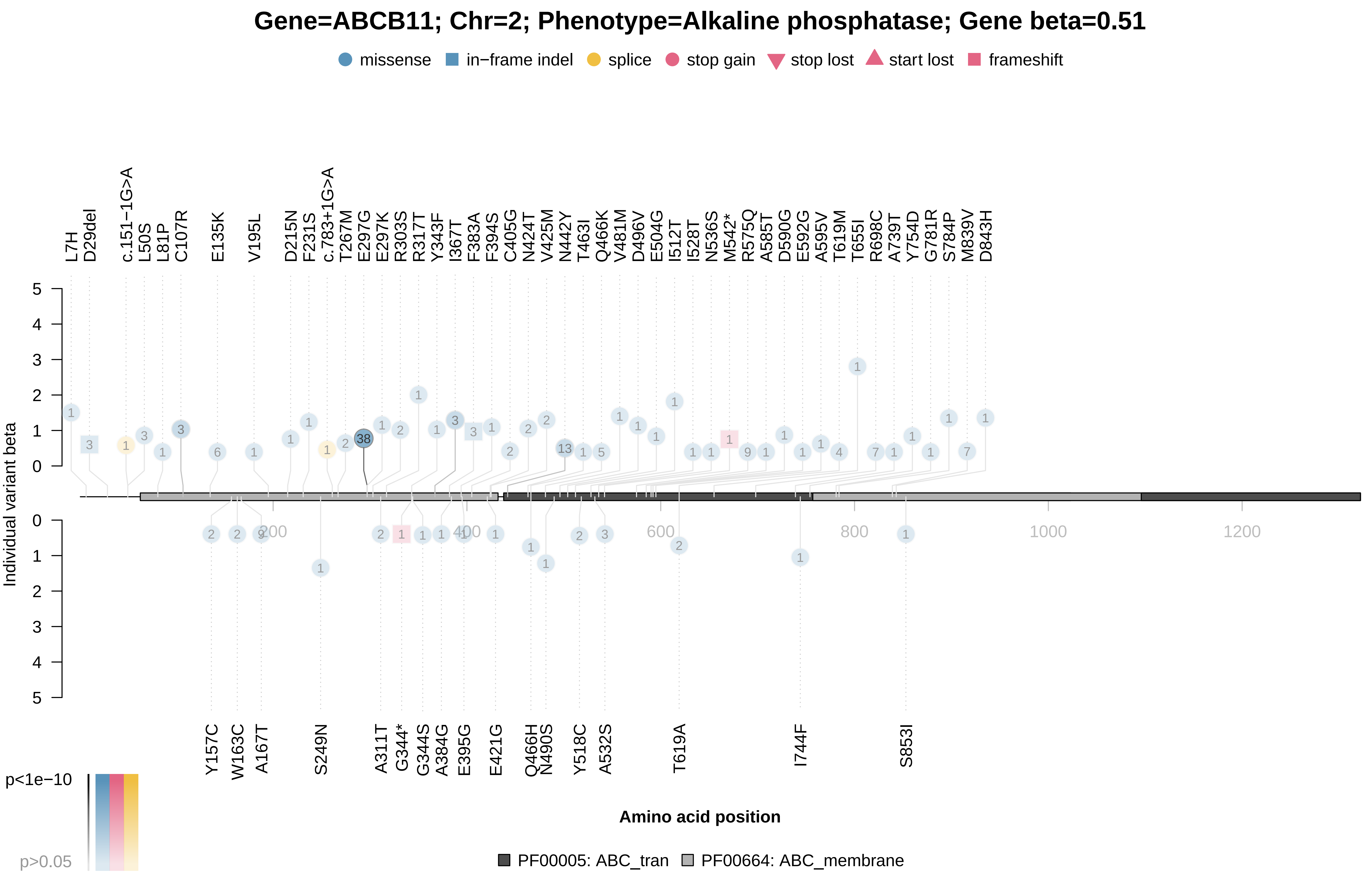

### ALB_x30600.pdf

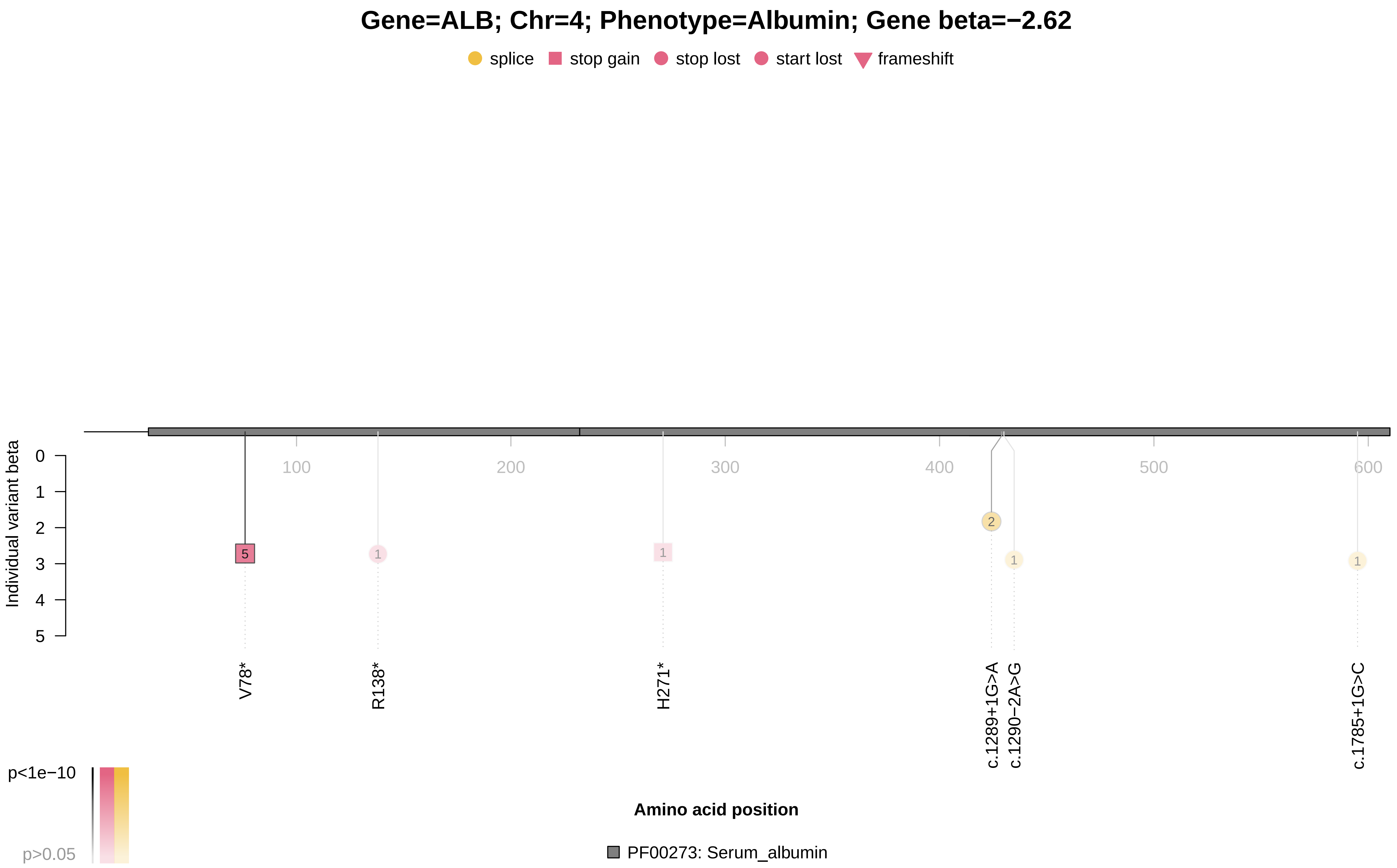

### ALPL_x30610.pdf

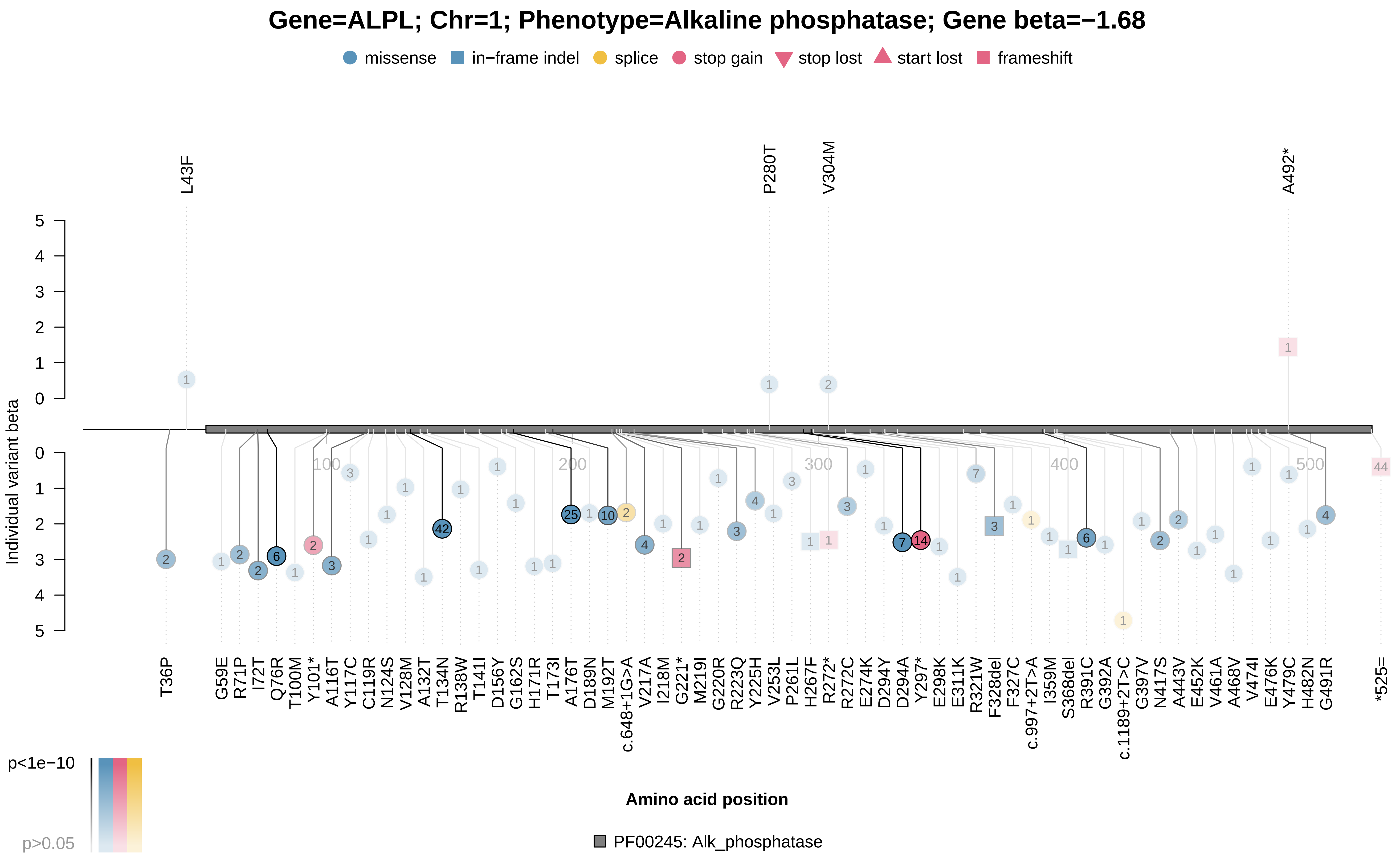

### ALPL_x30610.pdf

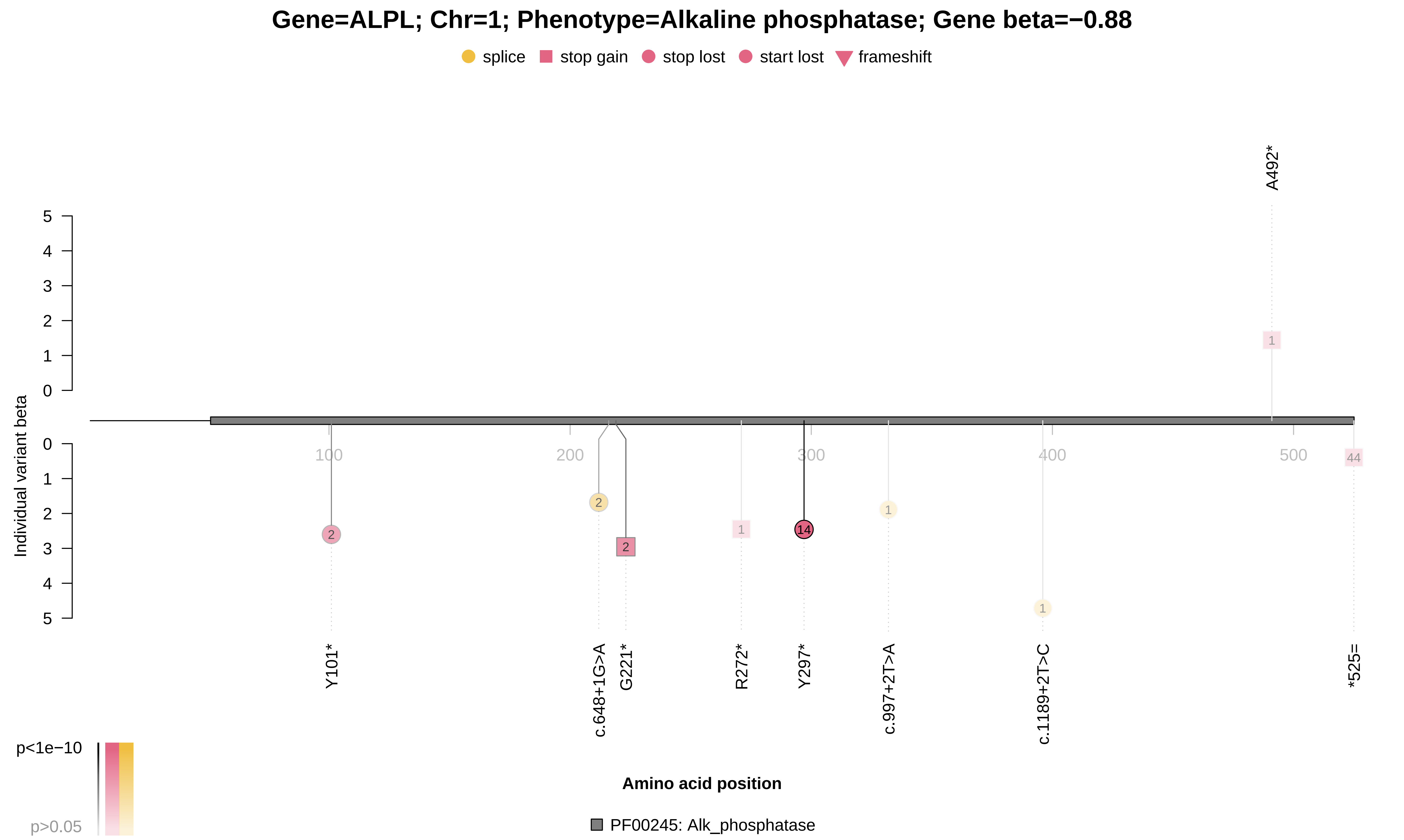

### ALPL_x30810.pdf

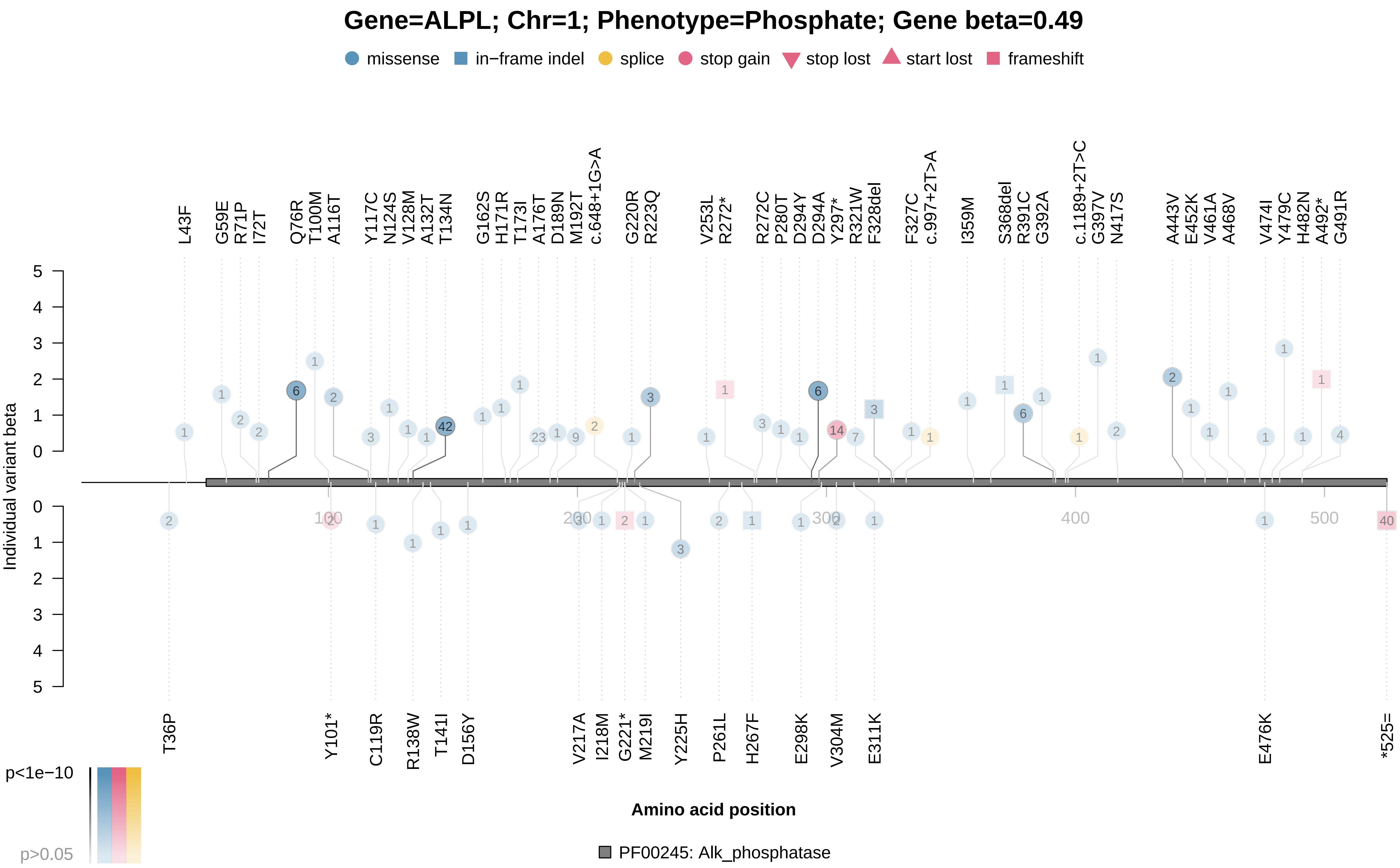

### ANGPTL3_x30630.pdf

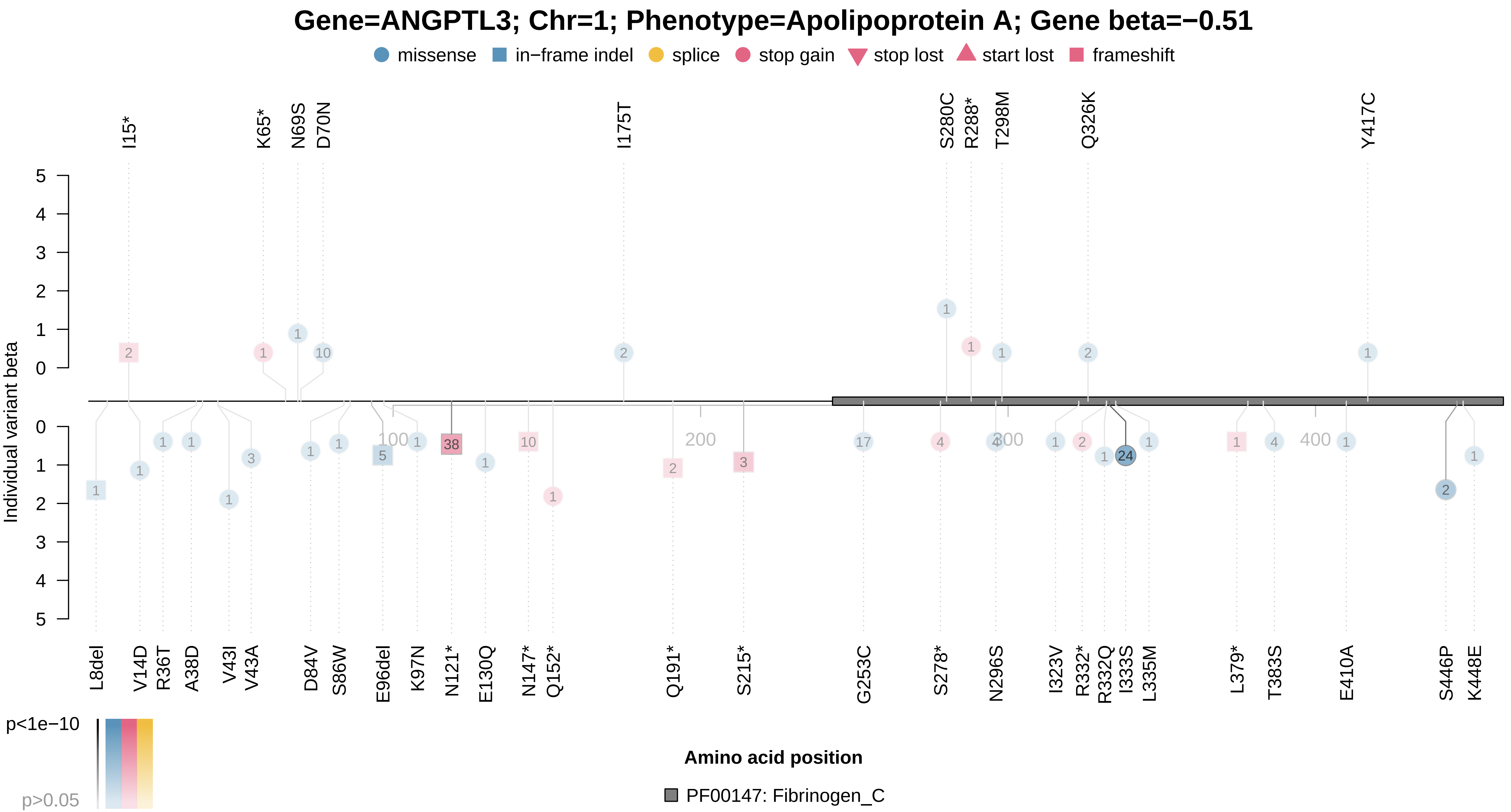

### ANGPTL3_x30870.pdf

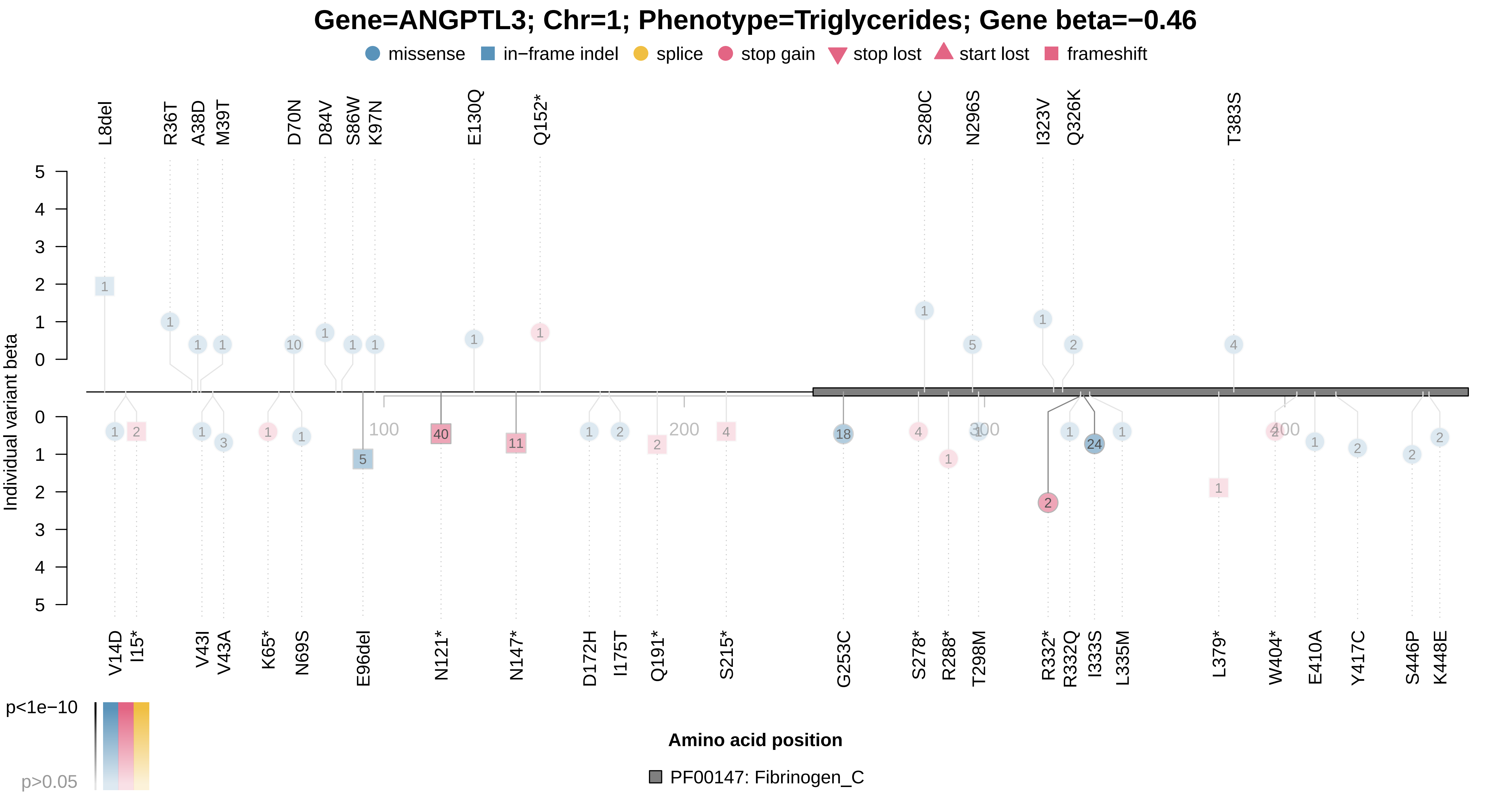

### APOA5_x30870.pdf

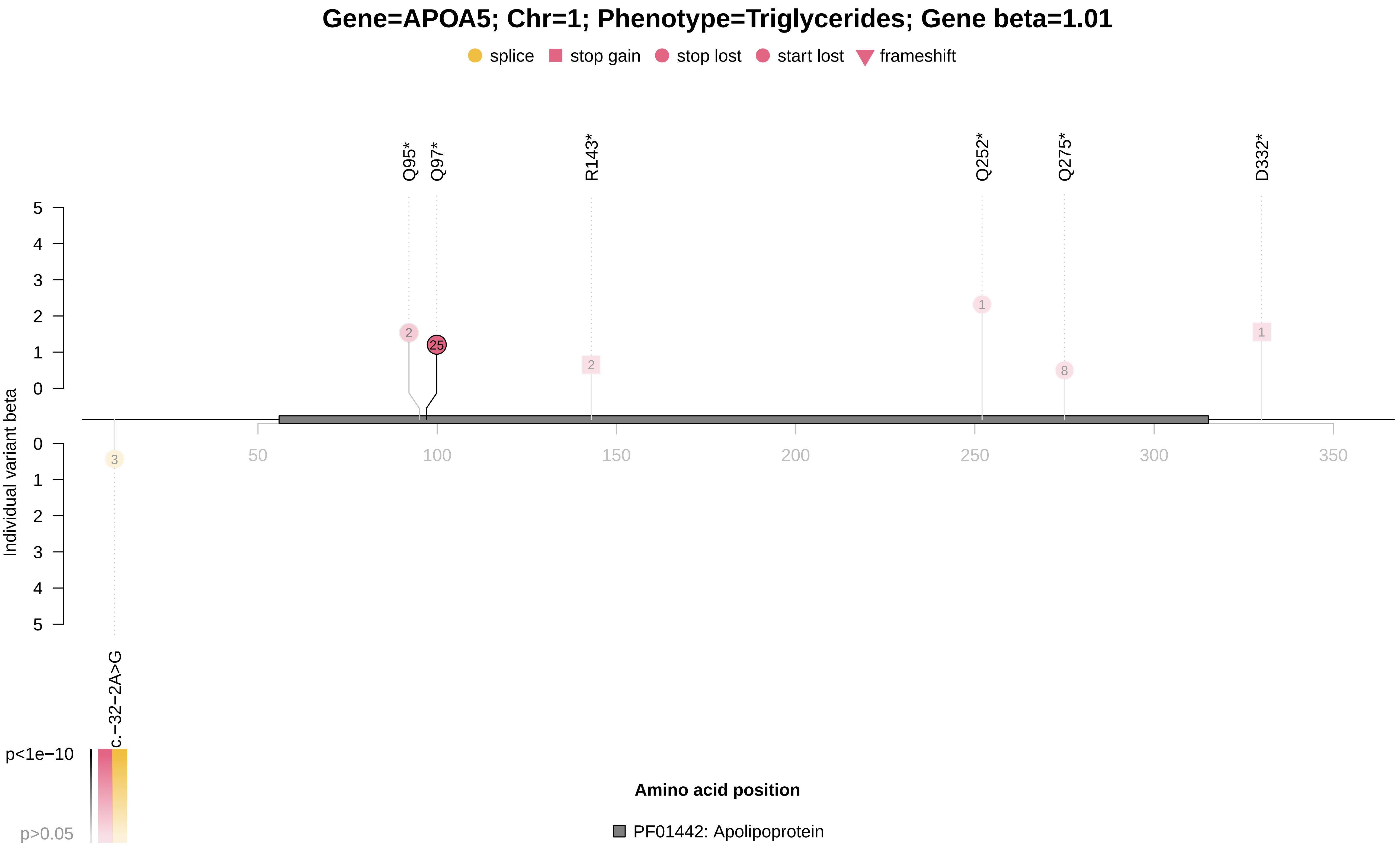

### APOB_x30640.pdf

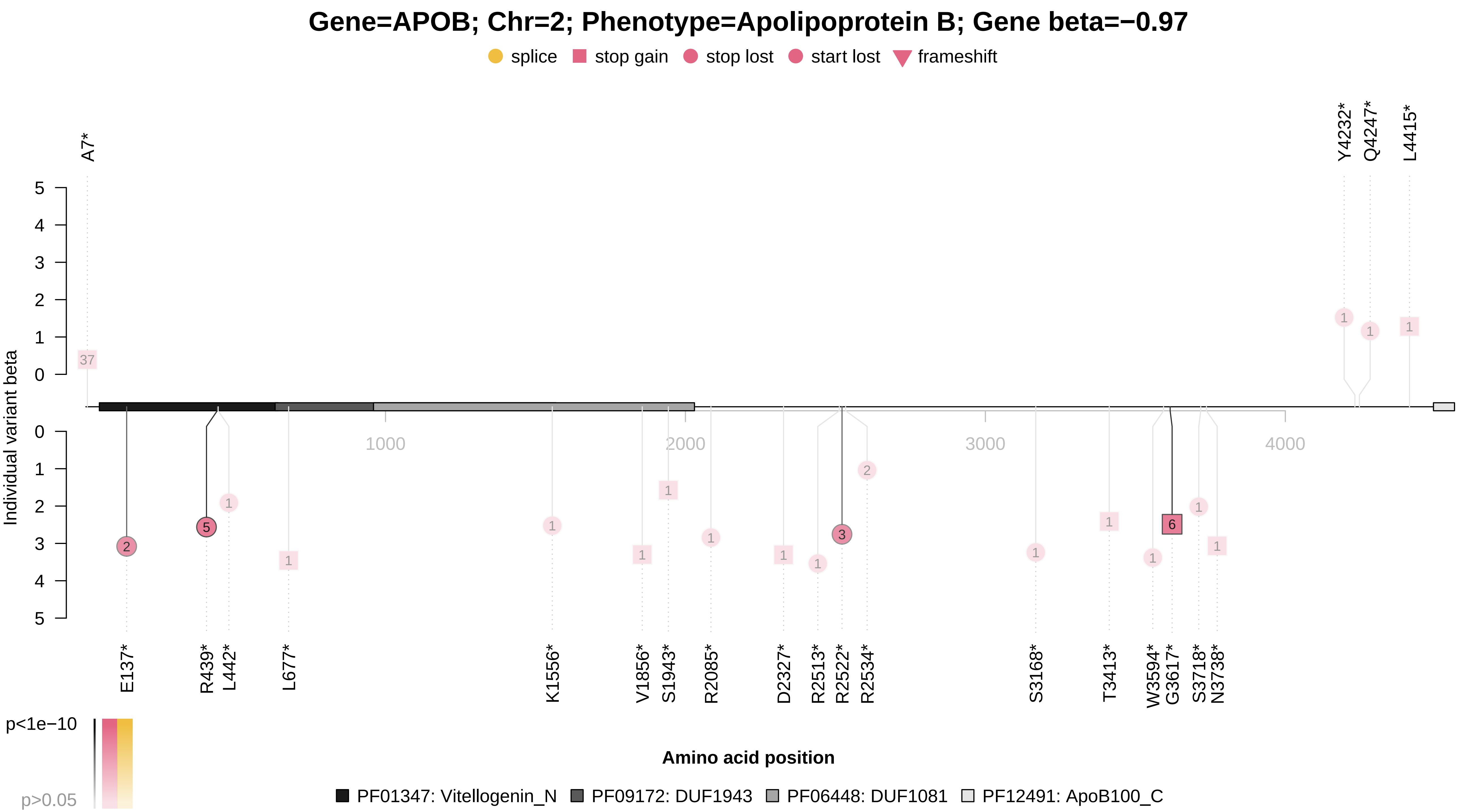

### APOB_x30690.pdf

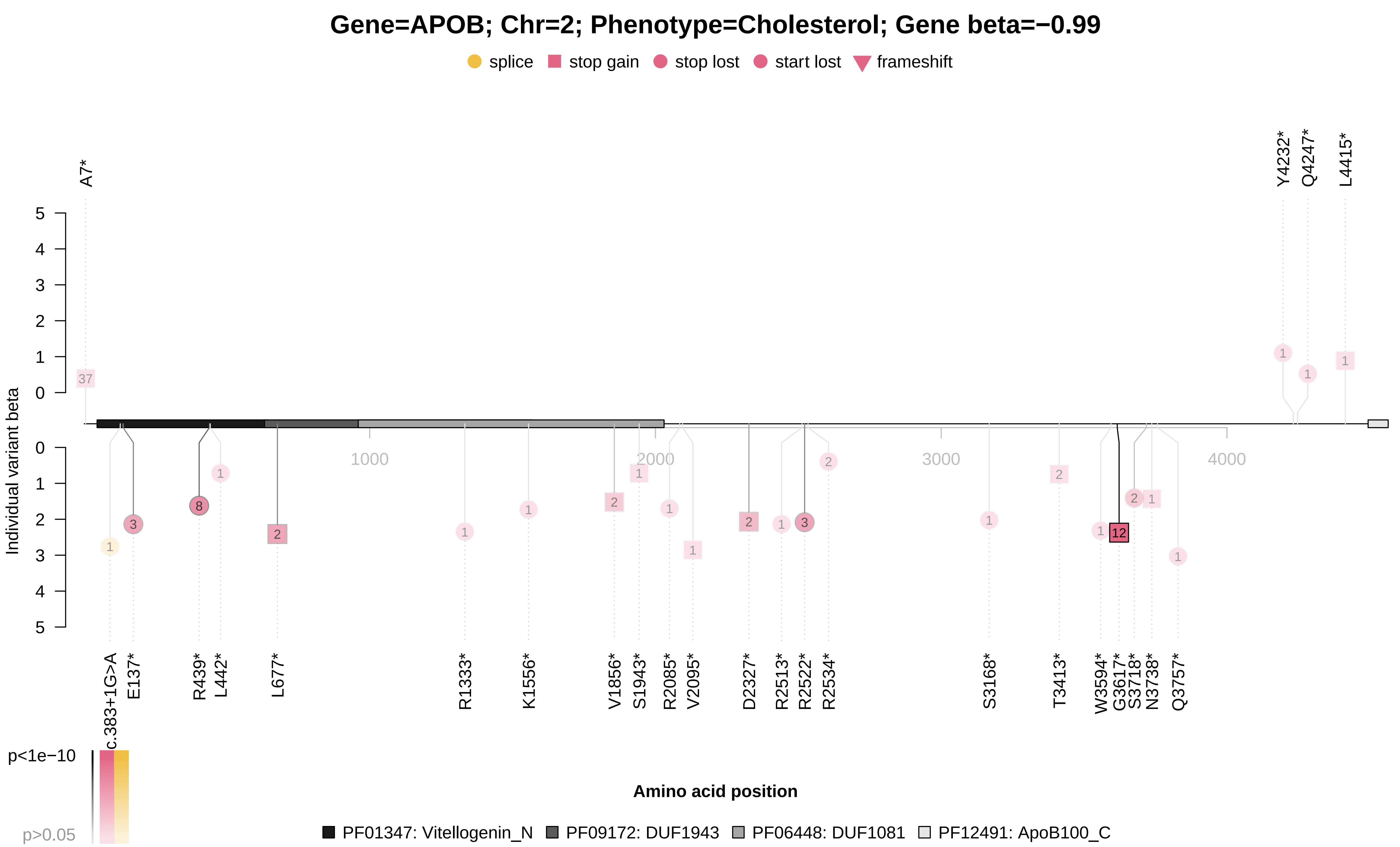

### APOB_x30780.pdf

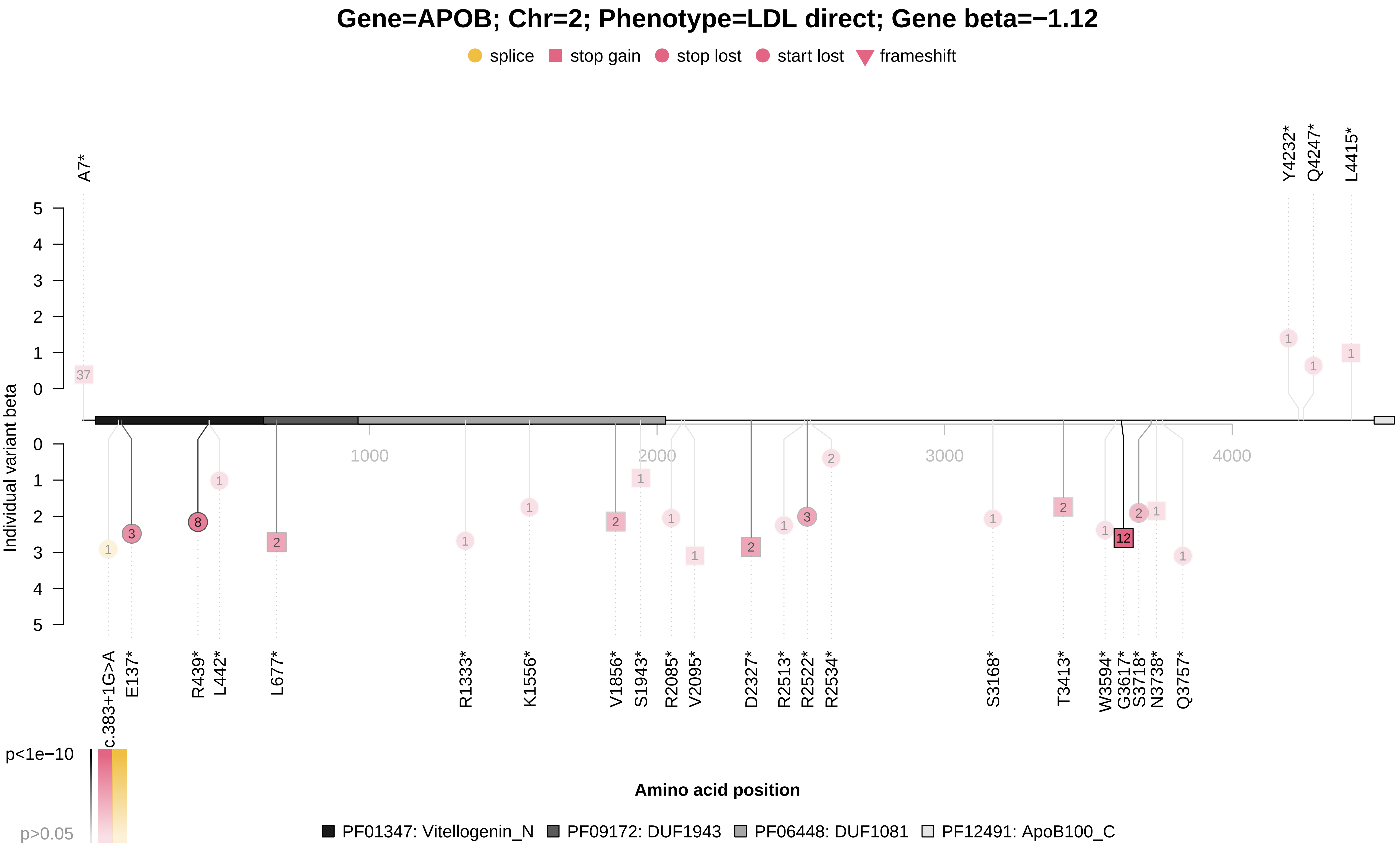

### APOB_x30870.pdf

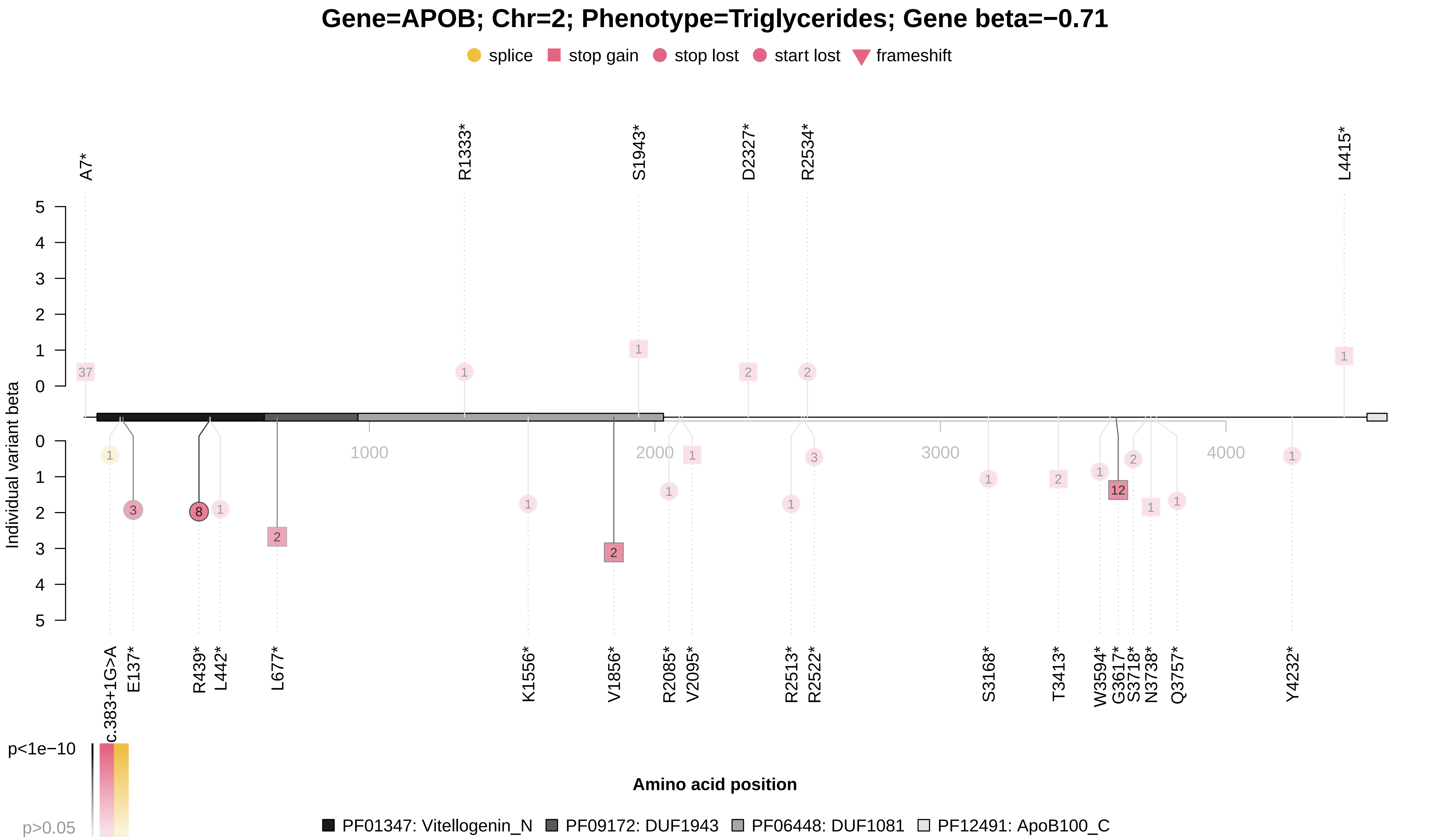

### ASGR1_x30610.pdf

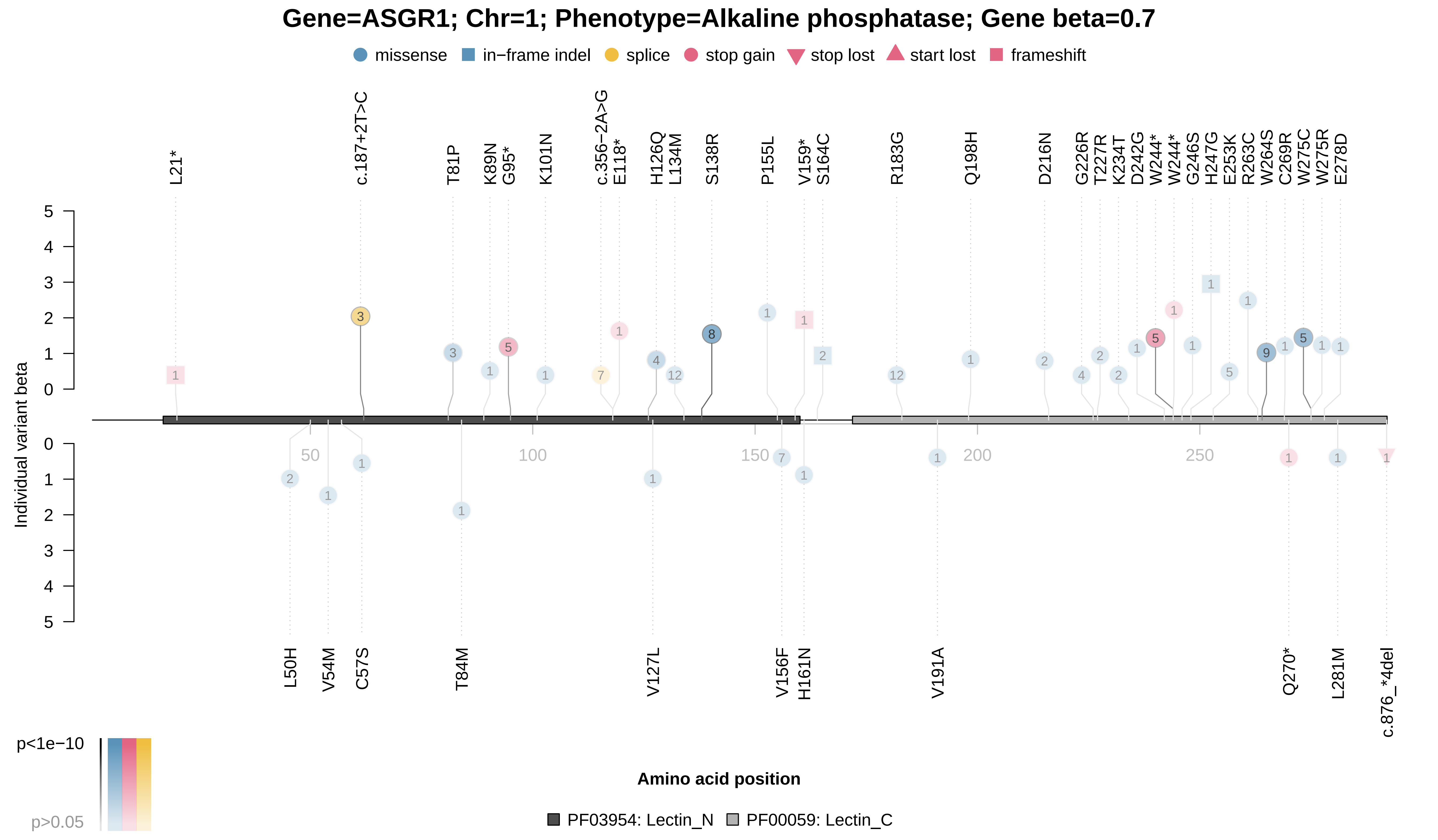

### ASGR1_x30610.pdf

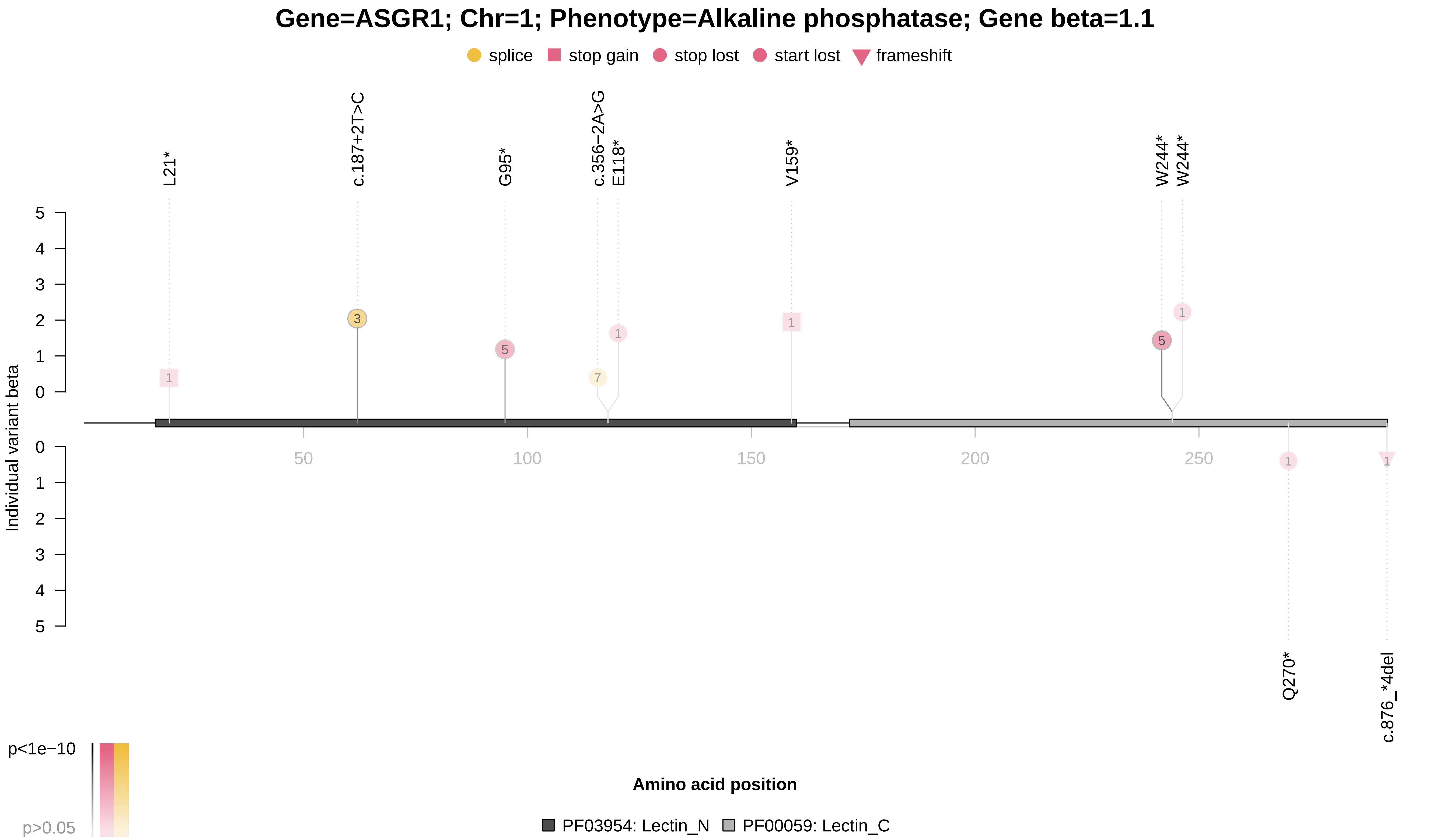

### CETP_x30760.pdf

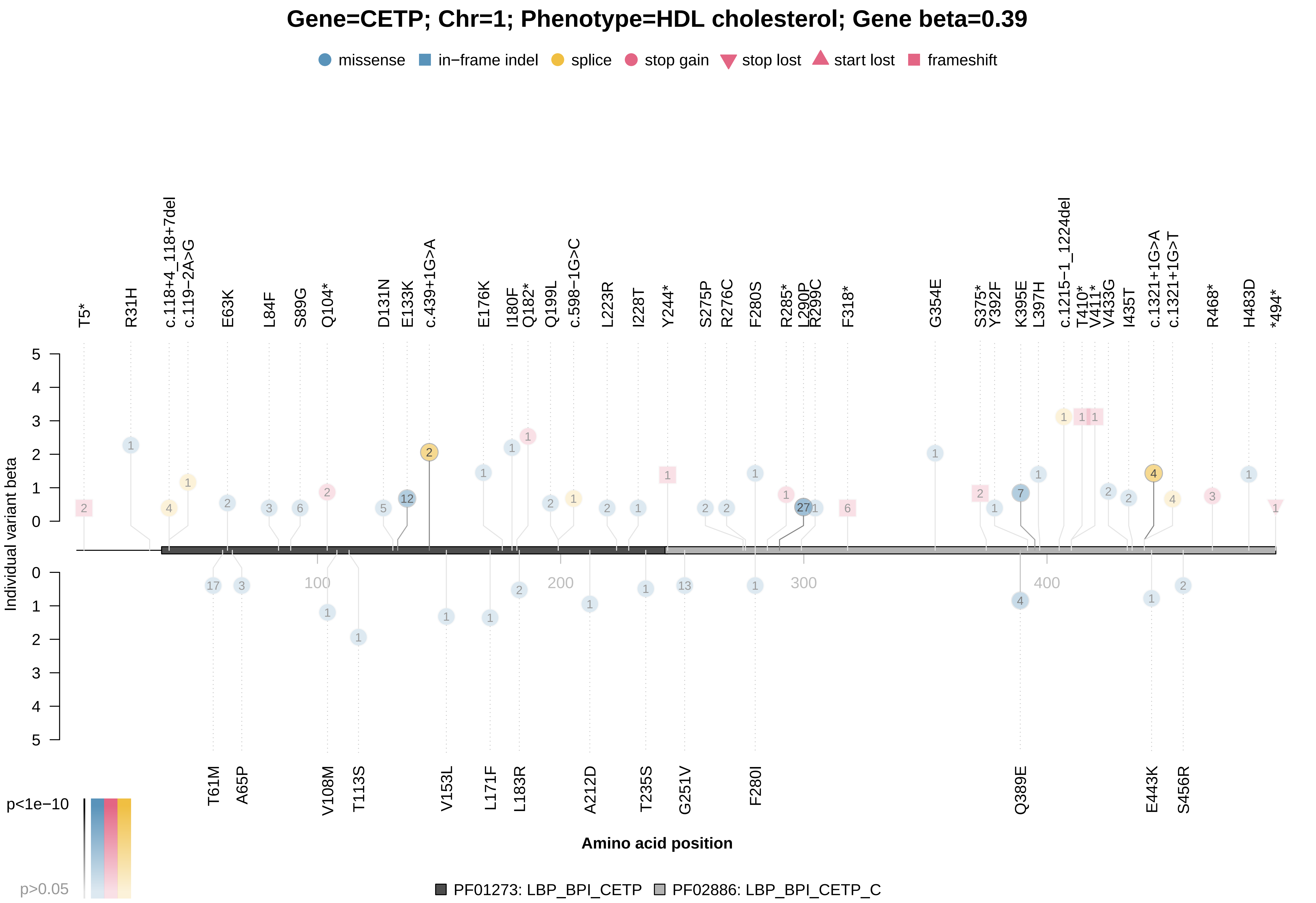

### CETP_x30760.pdf

#### Gene=CETP; Chr=1; Phenotype=HDL cholesterol; Gene beta=0.93

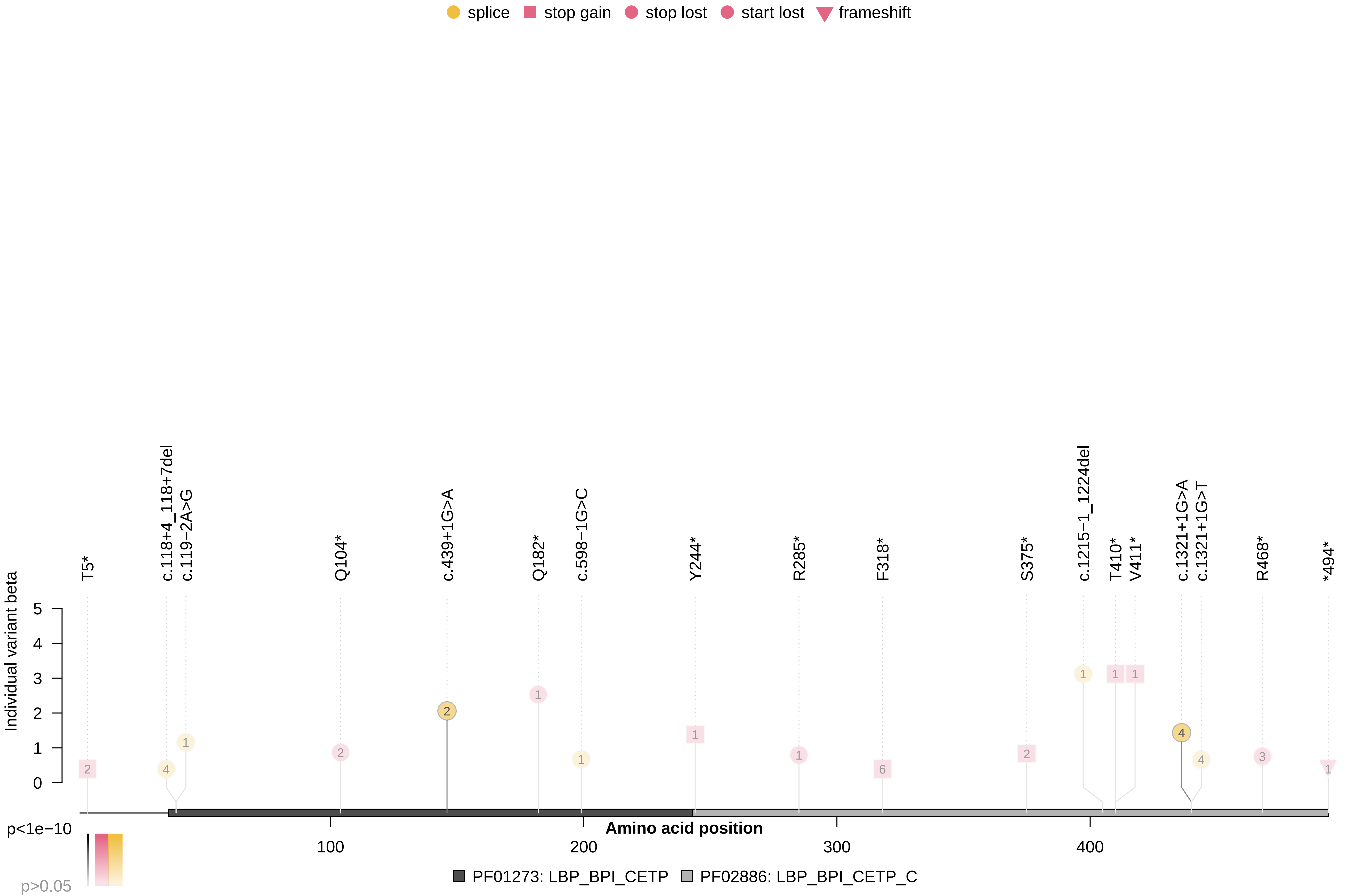

### COL4A4_R31.pdf

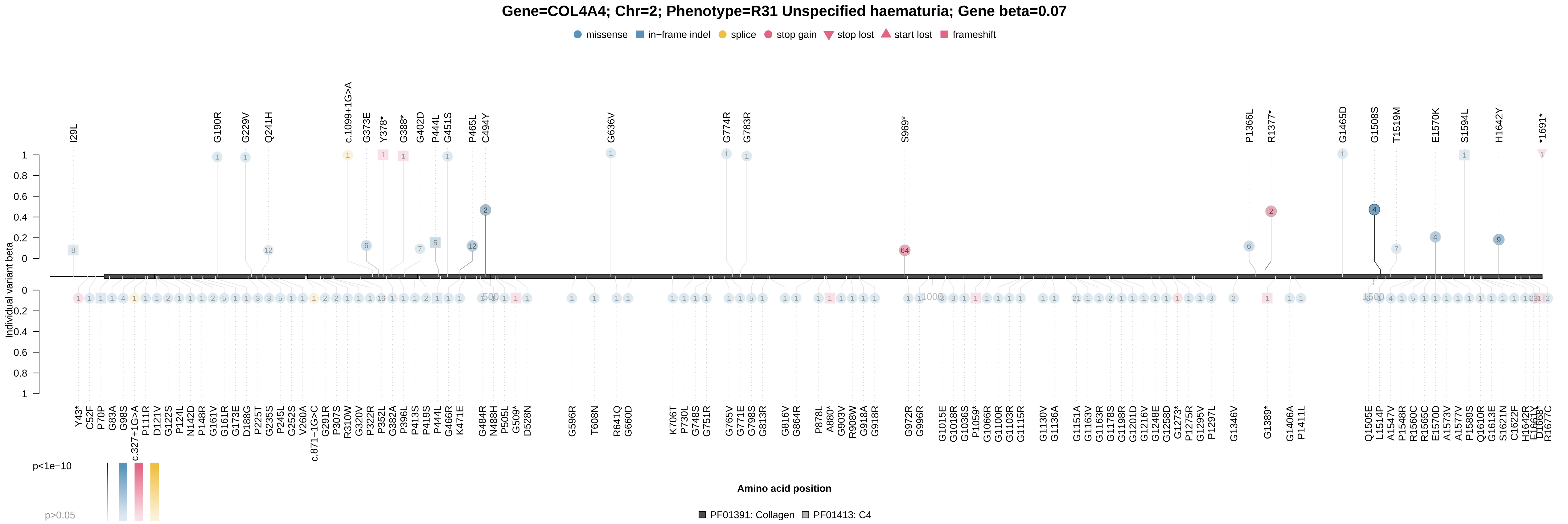

### CRP_x30710.pdf

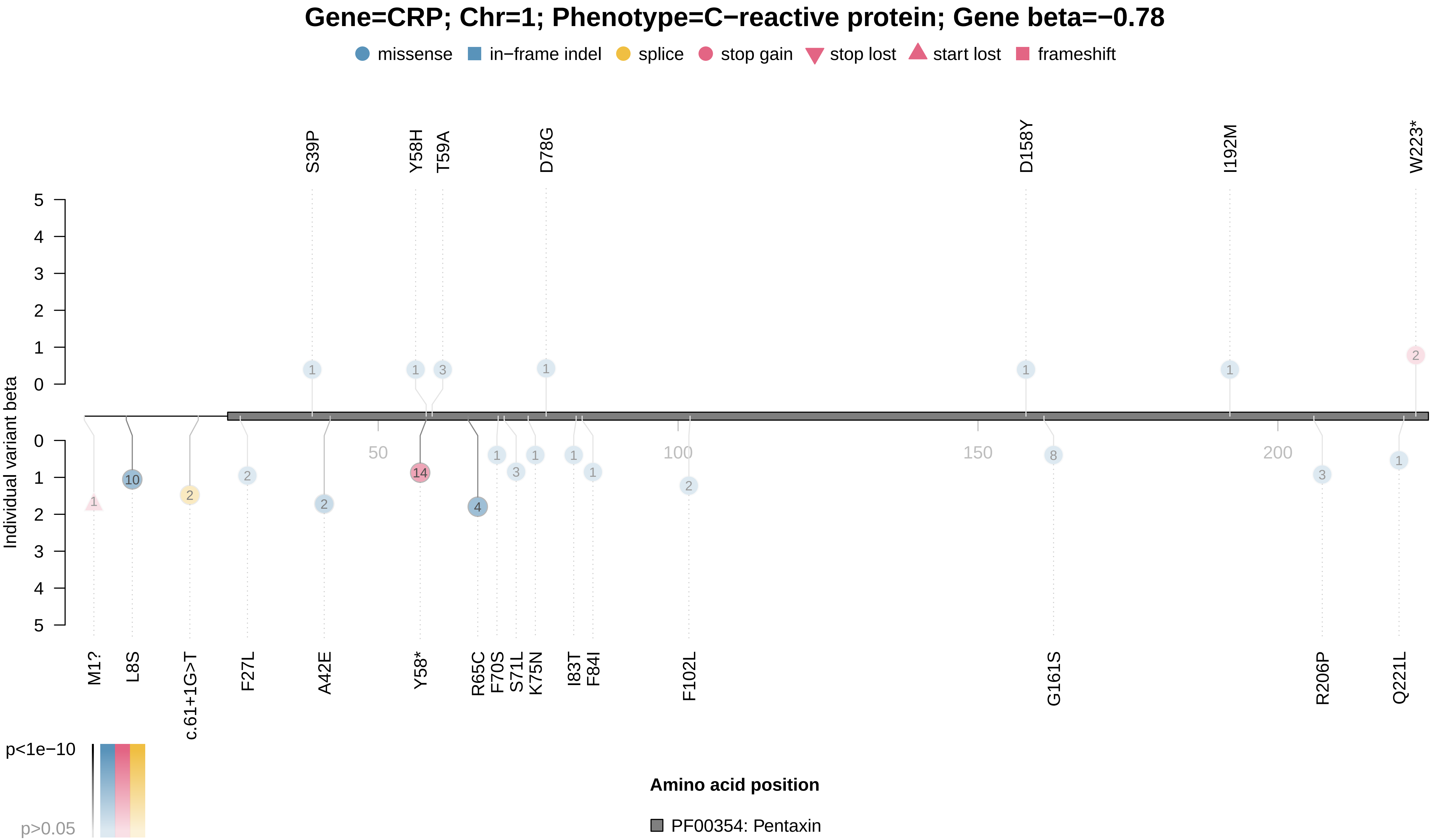

### CST3_x30720.pdf

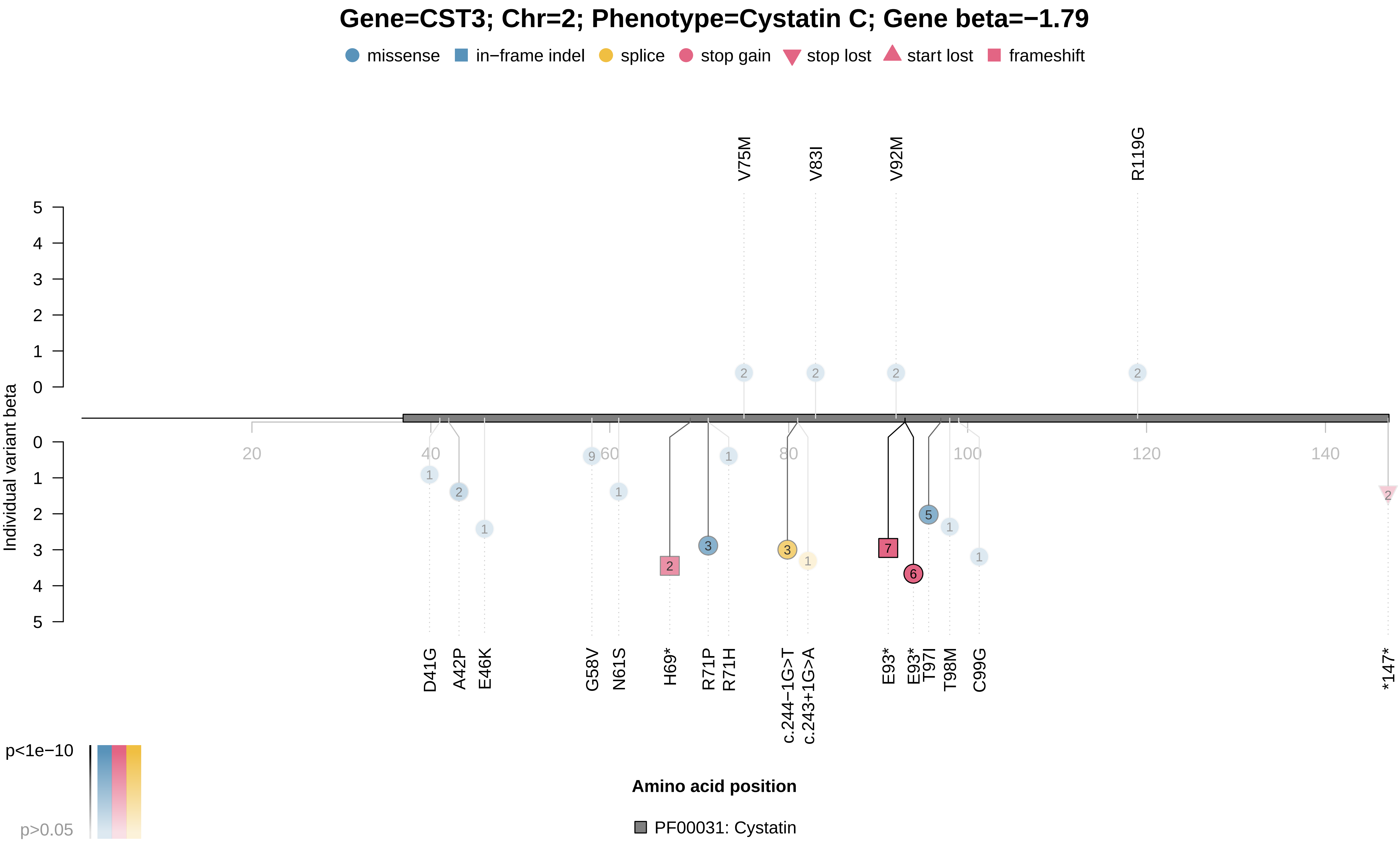

### CST3_x30720.pdf

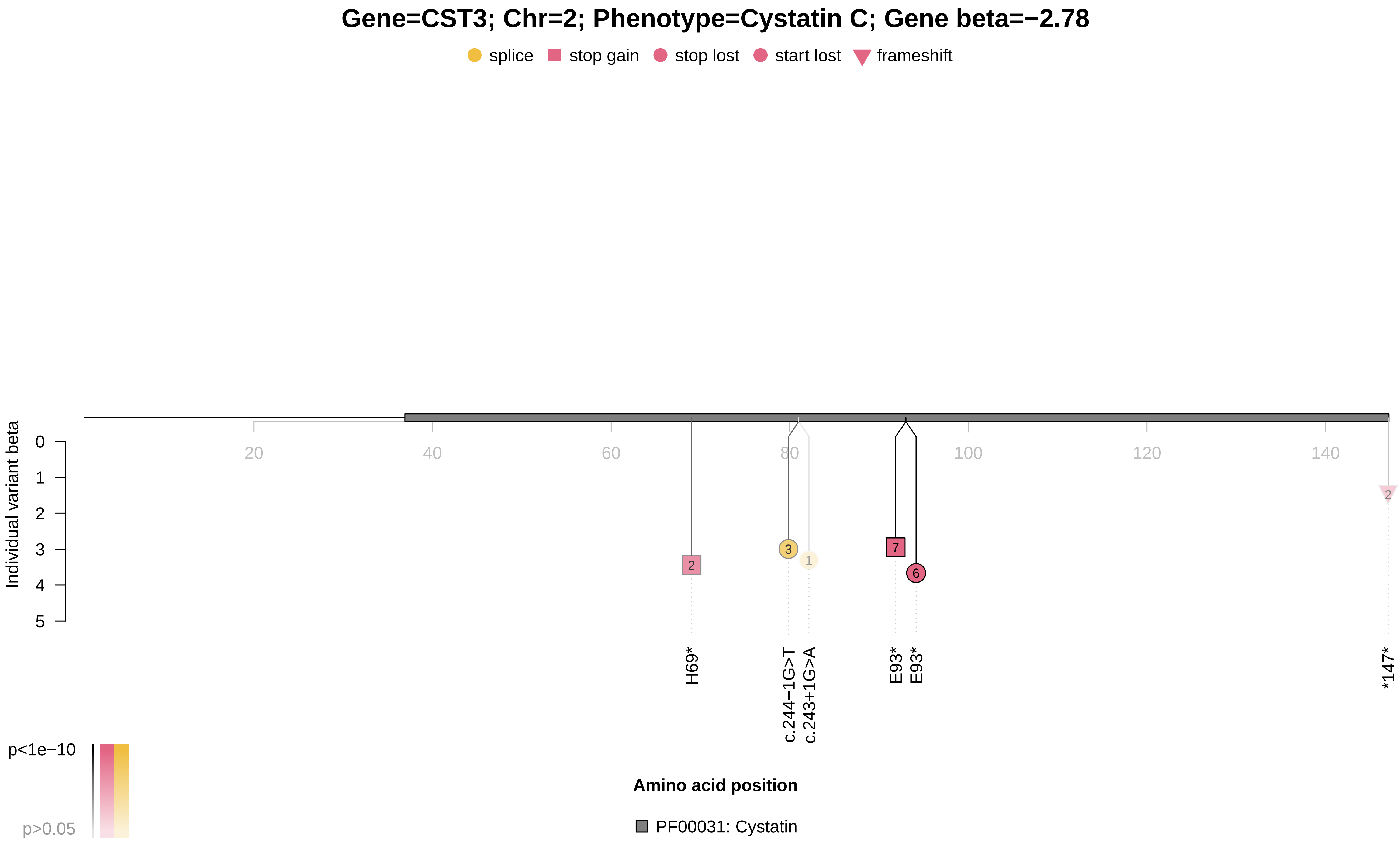

### GCK_x30740.pdf

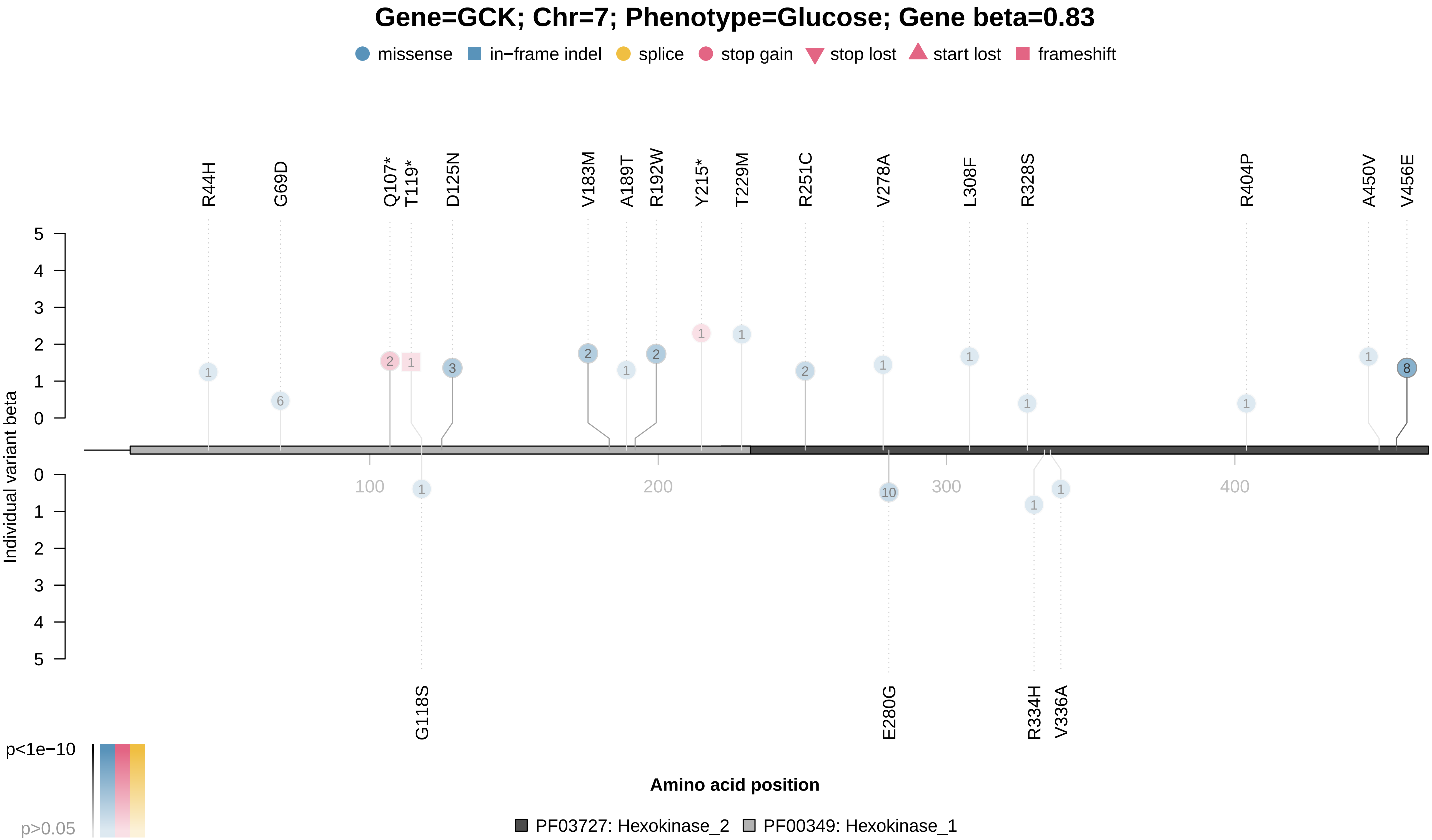

### GCK_x30750.pdf

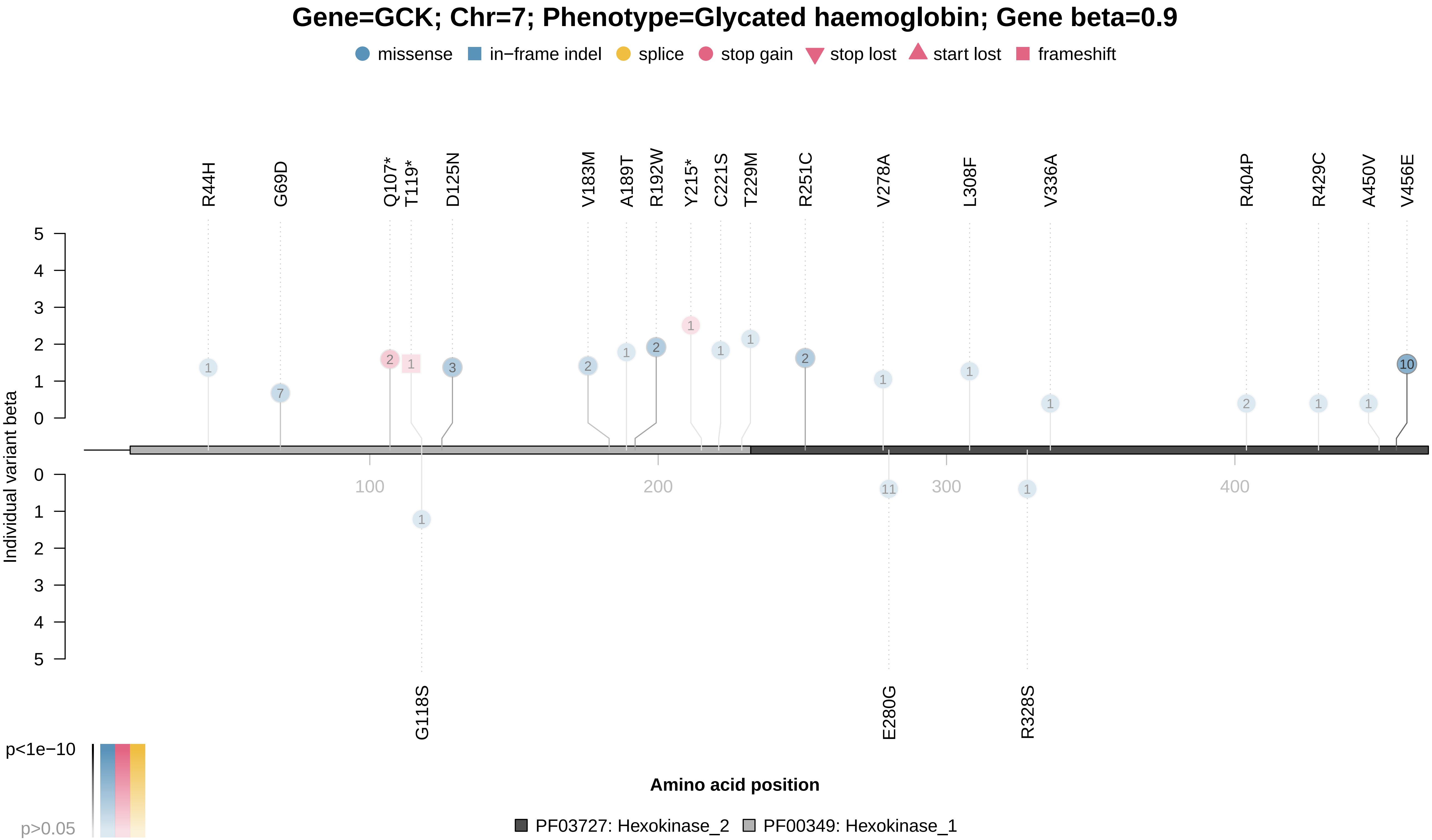

### GFI1B_x30100.pdf

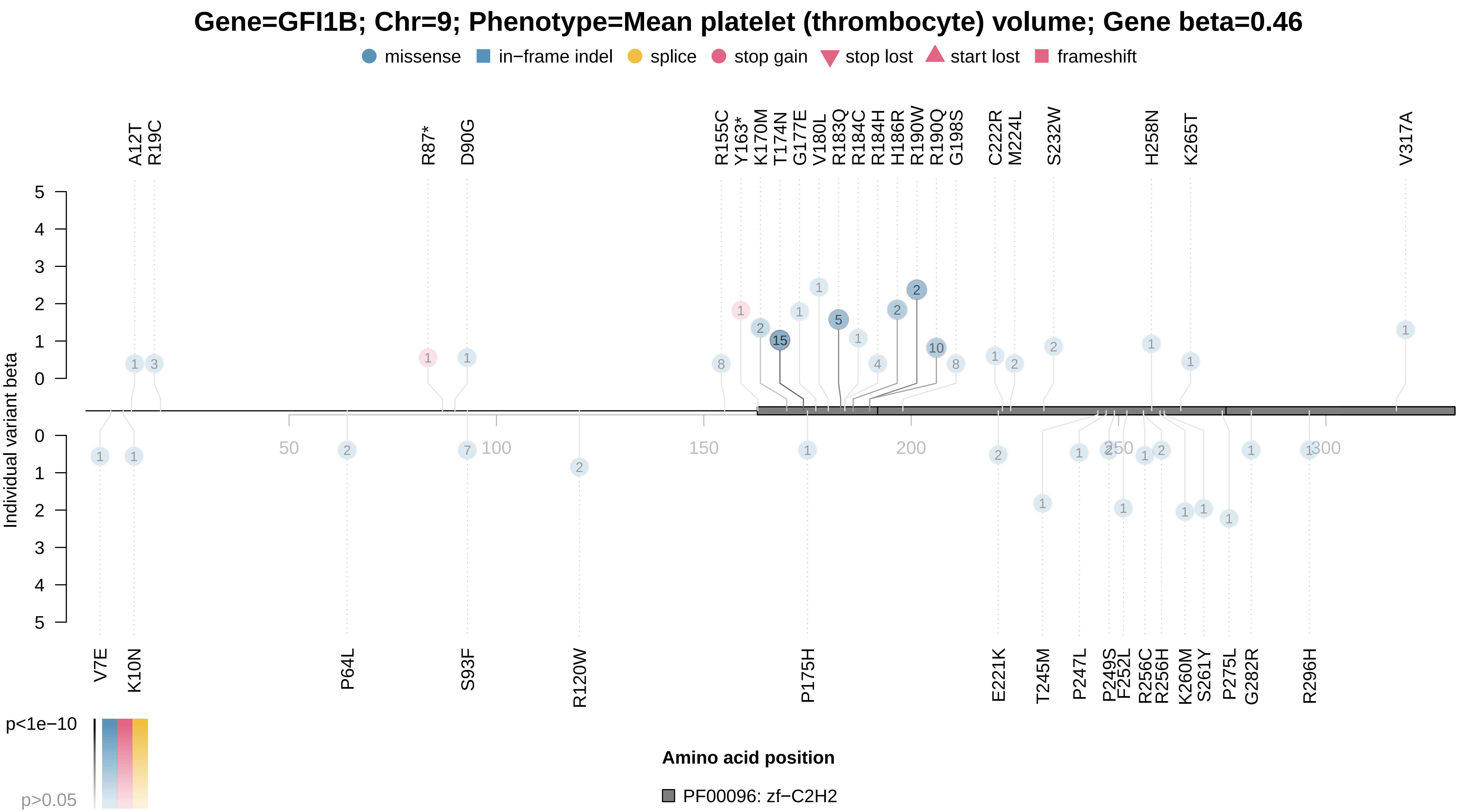

### GOT1_x30650.pdf

#### Gene=GOT1; Chr=1; Phenotype=Aspartate aminotransferase; Gene beta=−1.72

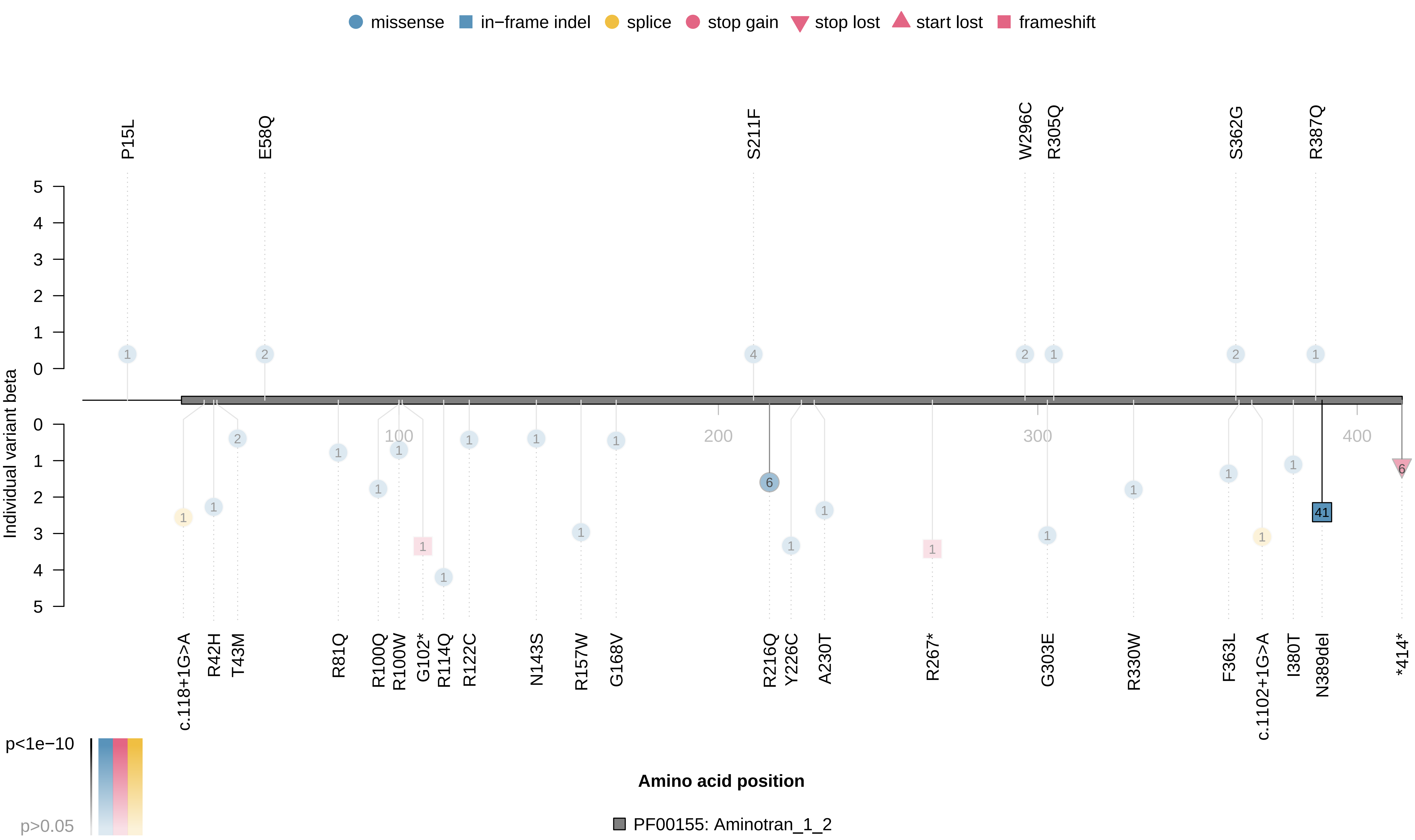

### GOT1_x30650.pdf

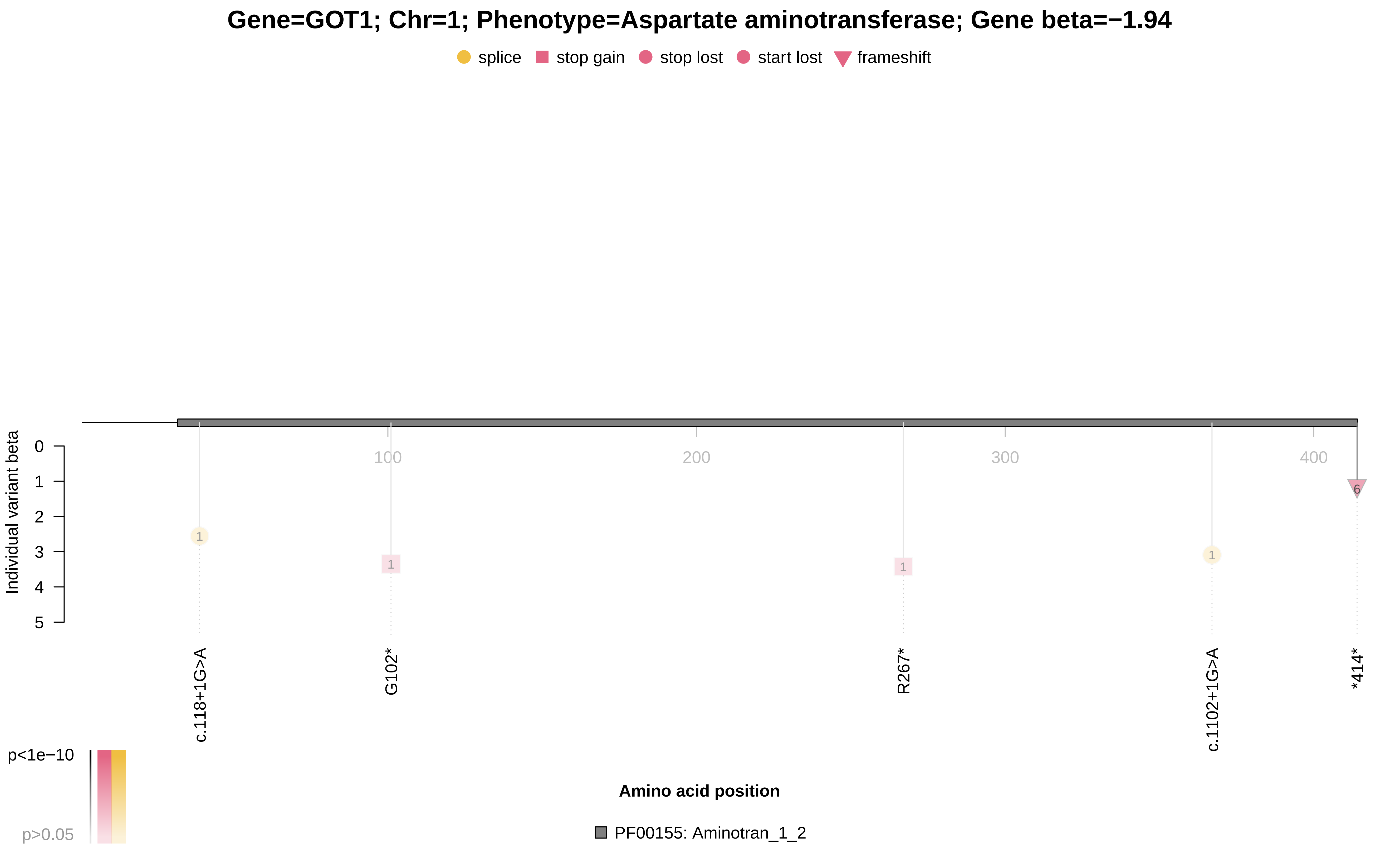

### GP1BA_x30100.pdf

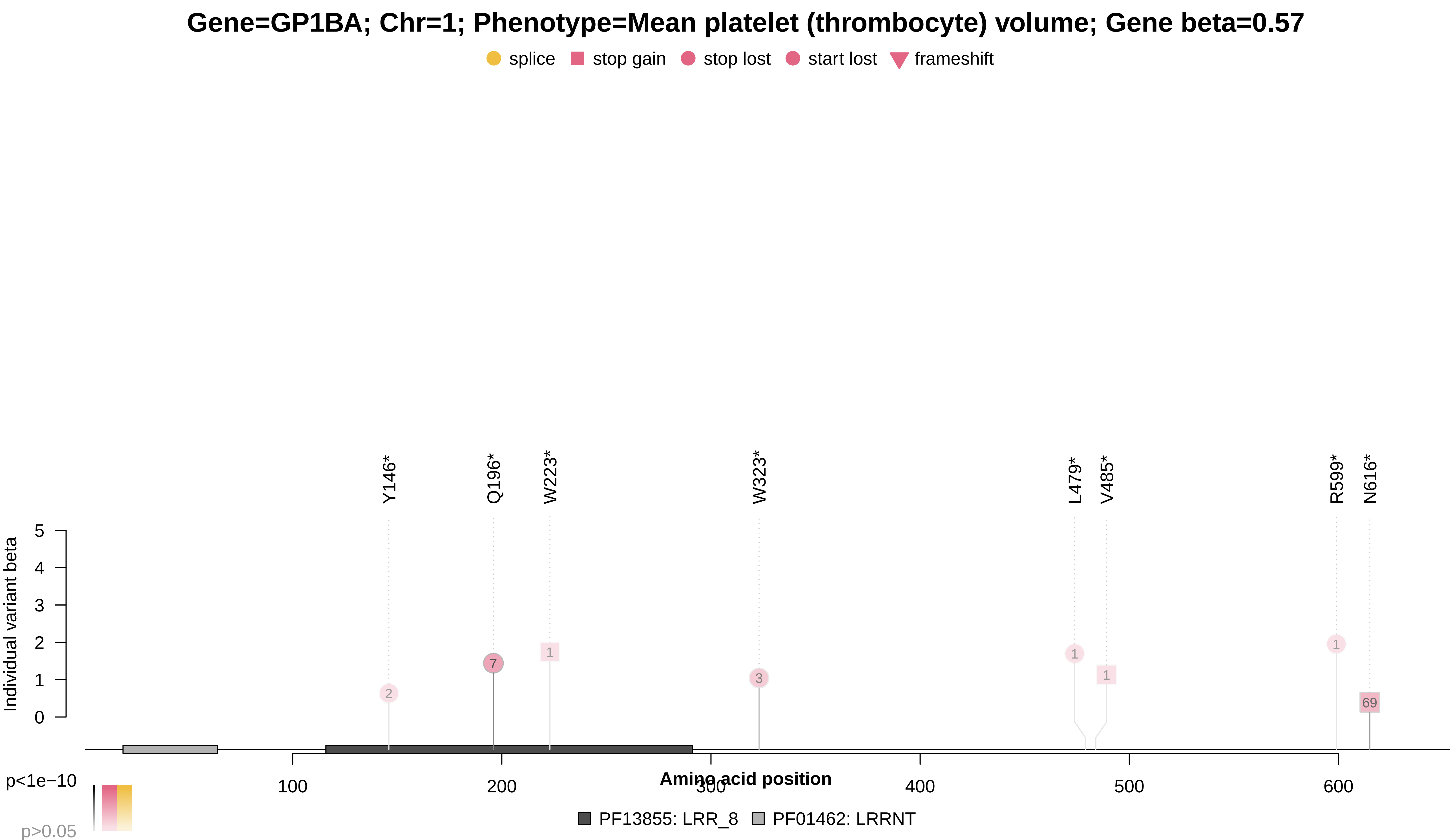

### GP1BB_x30100.pdf

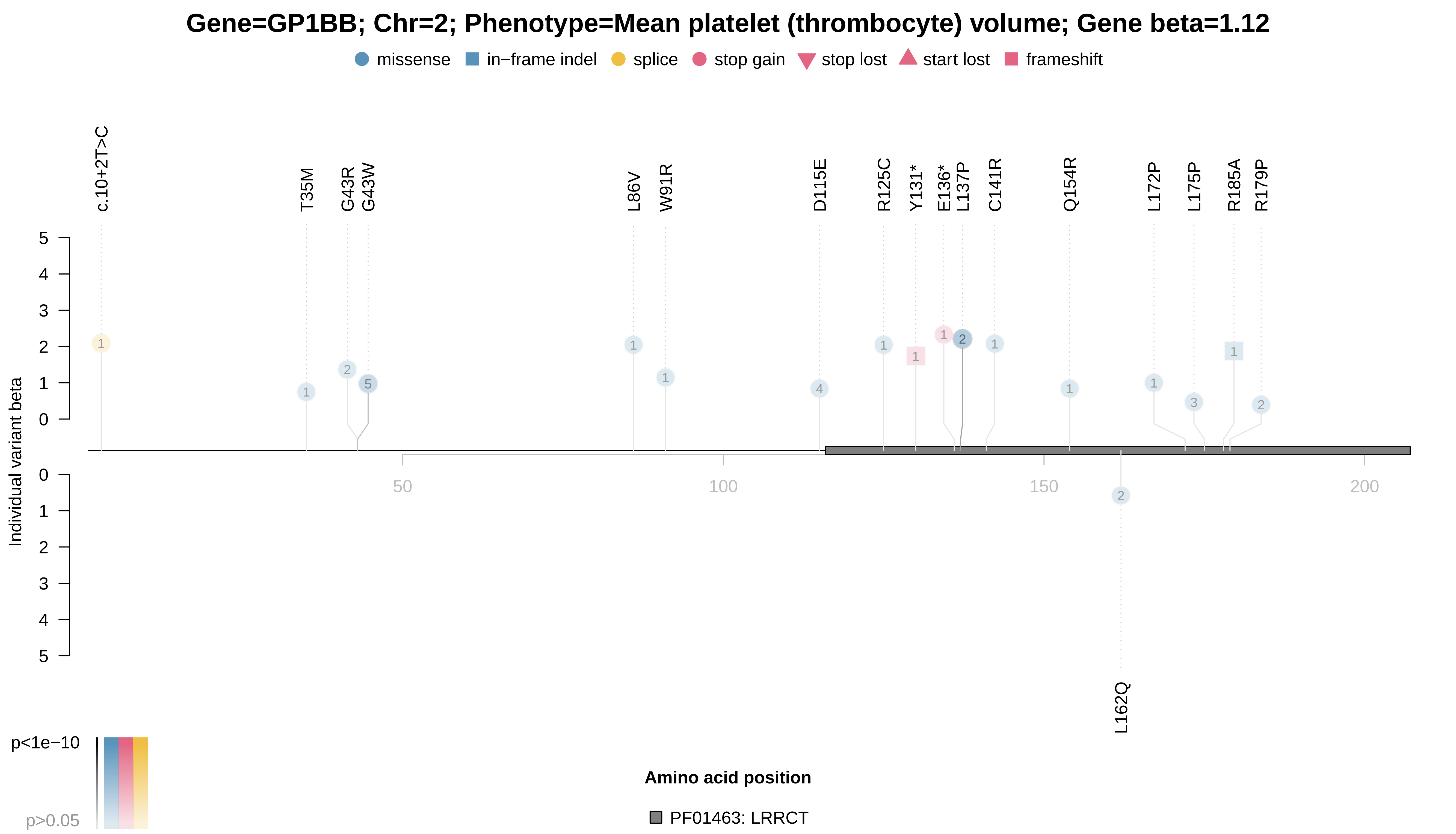

### GP9_x30080.pdf

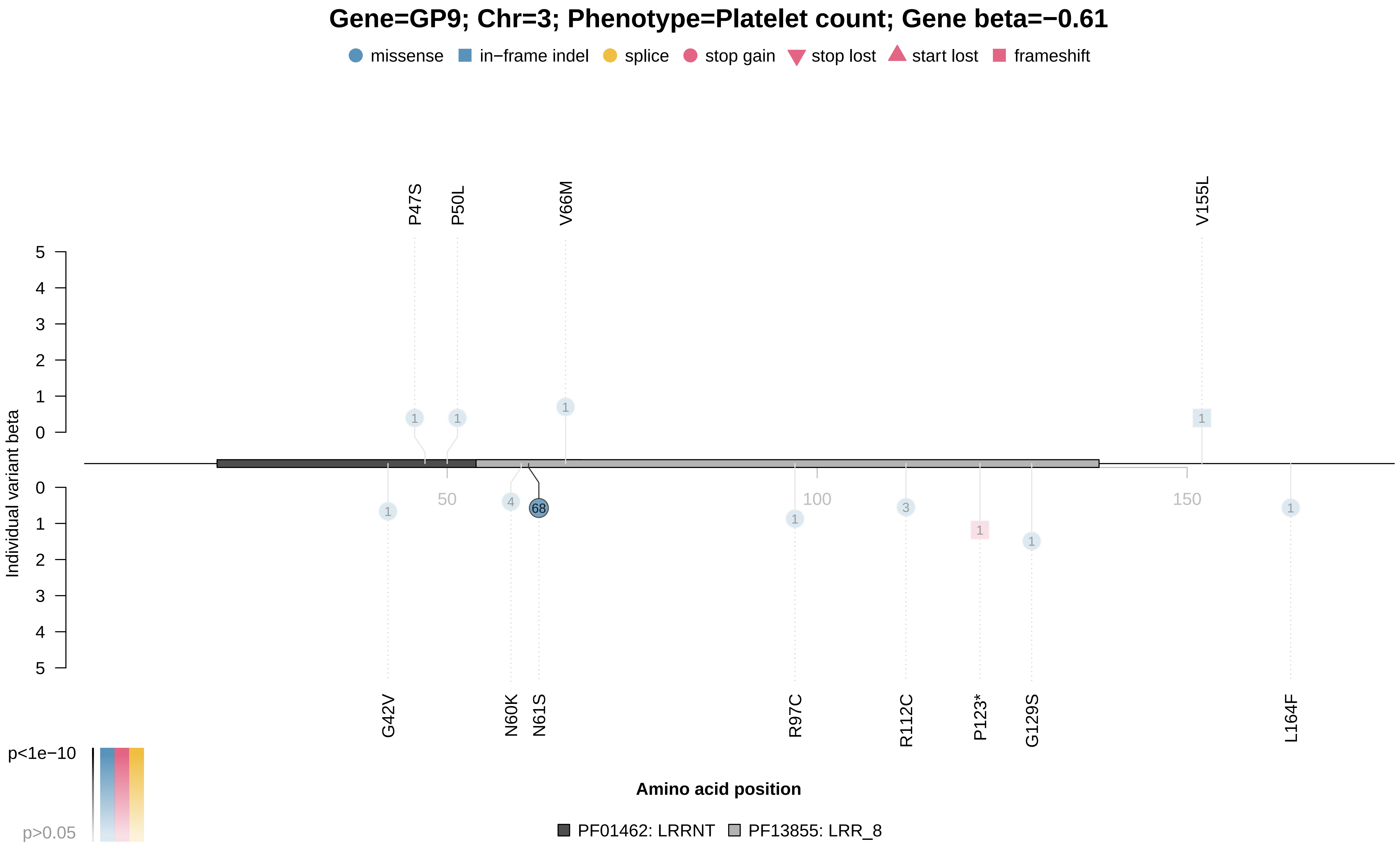

### GP9_x30100.pdf

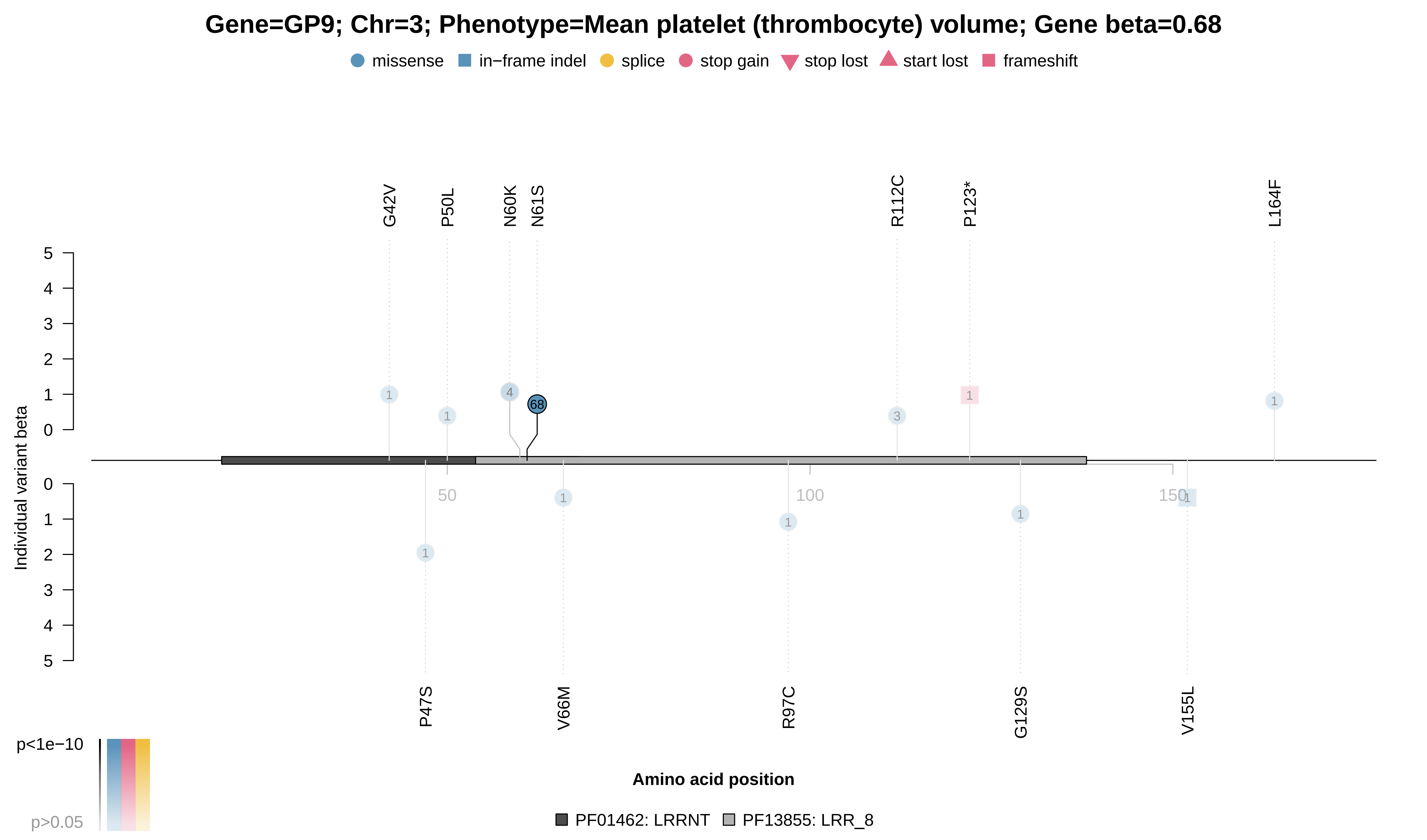

### GPLD1_x30610.pdf

#### Gene=GPLD1; Chr=6; Phenotype=Alkaline phosphatase; Gene beta=−0.64

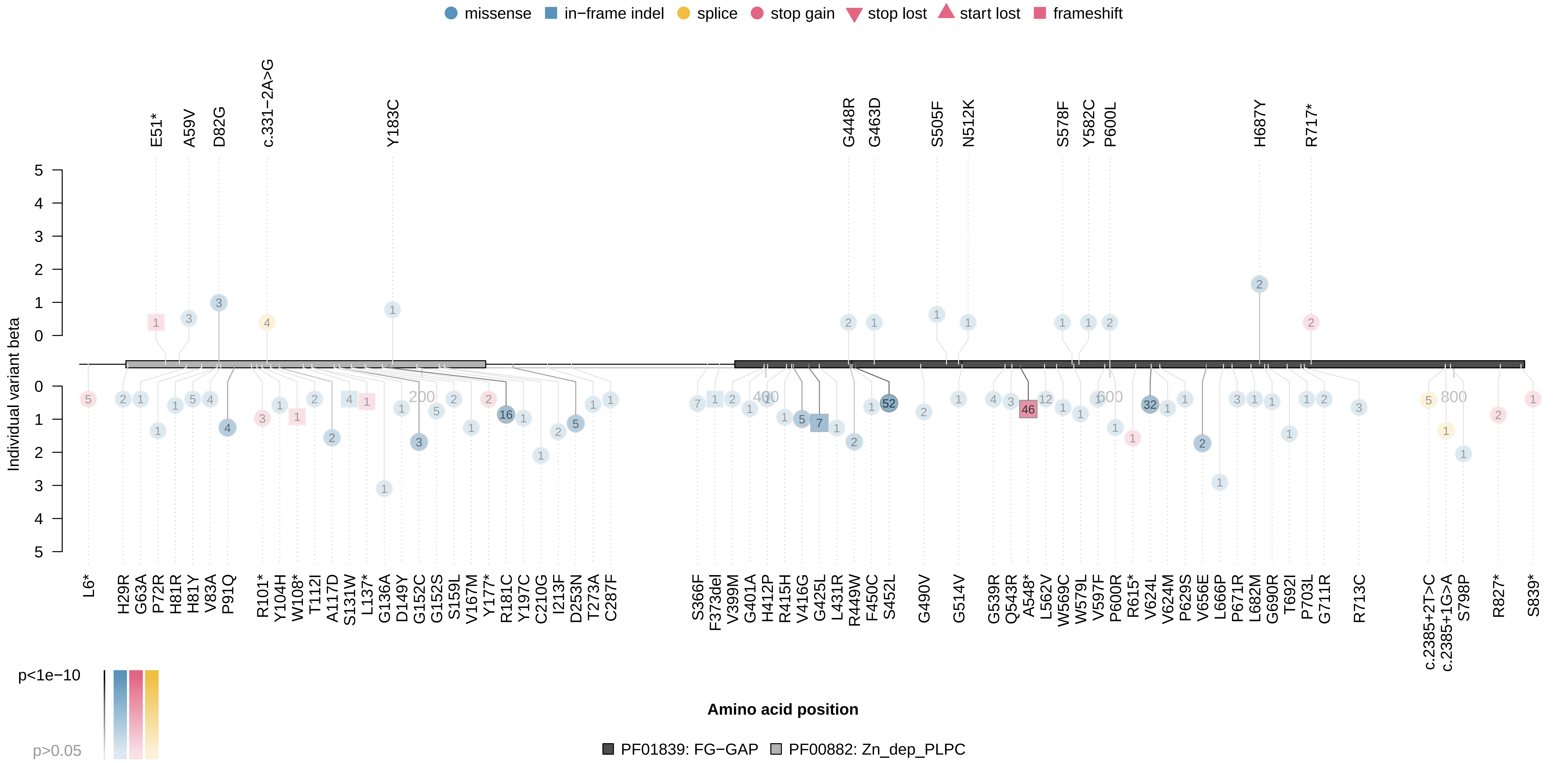

### GPLD1_x30610.pdf

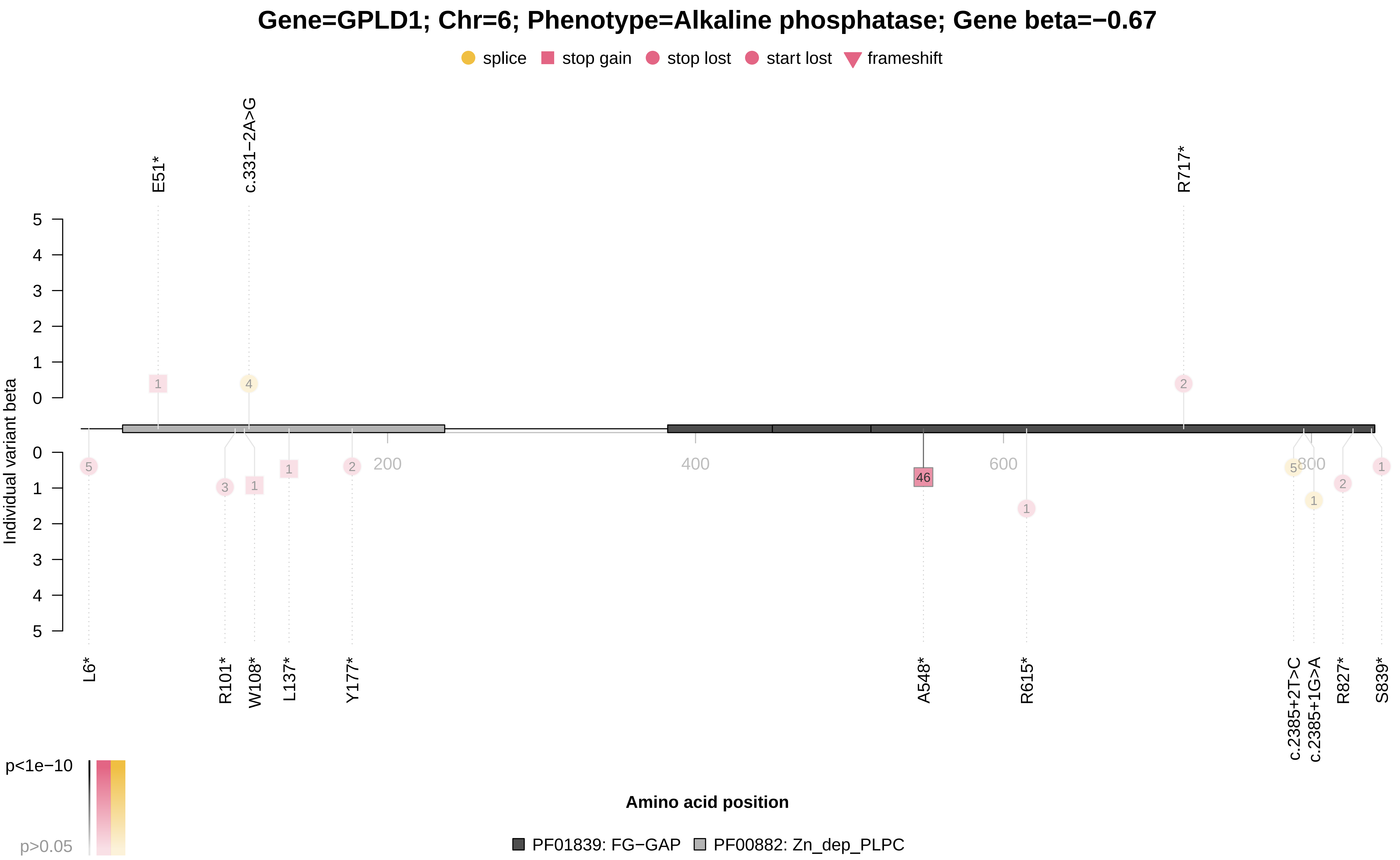

### GPT_x30620.pdf

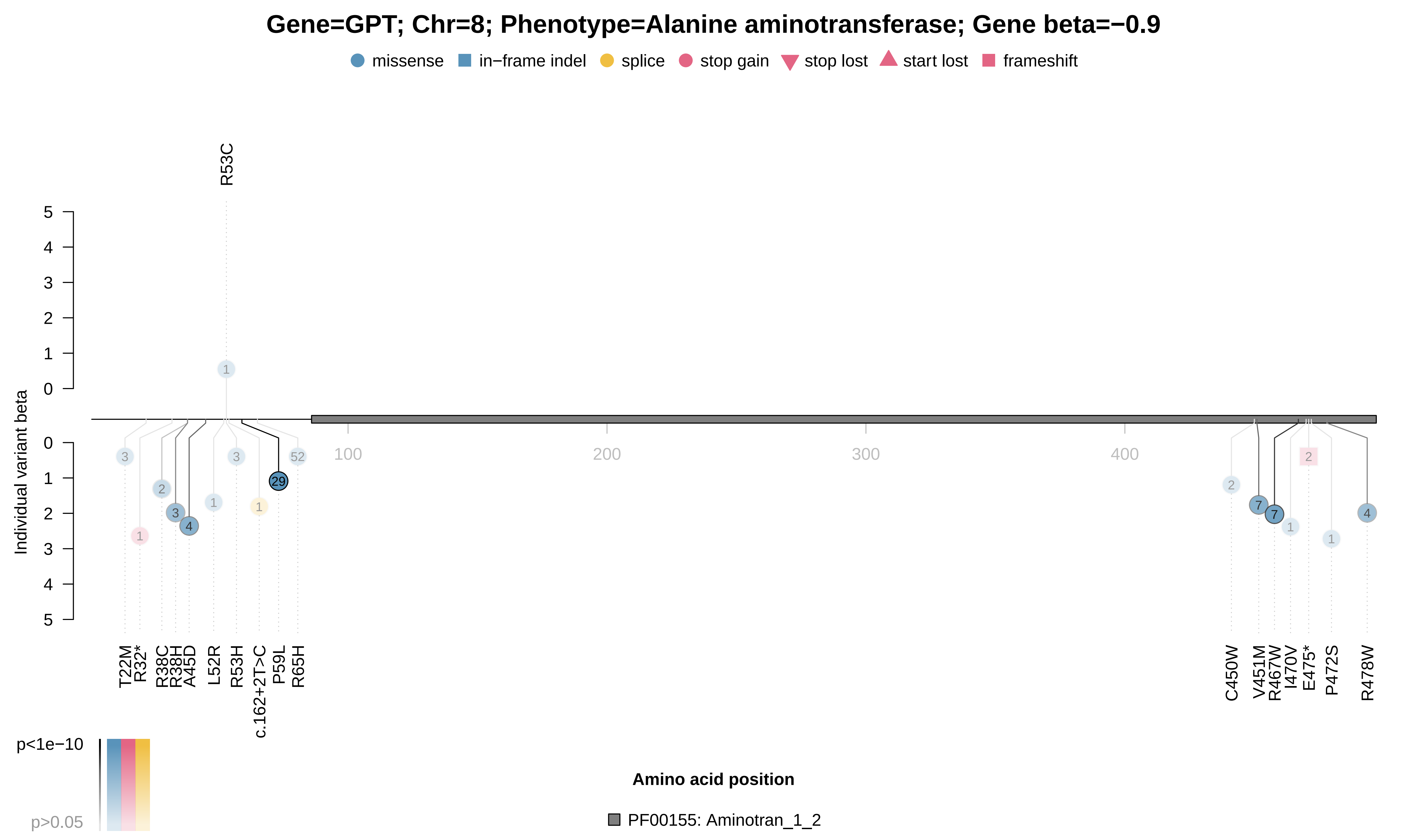

### IQGAP2_x30100.pdf

**Gene=IQGAP2; Chr=5; Phenotype=Mean platelet (thrombocyte) volume; Gene beta=0.24**

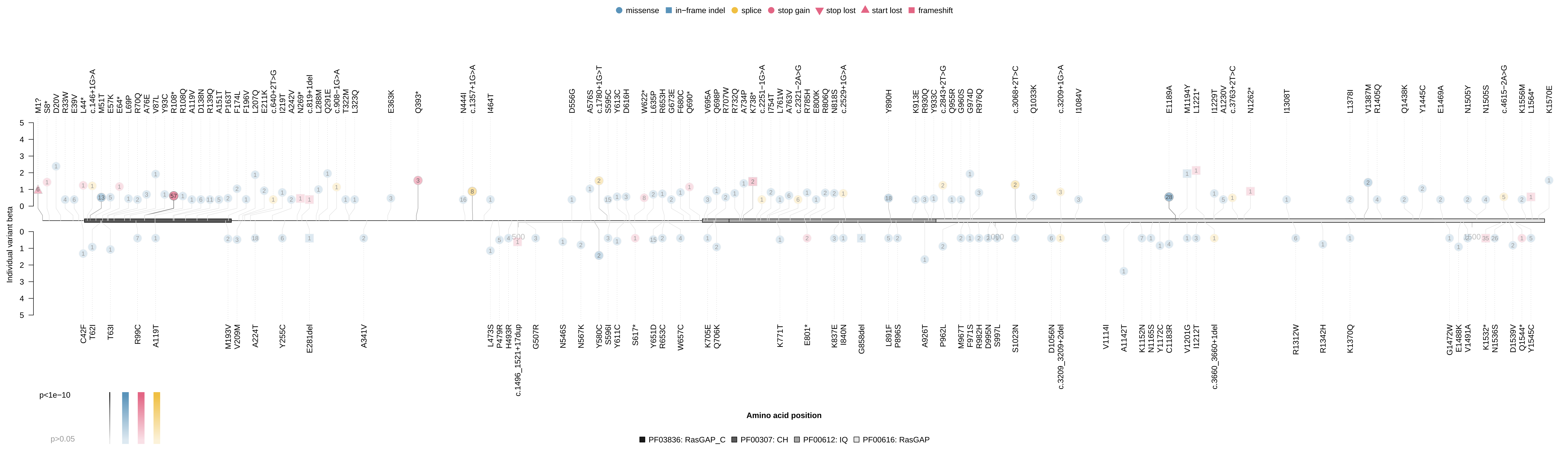

### IQGAP2_x30100.pdf

#### Gene=IQGAP2; Chr=5; Phenotype=Mean platelet (thrombocyte) volume; Gene beta=0.55

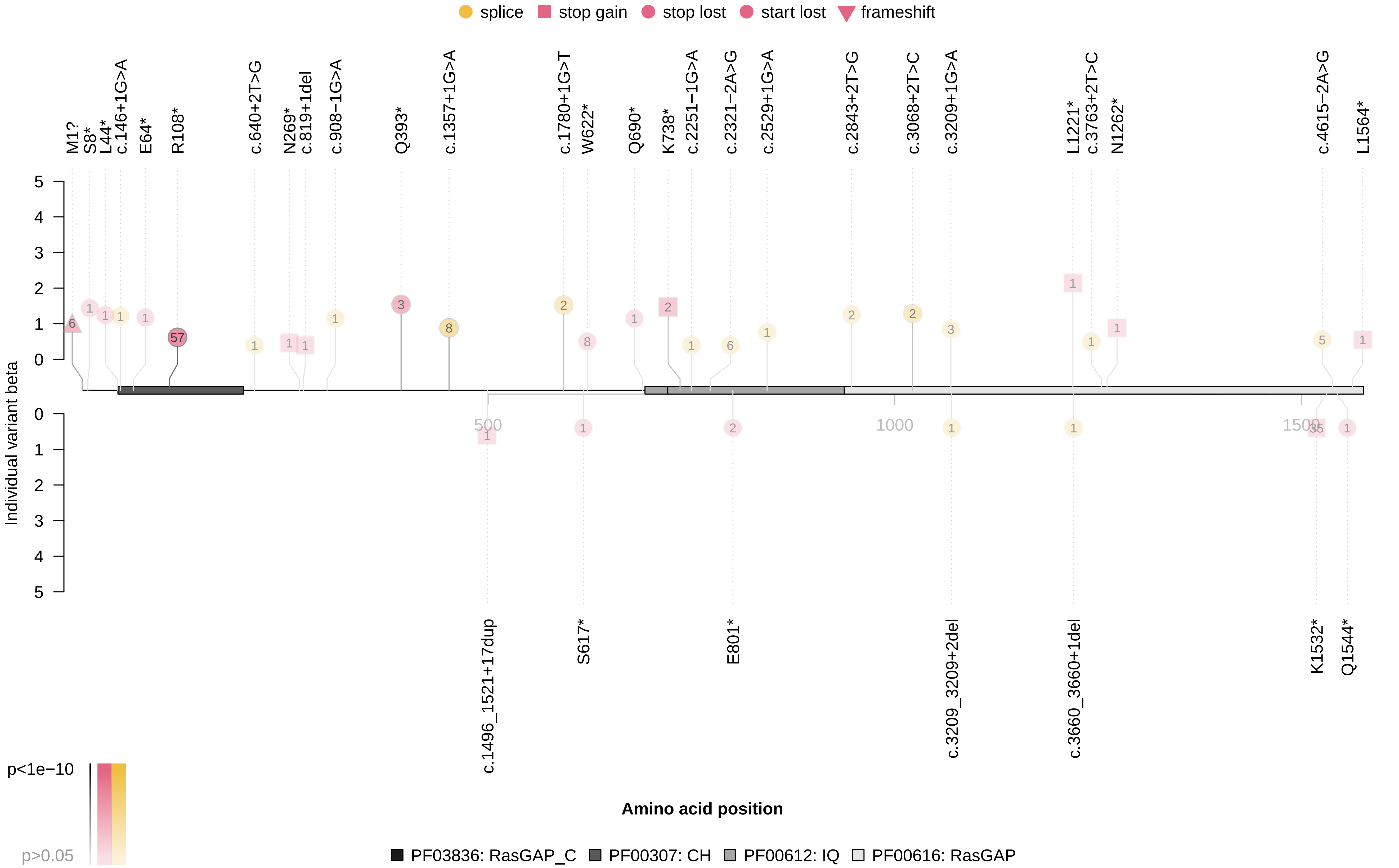

### ITGA2B_x30080.pdf

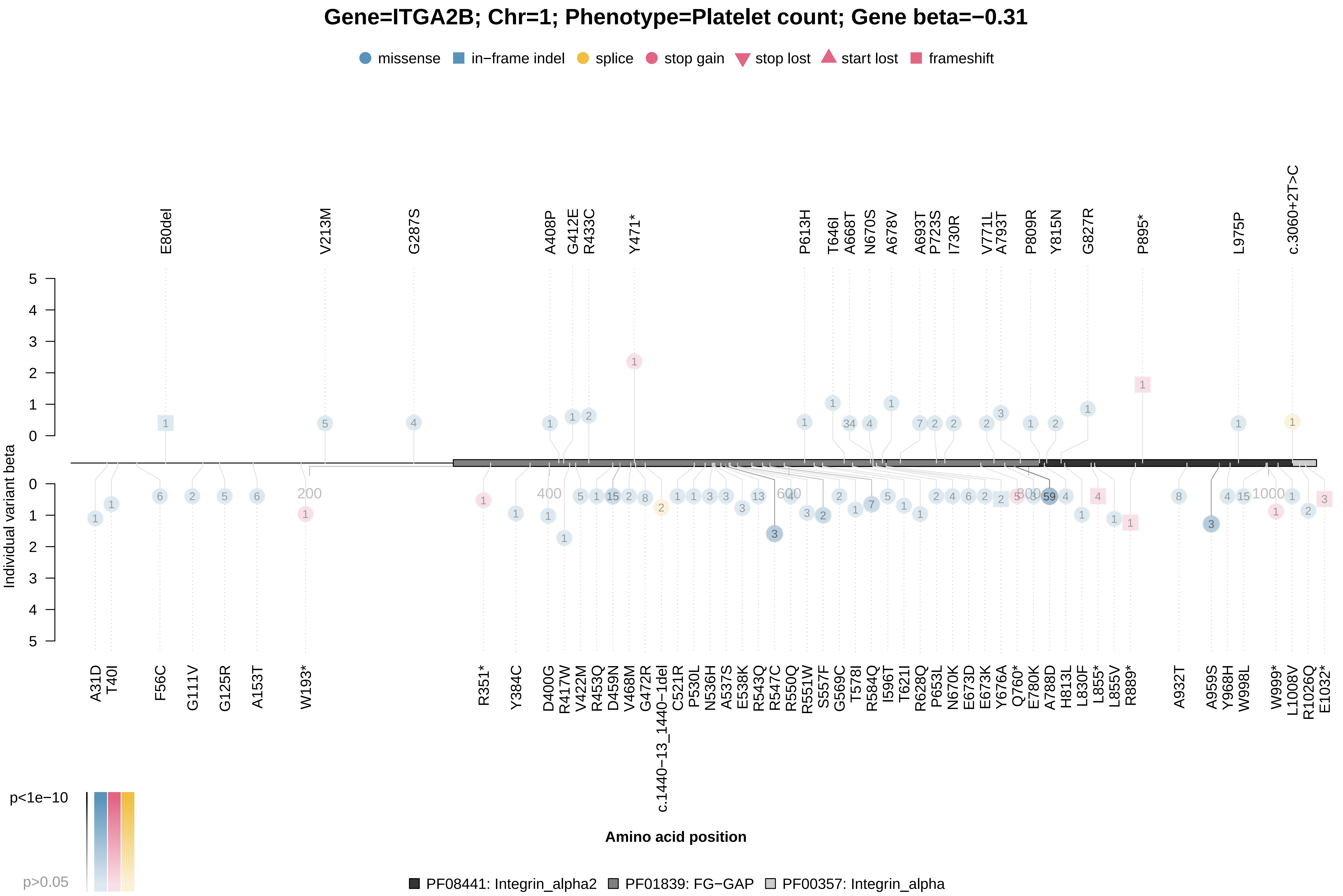

### JAK2_x30080.pdf

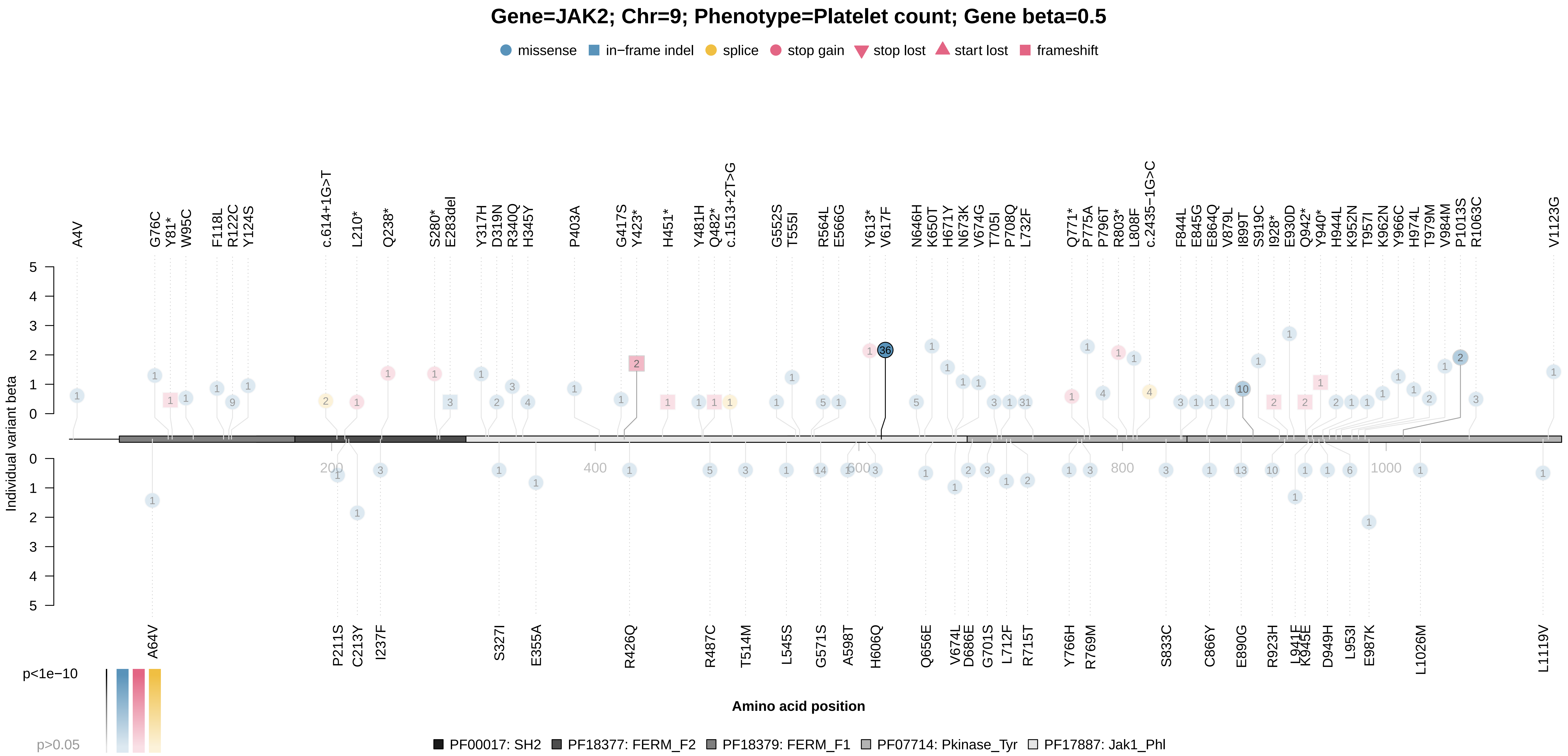

### JAK2_x30090.pdf

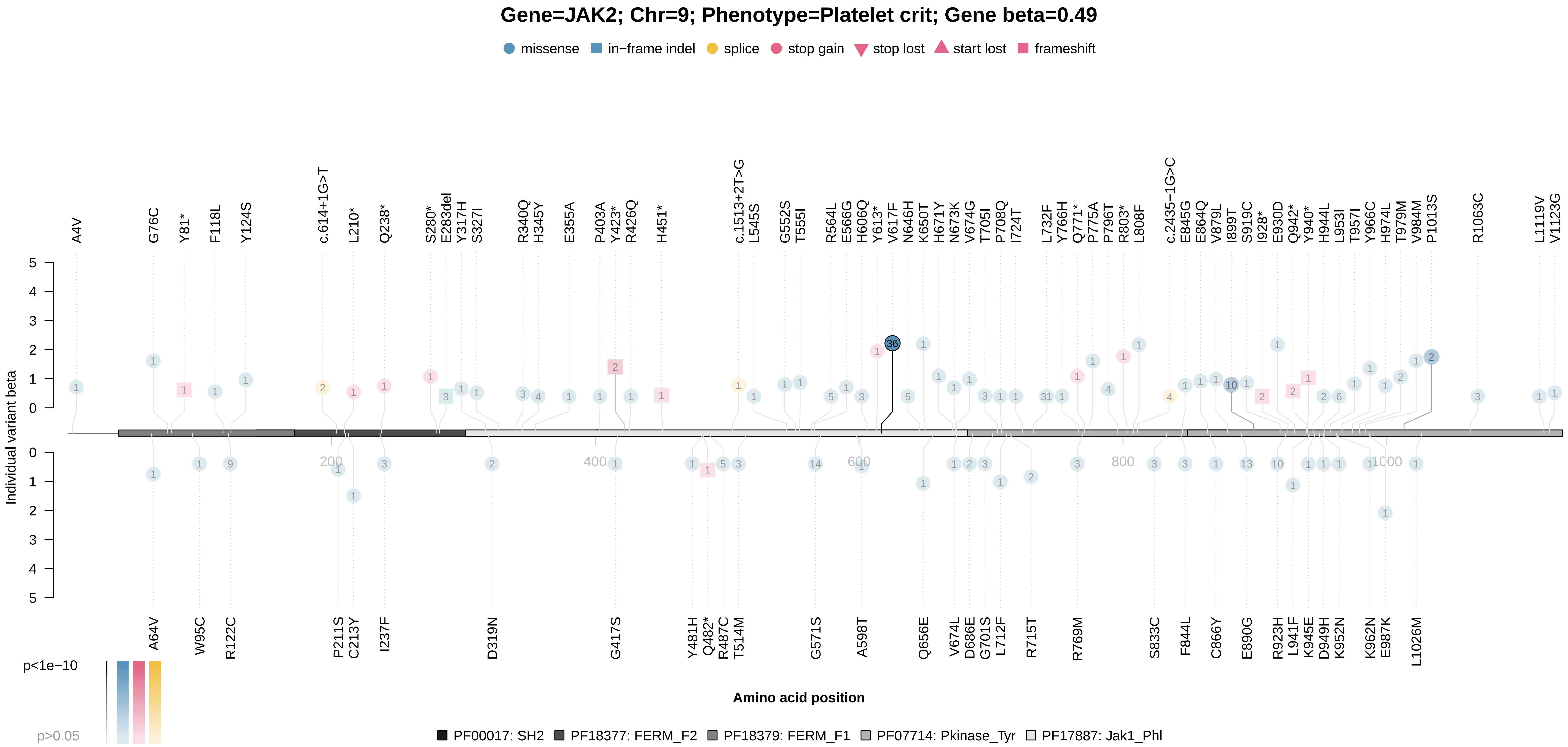

### KLF1_x30040.pdf

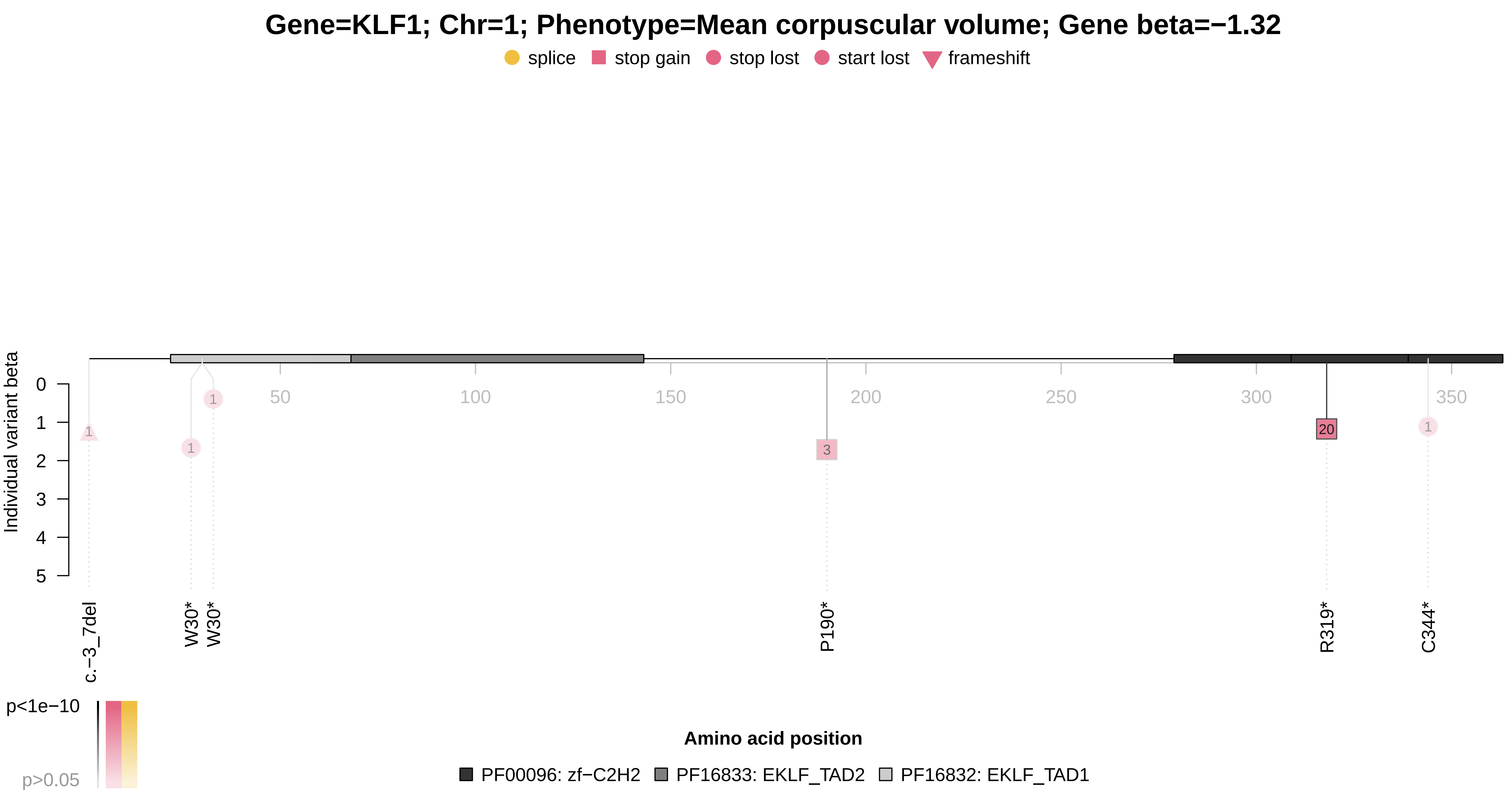

### KLF1_x30050.pdf

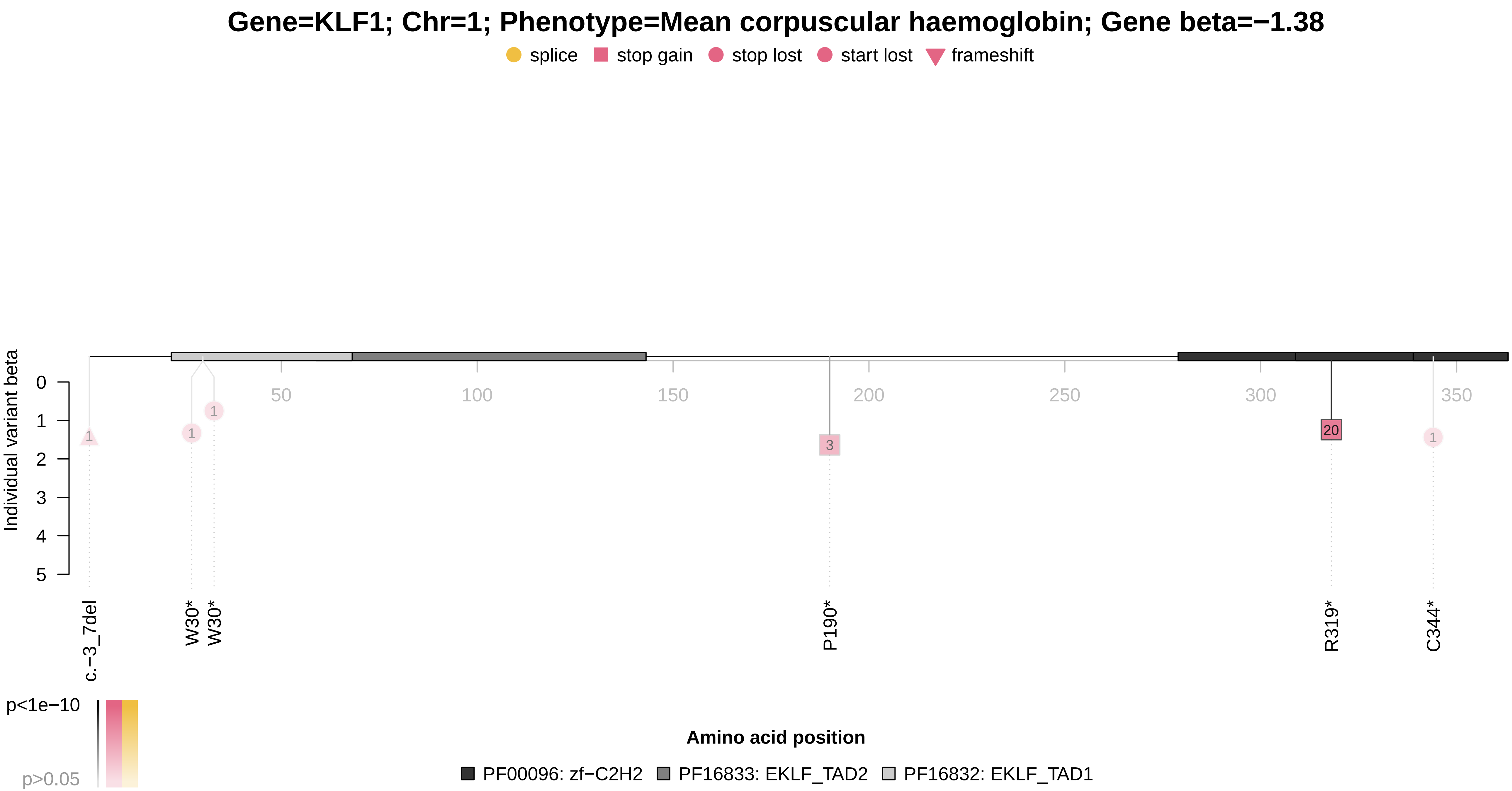

### KLF1_x30070.pdf

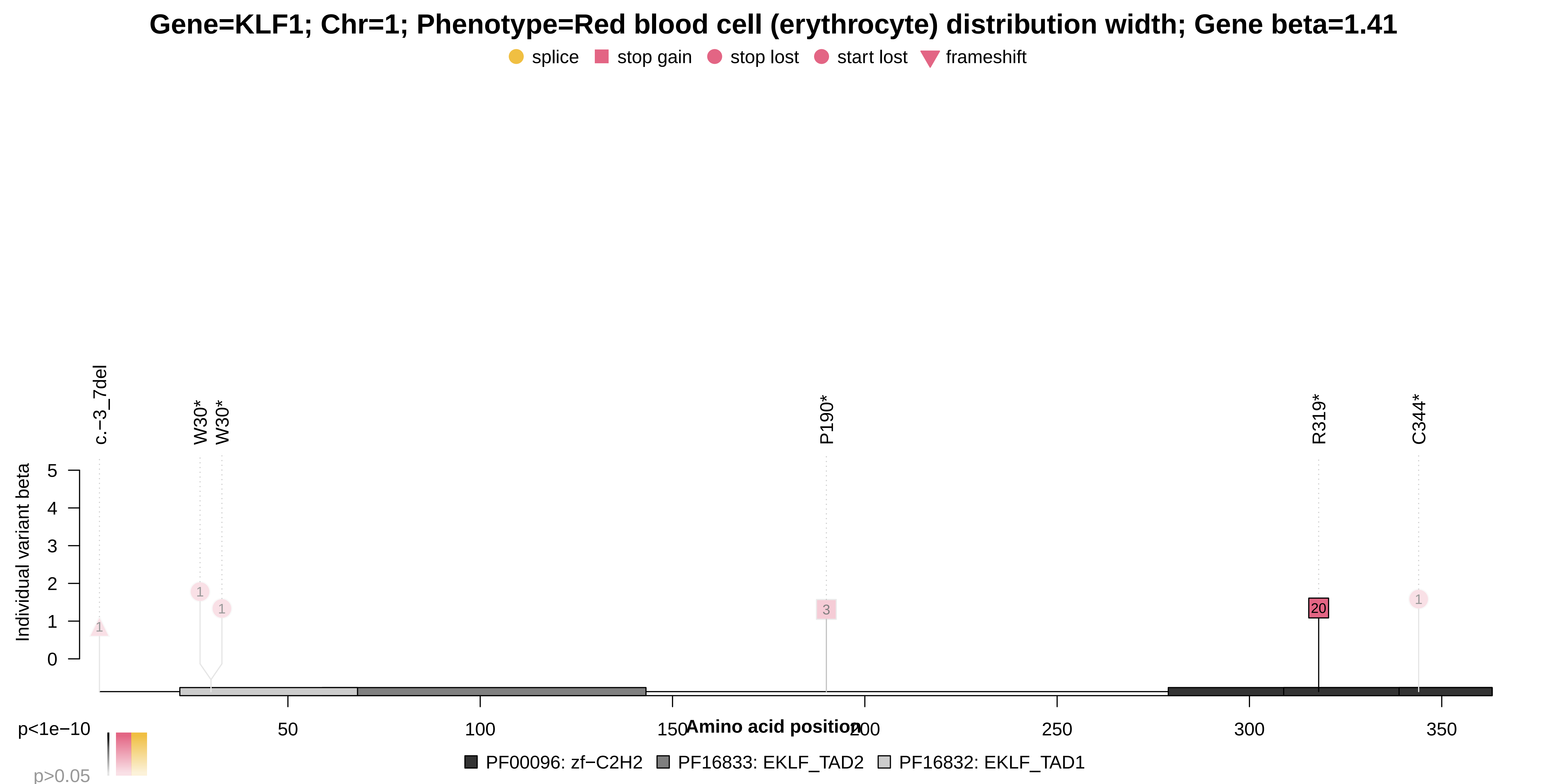

### LCAT_x30630.pdf

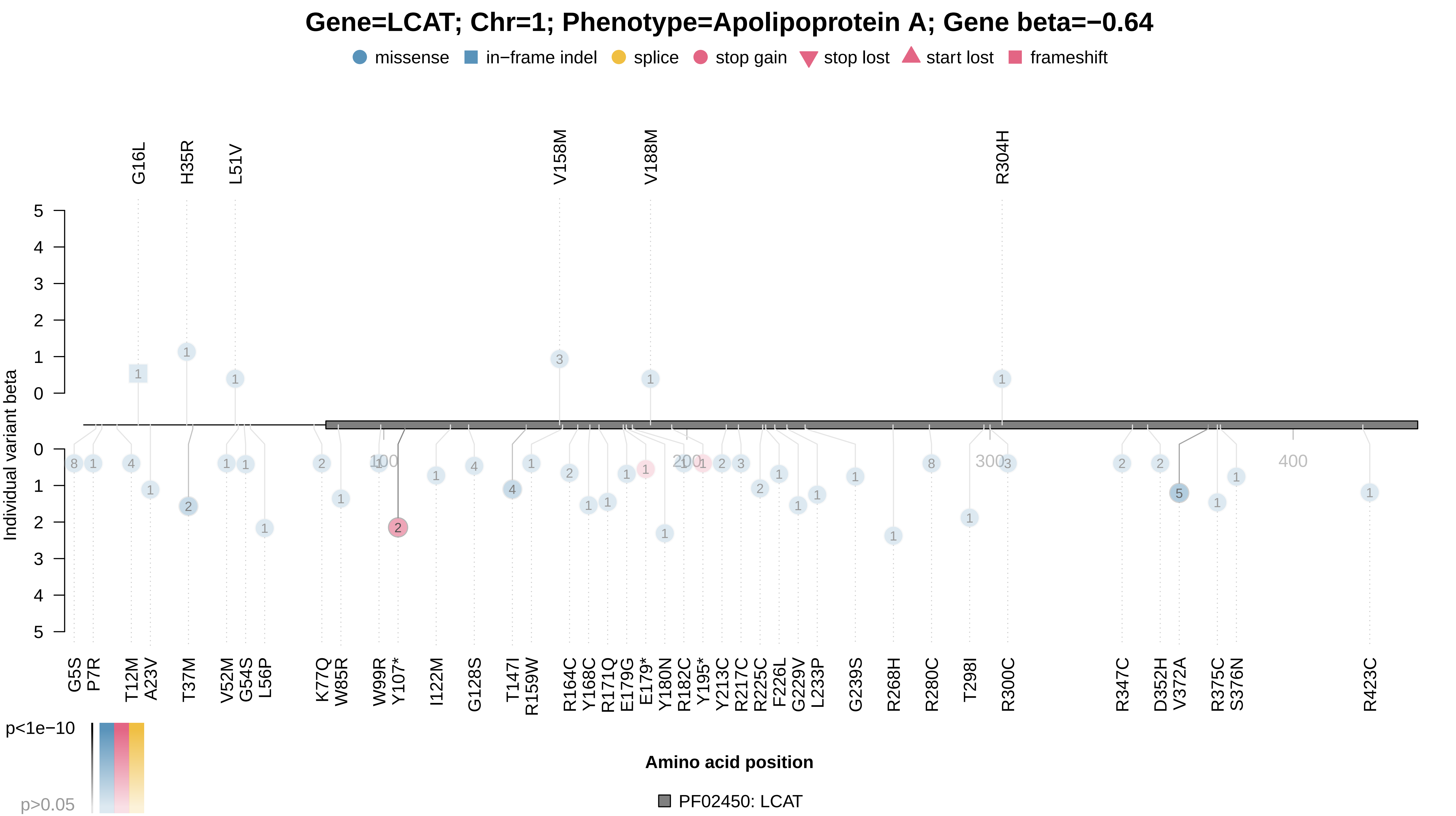

### LCAT_x30760.pdf

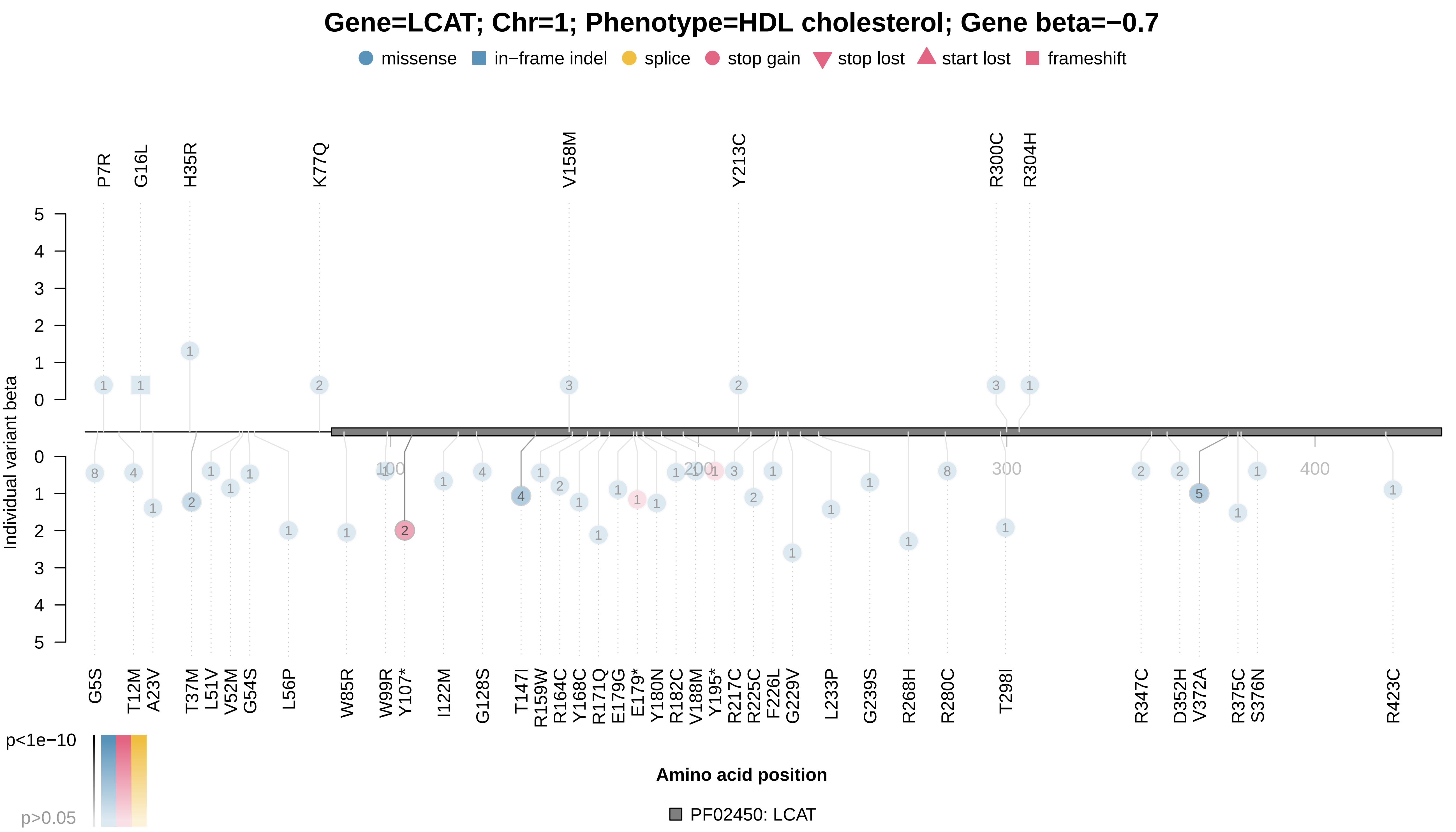

### LDLR_x6153_1.pdf

**Gene=LDLR; Chr=1; Phenotype=Cholesterol lowering medication; Gene beta=0.18**

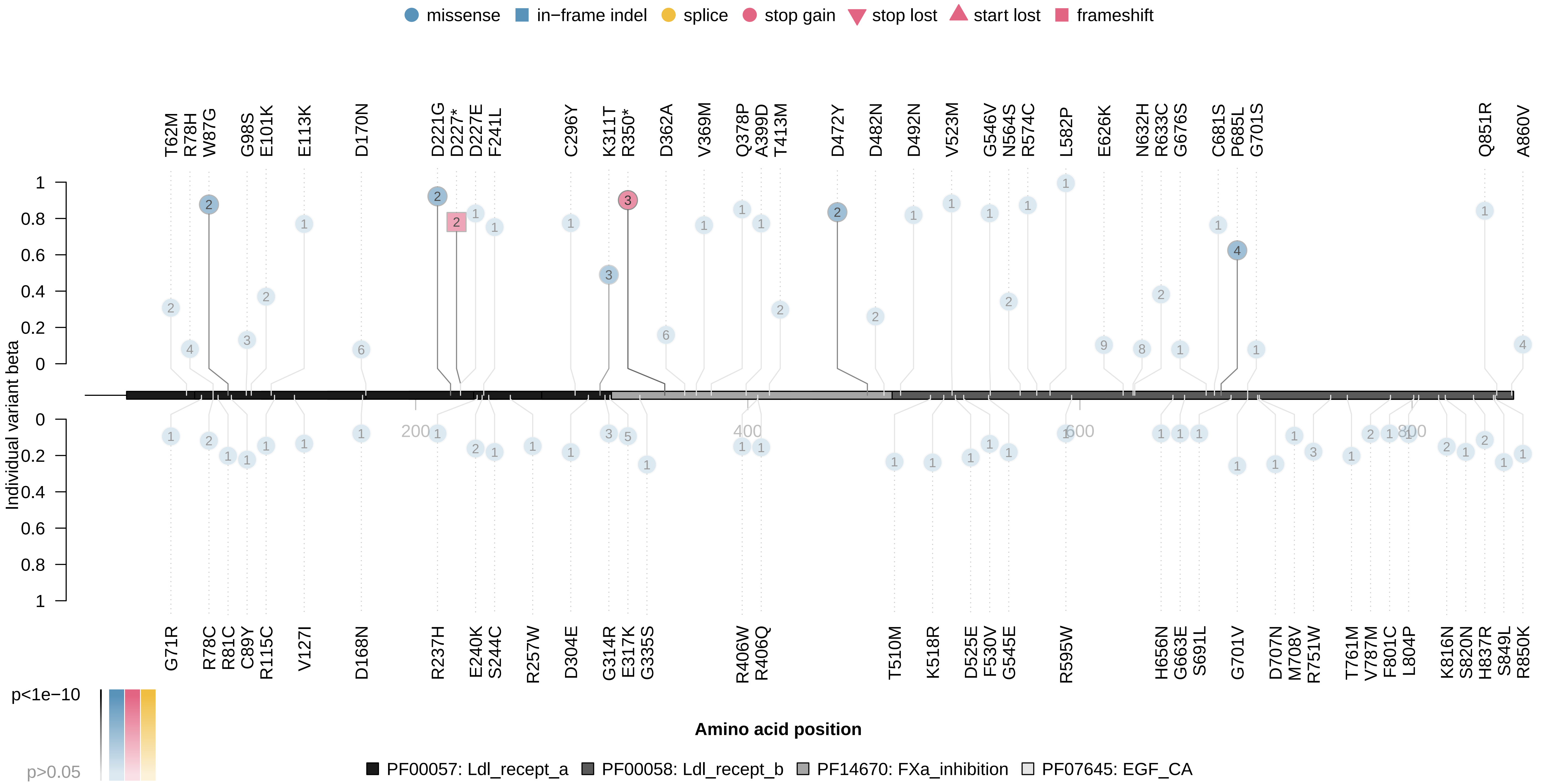

### LDLR_x20002_1473.pdf

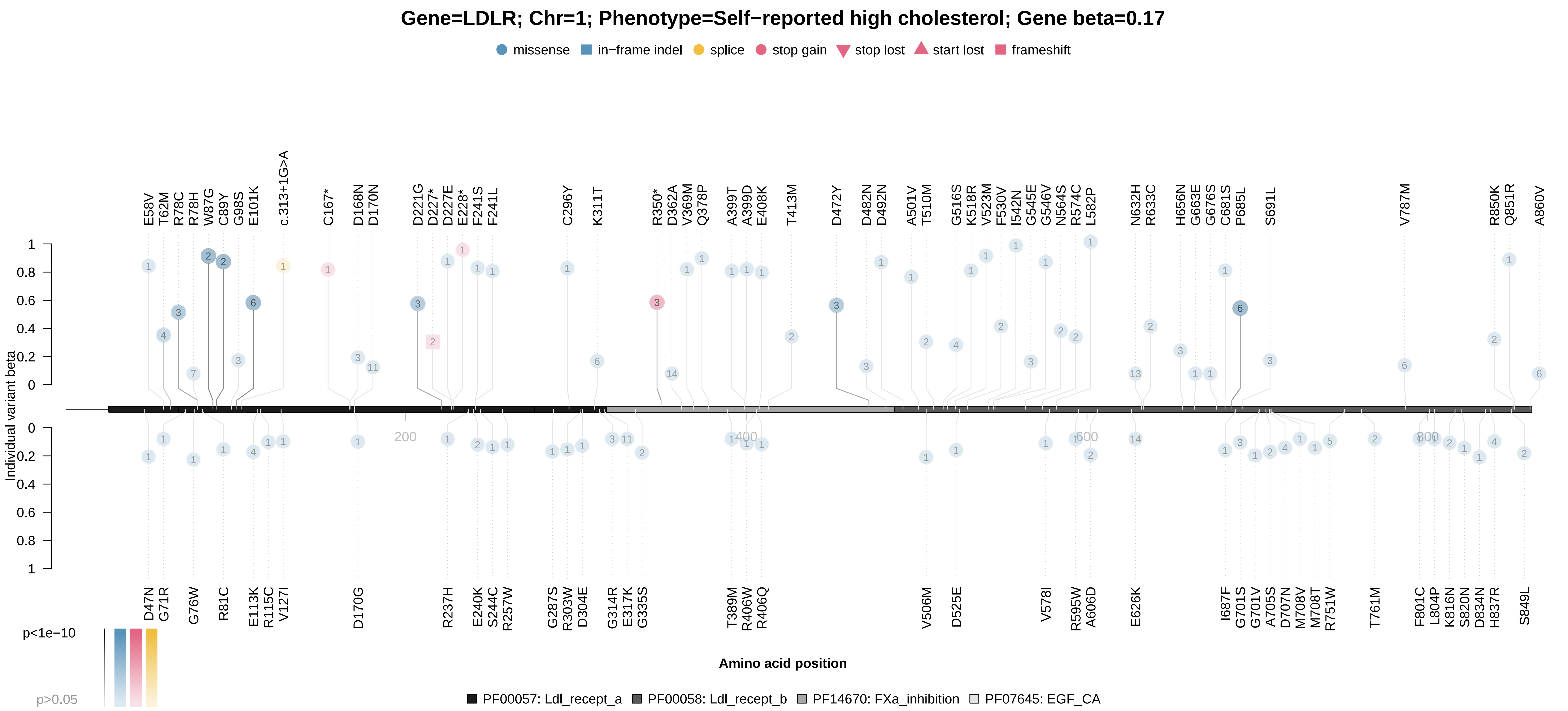

### LIPC_x30630.pdf

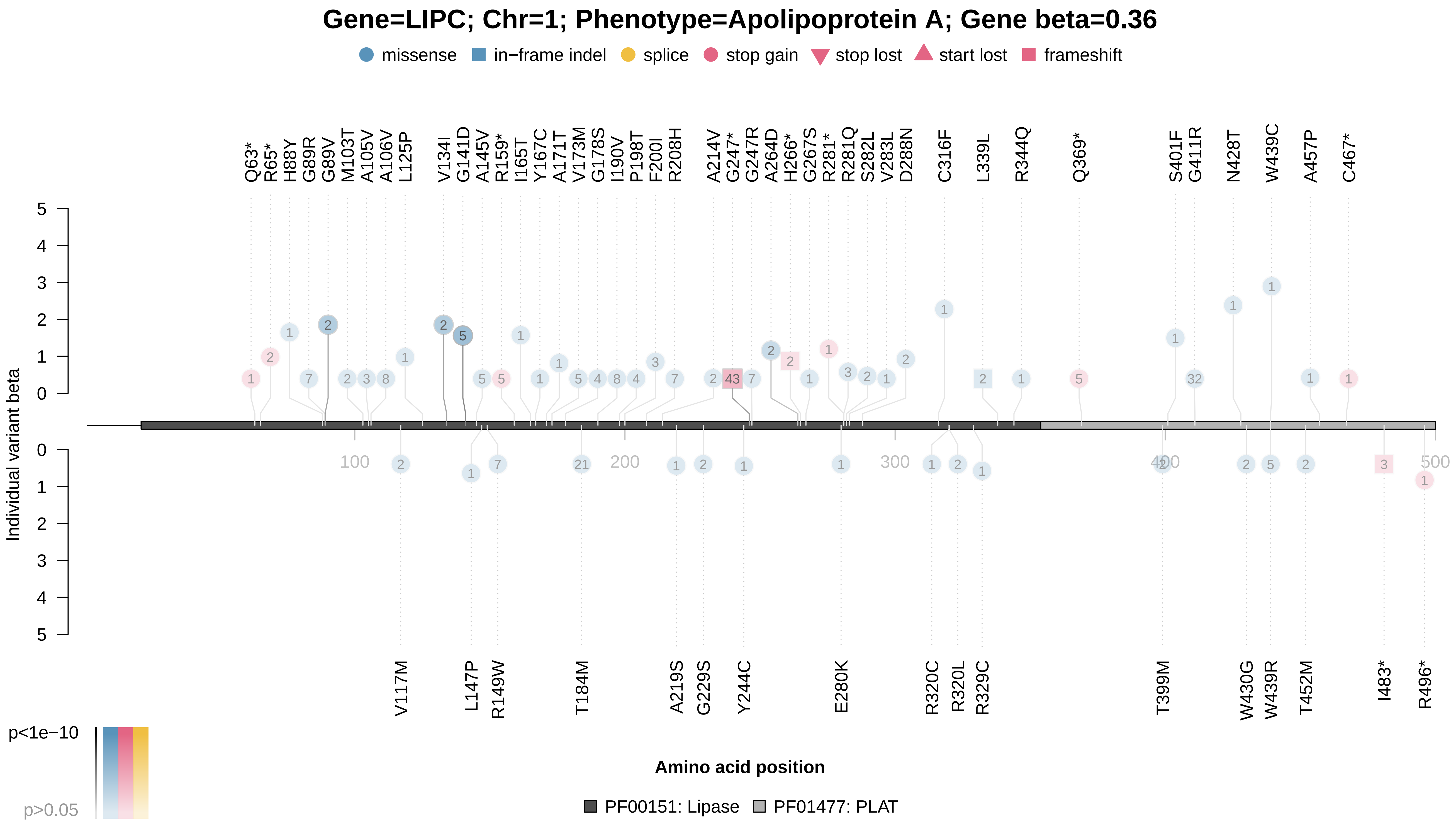

### MC1R_x1747_2.pdf

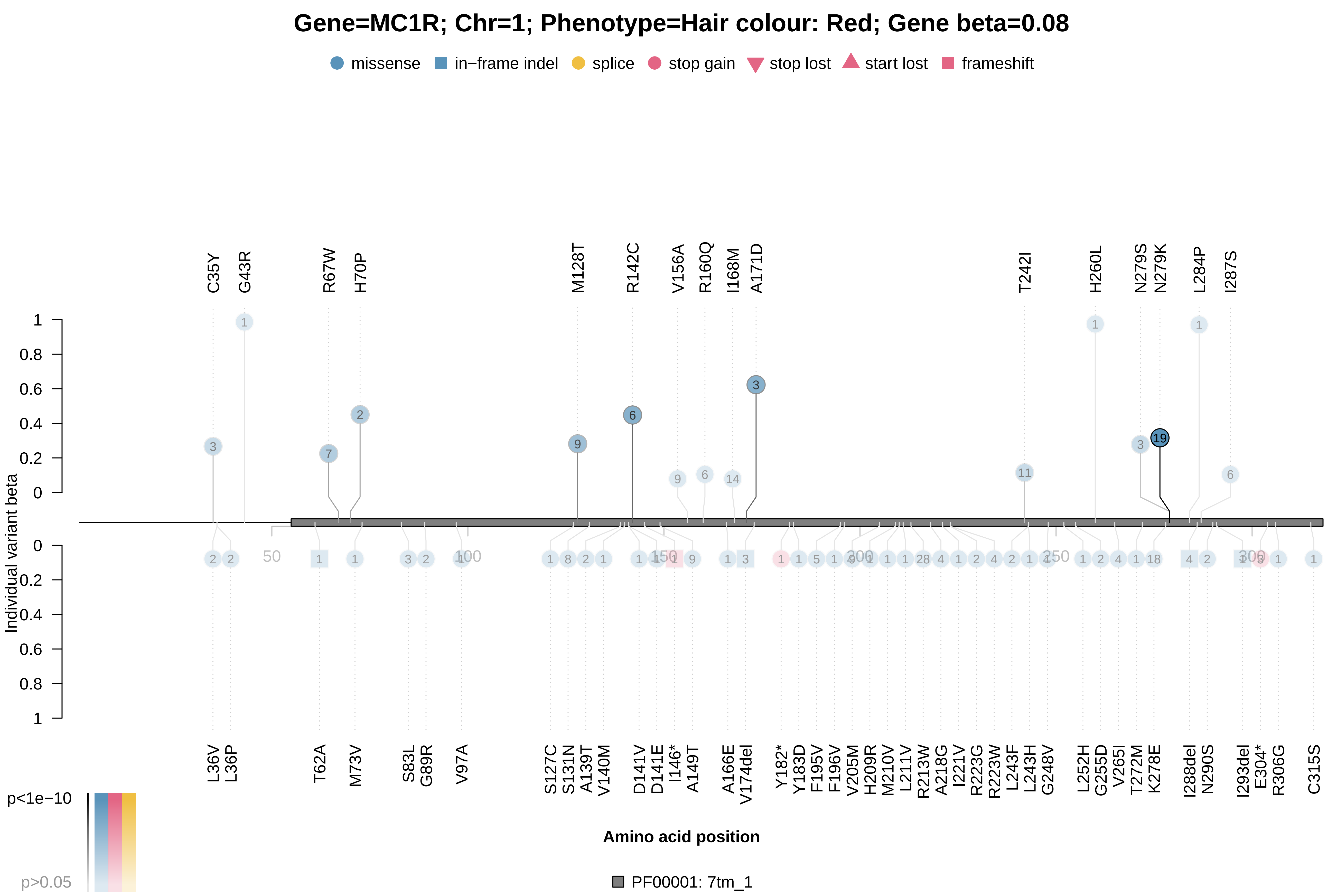

### MPL_x30080.pdf

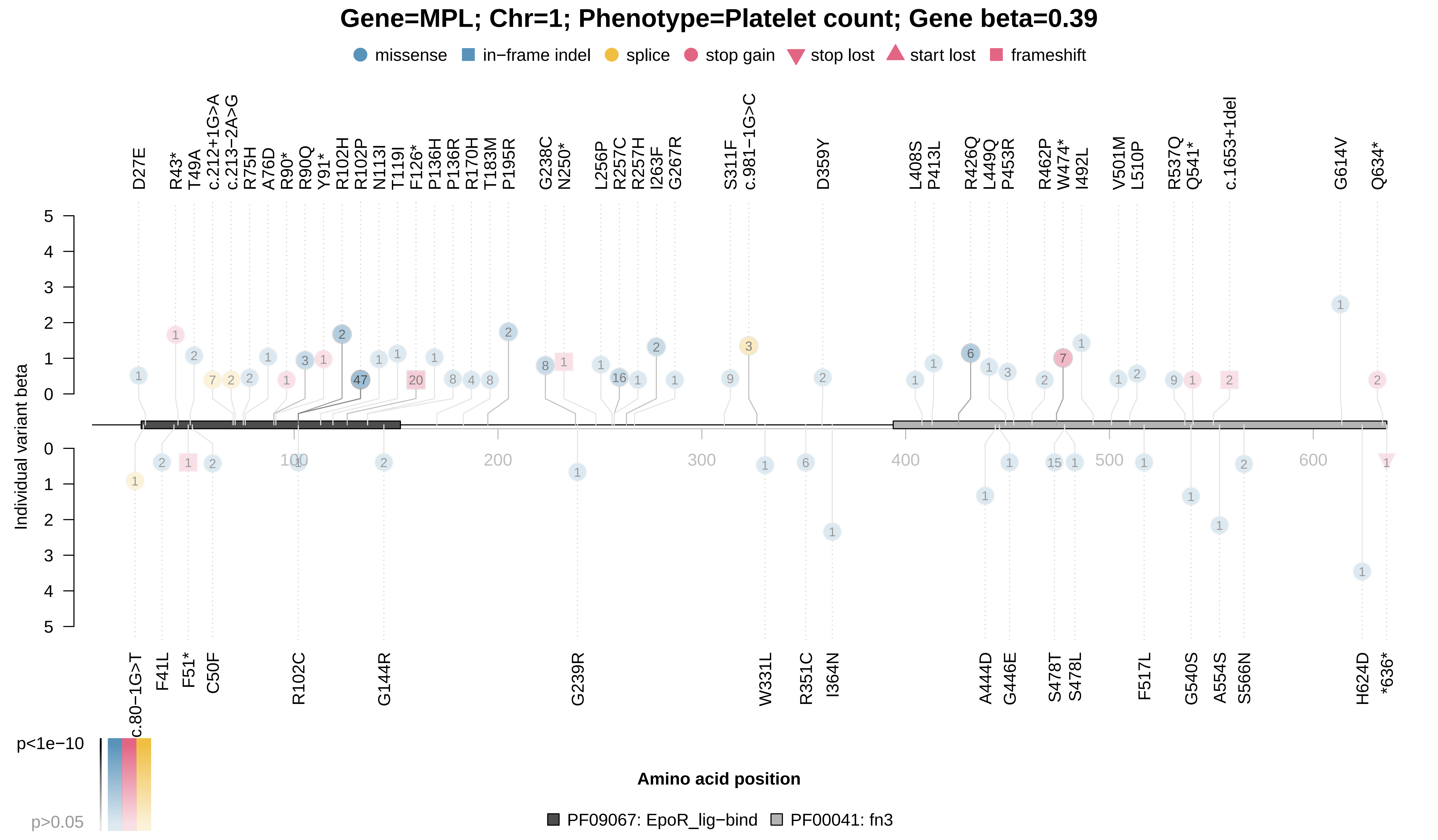

### MPL_x30090.pdf

#### Gene=MPL; Chr=1; Phenotype=Platelet crit; Gene beta=0.33

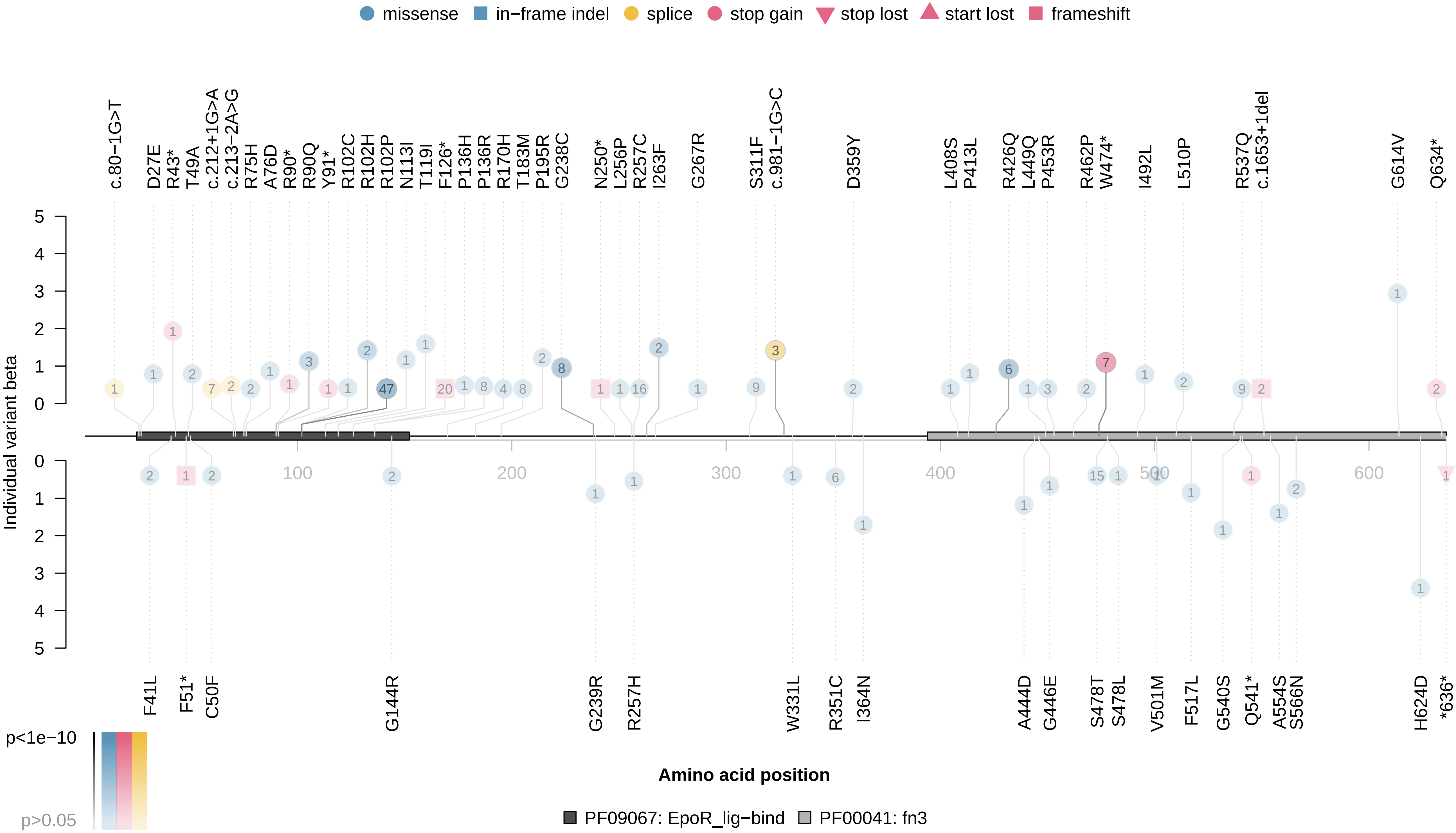

### PAPPA_x50.pdf

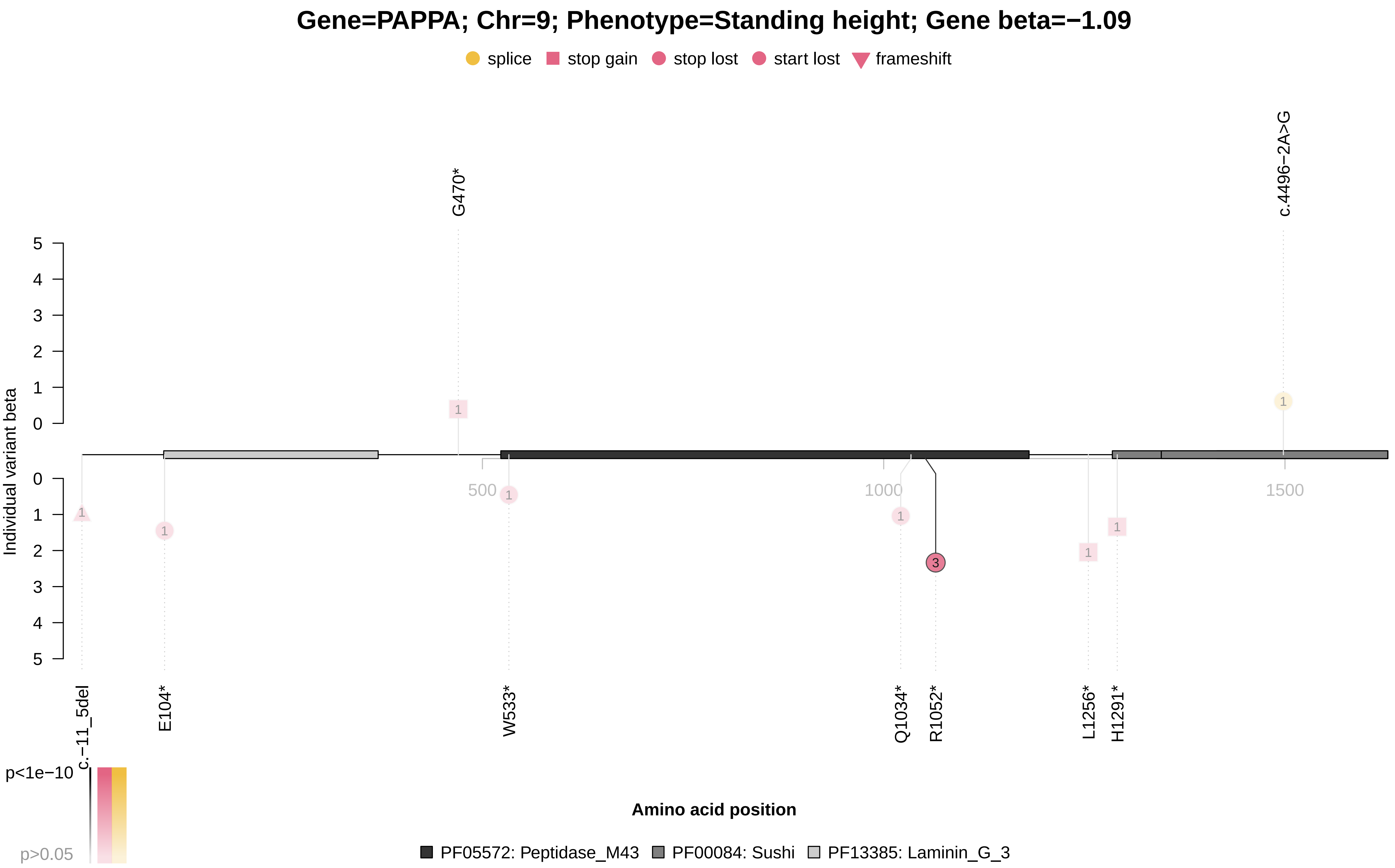

### PCSK9_x30640.pdf

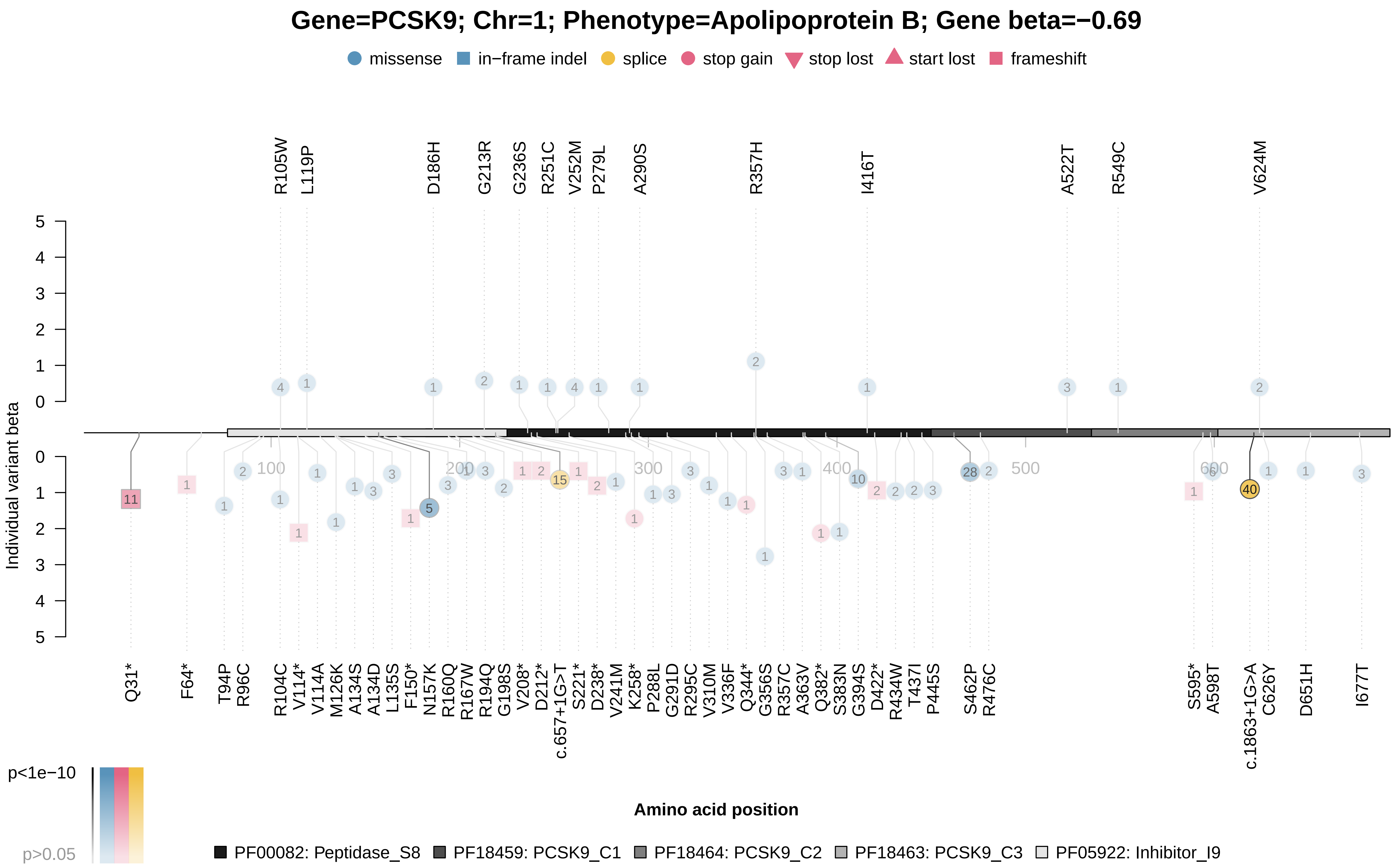

### PCSK9_x30640.pdf

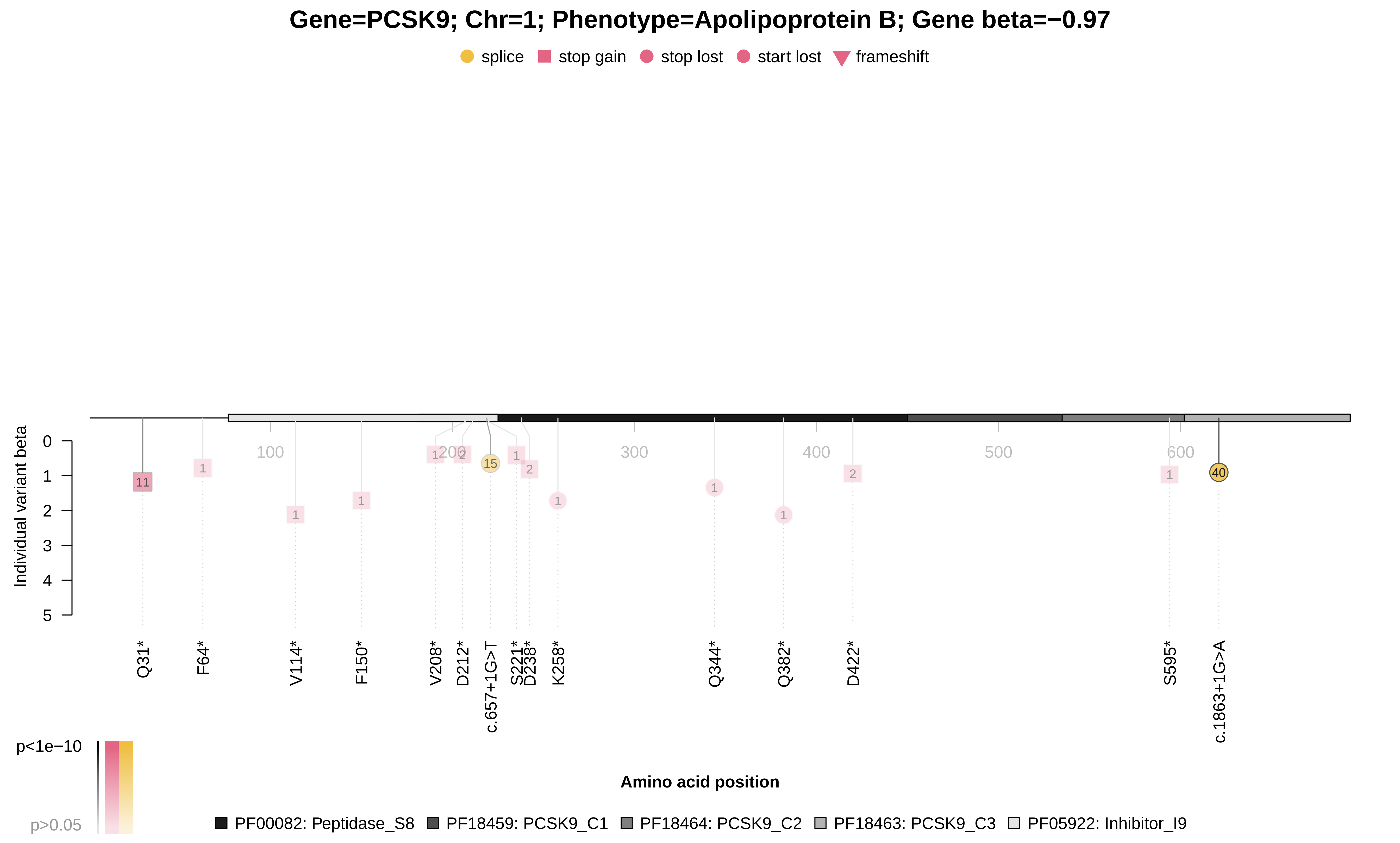

### PCSK9_x30690.pdf

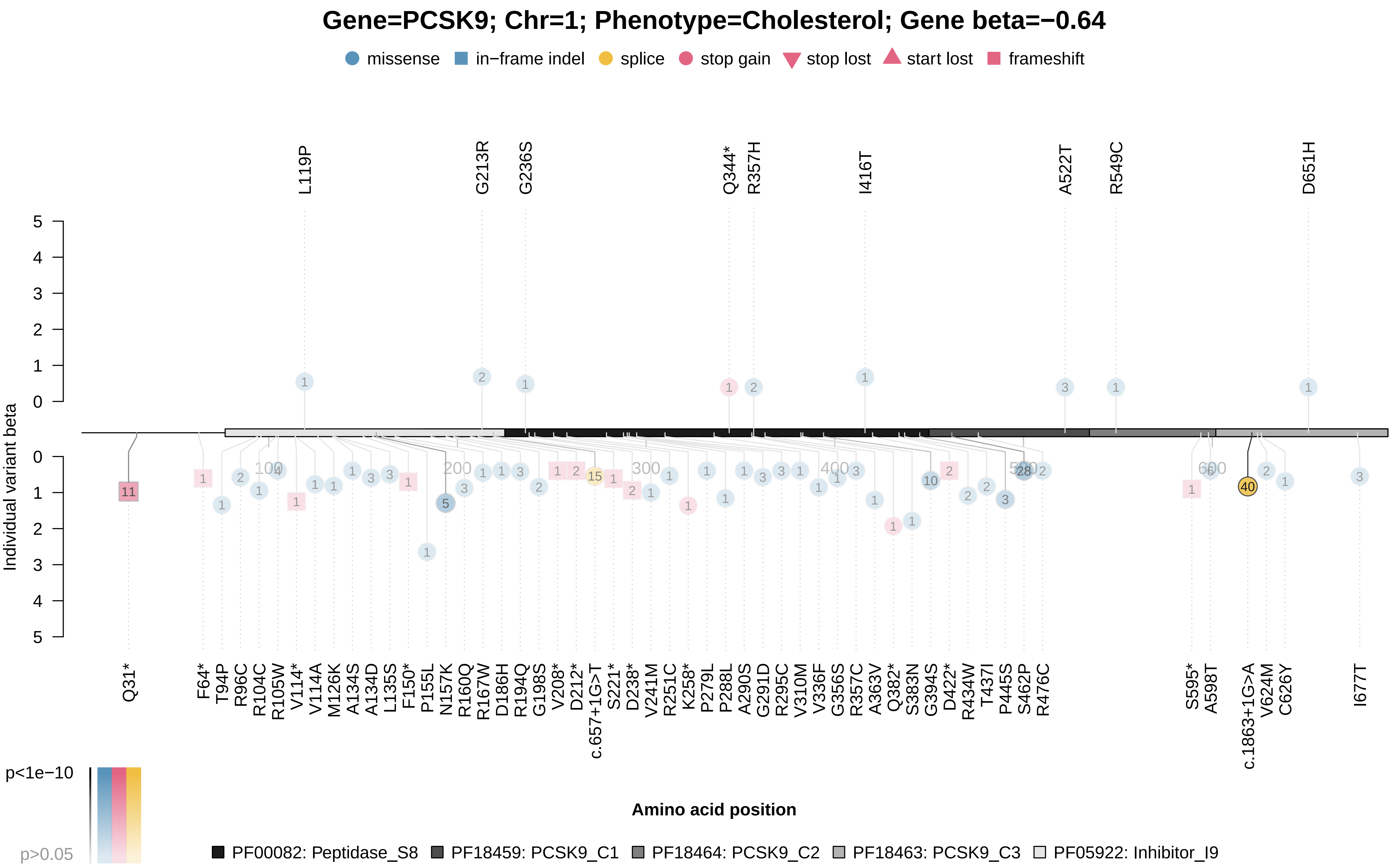

### PCSK9_x30690.pdf

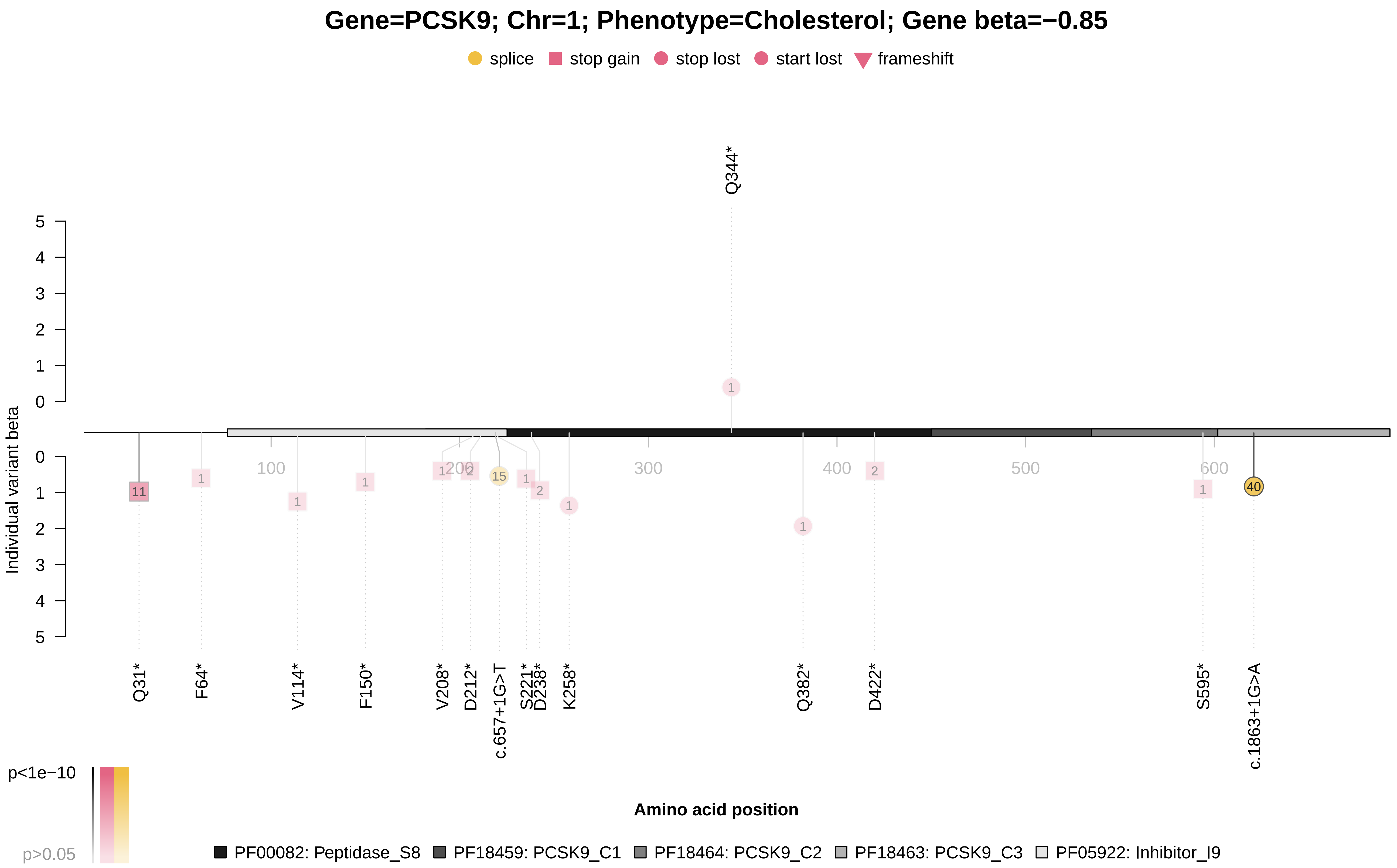

### PCSK9_x30780.pdf

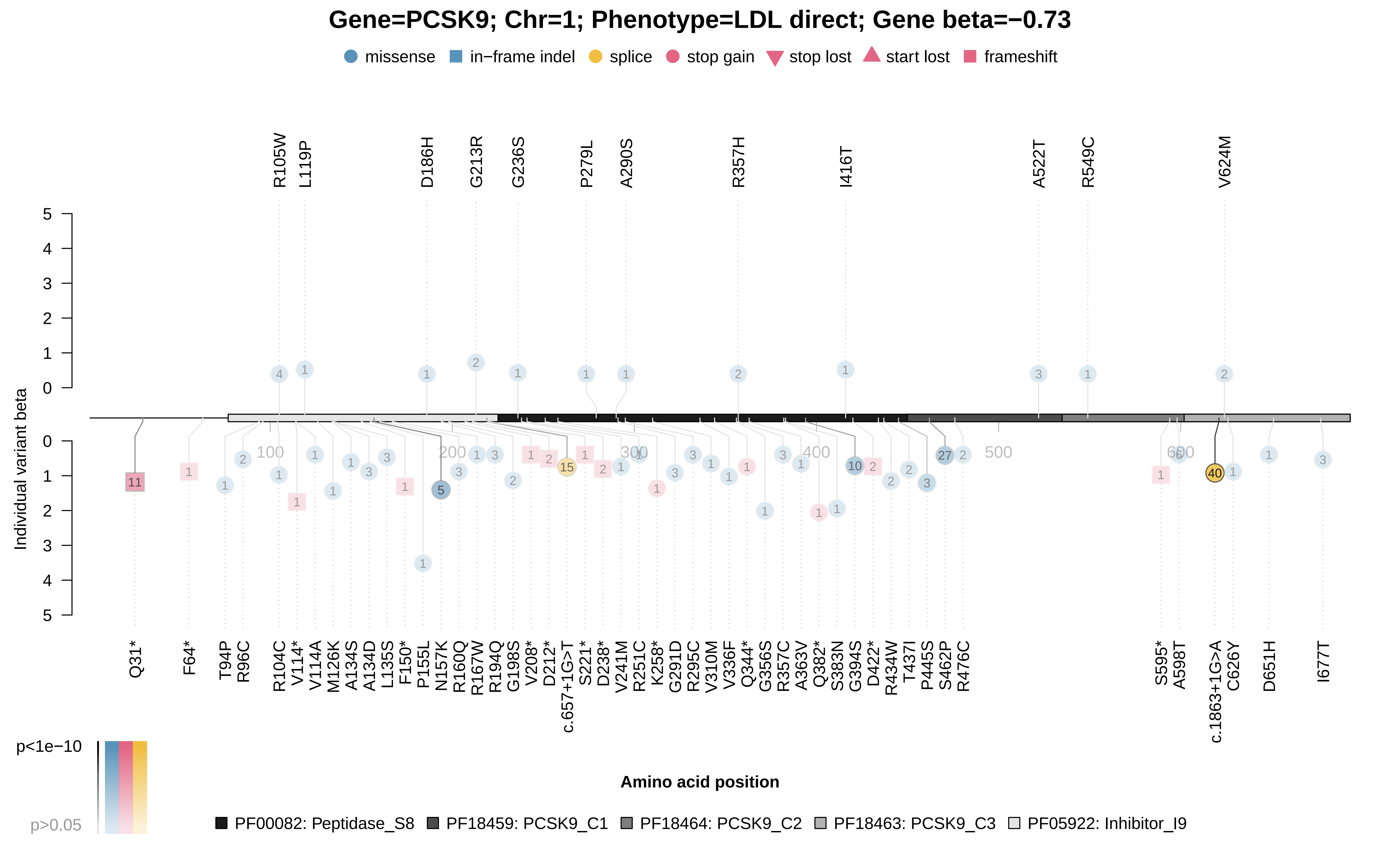

### PCSK9_x30780.pdf

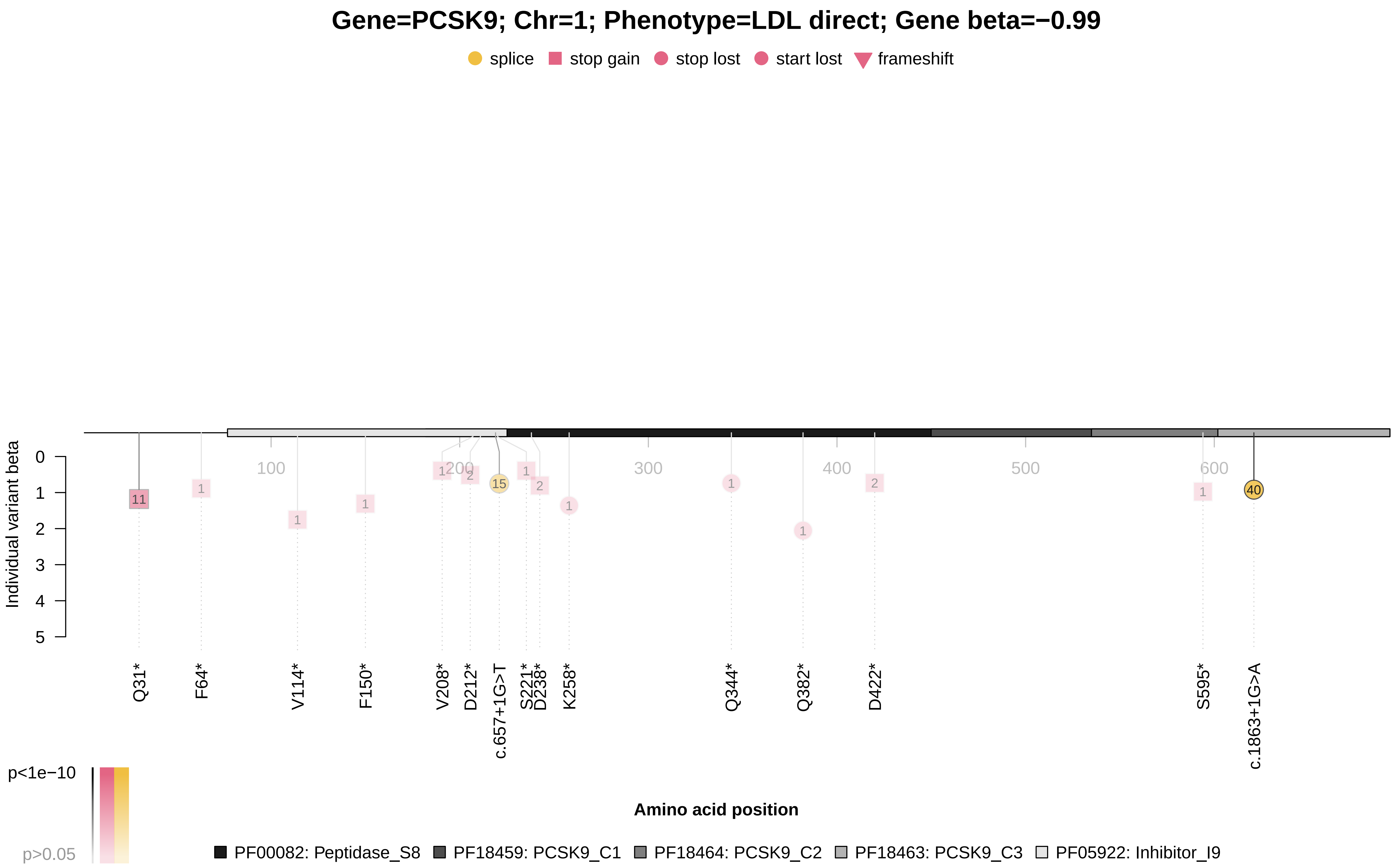

### SCARB1_x30760.pdf

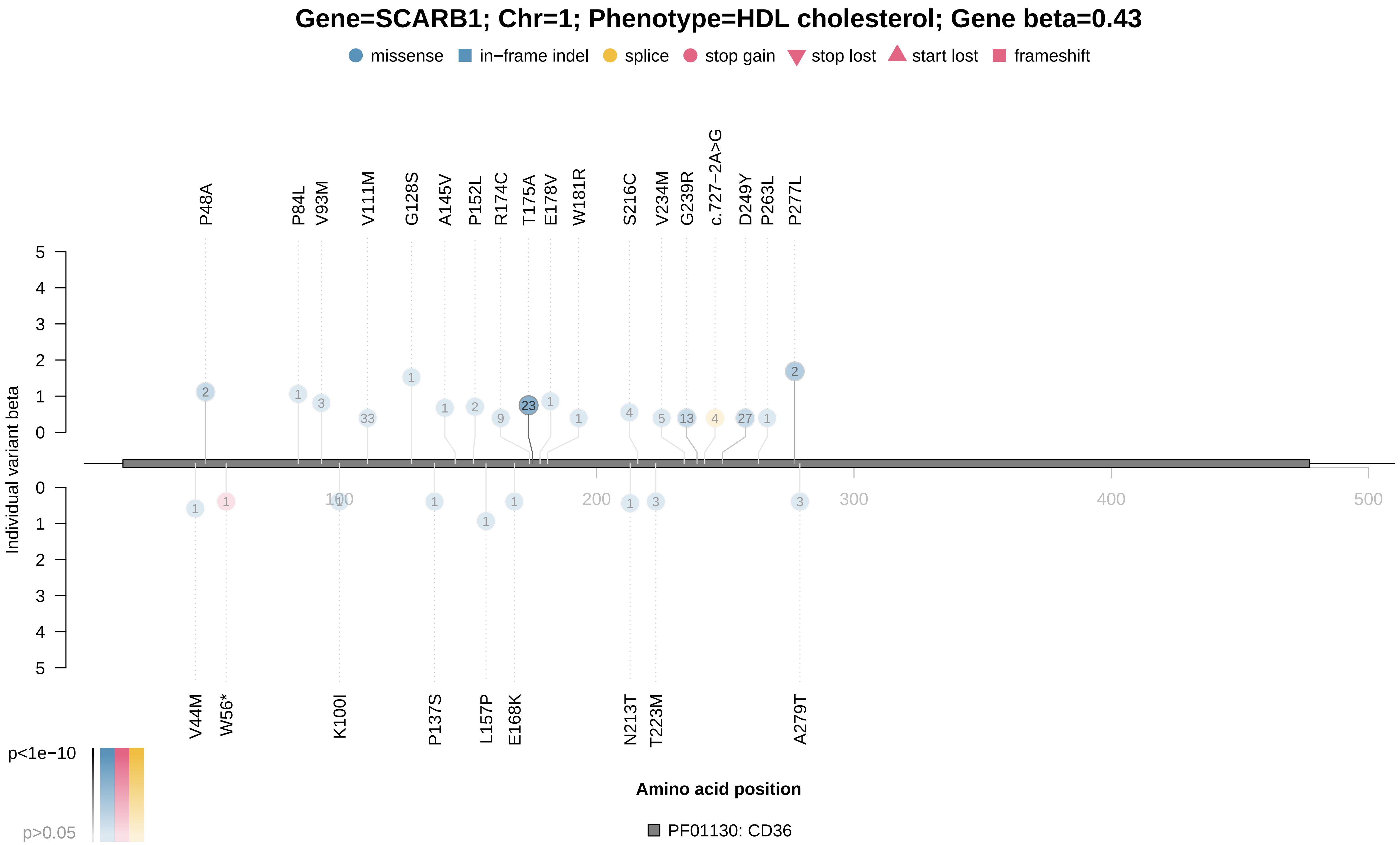

### SHBG_x30830.pdf

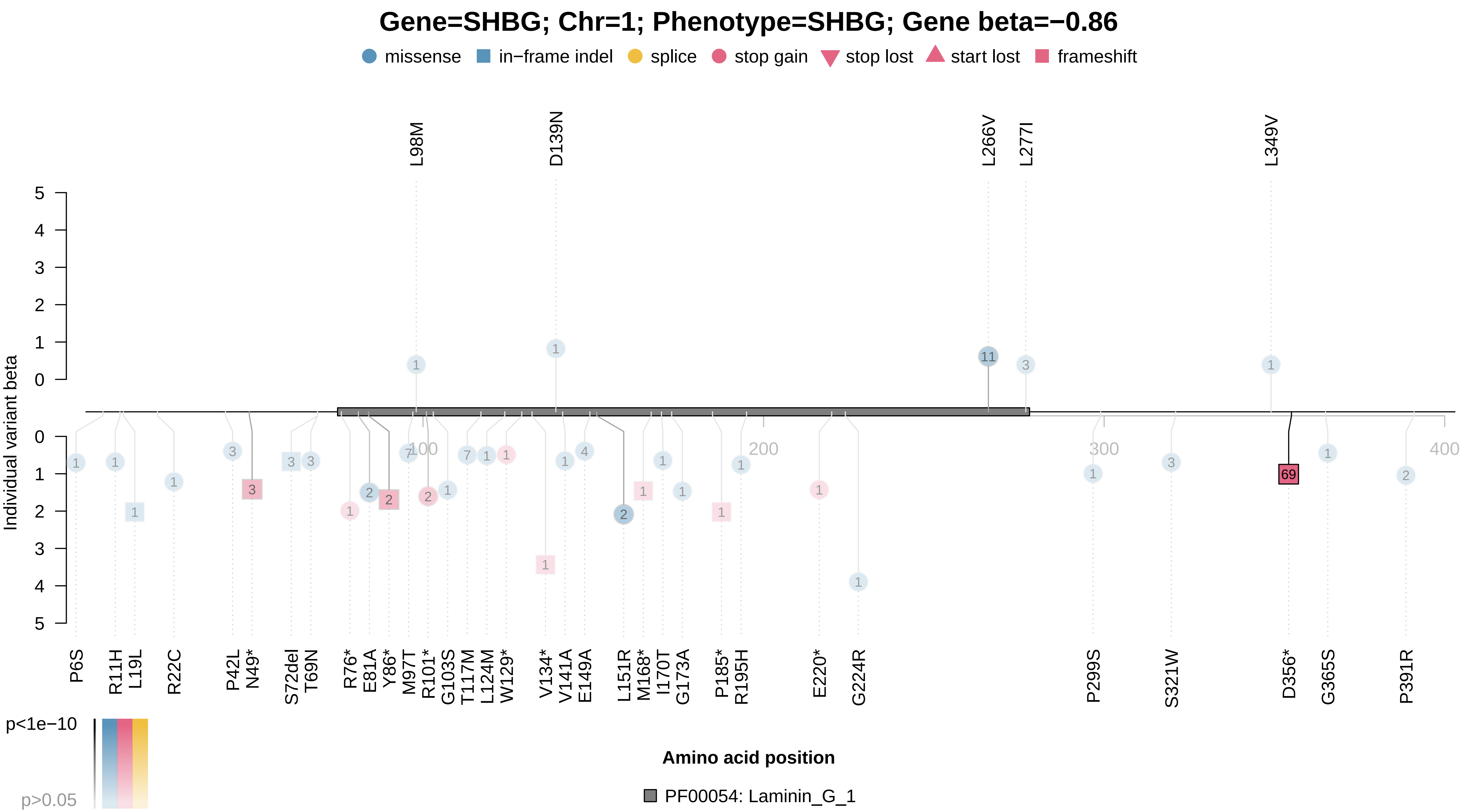

### SHBG_x30830.pdf

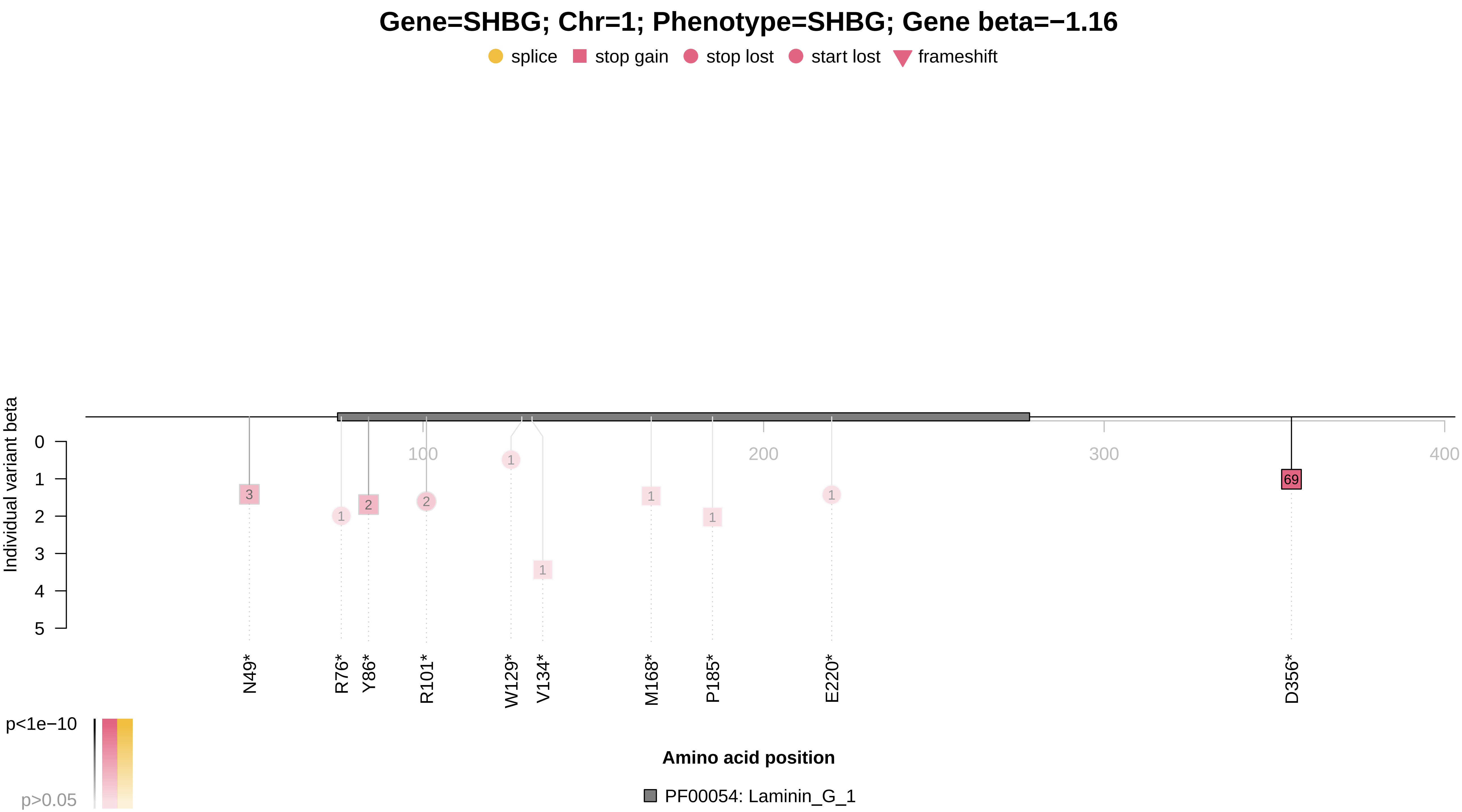

### SLC2A9_x30880.pdf

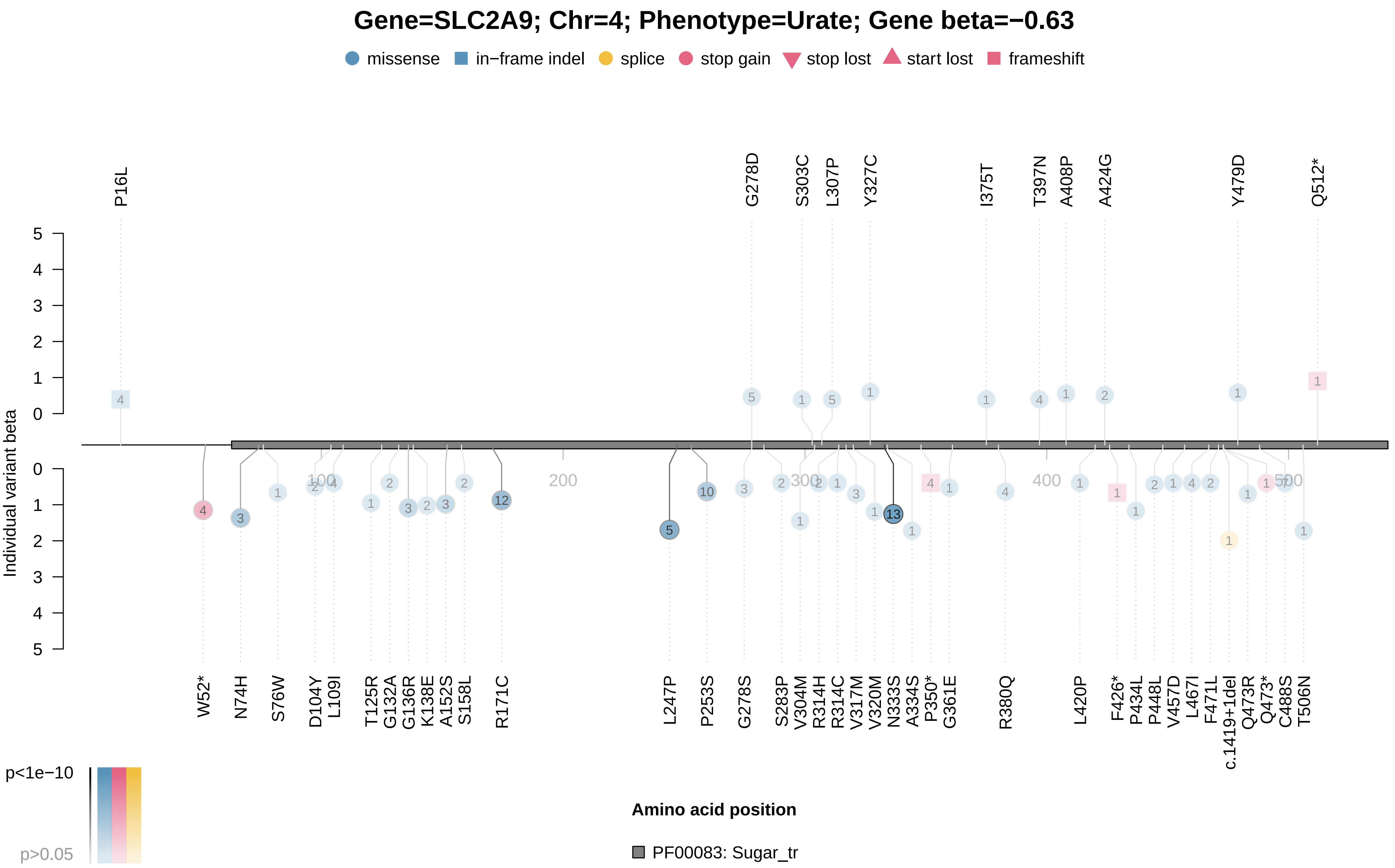

### SLC22A12_x30880.pdf

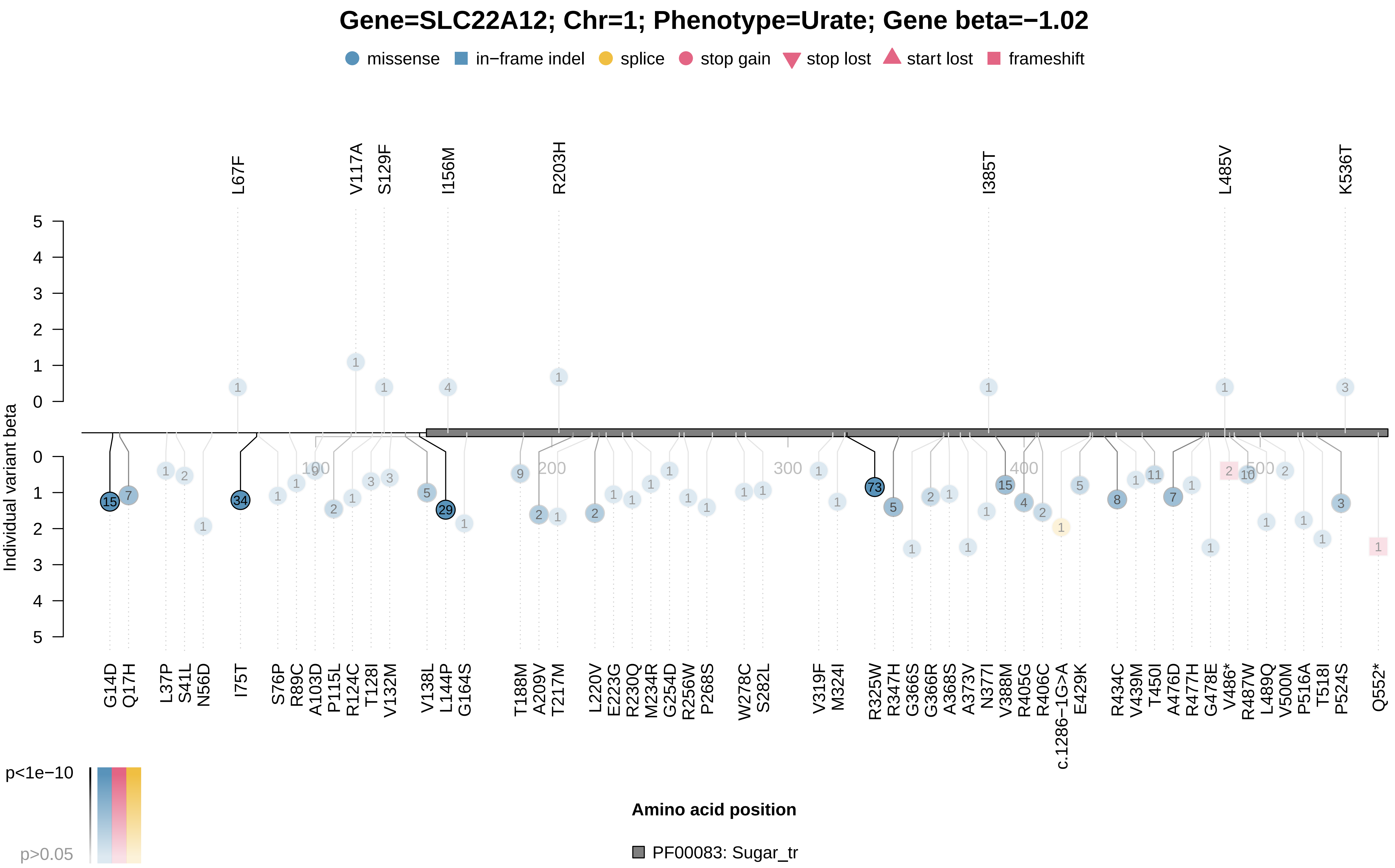

### SLC45A2_x1747_1.pdf

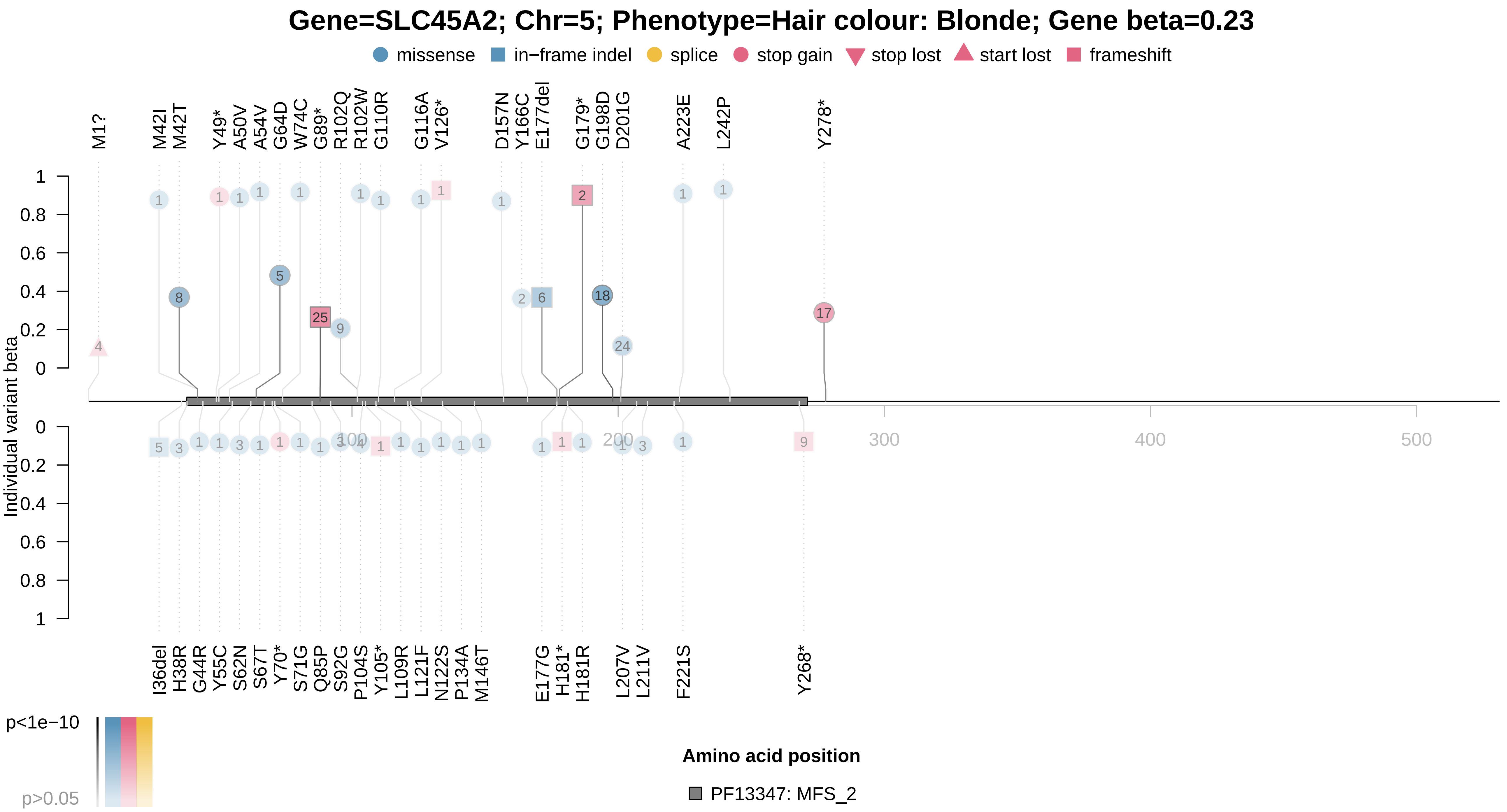

### SLC45A2_x1747_4.pdf

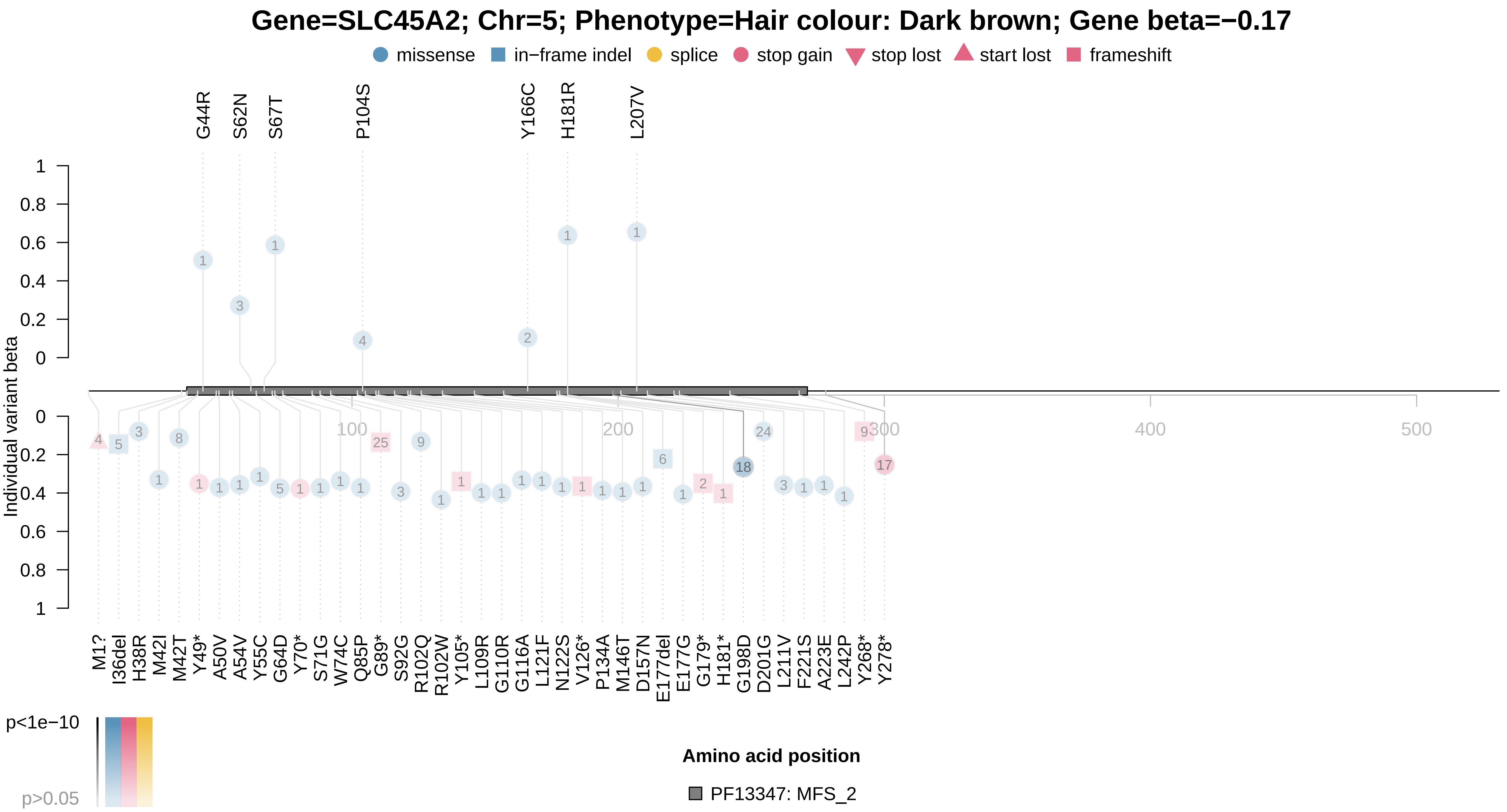

### STAB1_x25030.pdf

##### Gene=STAB1; Chr=3; Phenotype=Median T2star in putamen (left); Gene beta=0.4

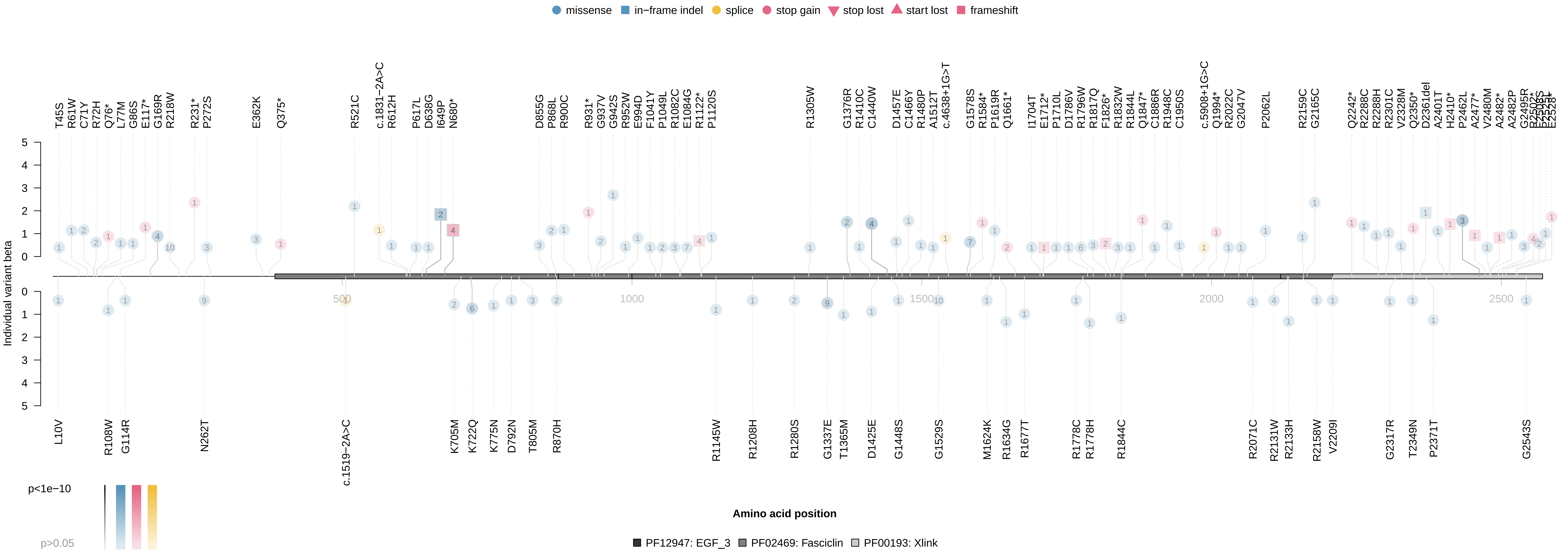

### STAB1_x25030.pdf

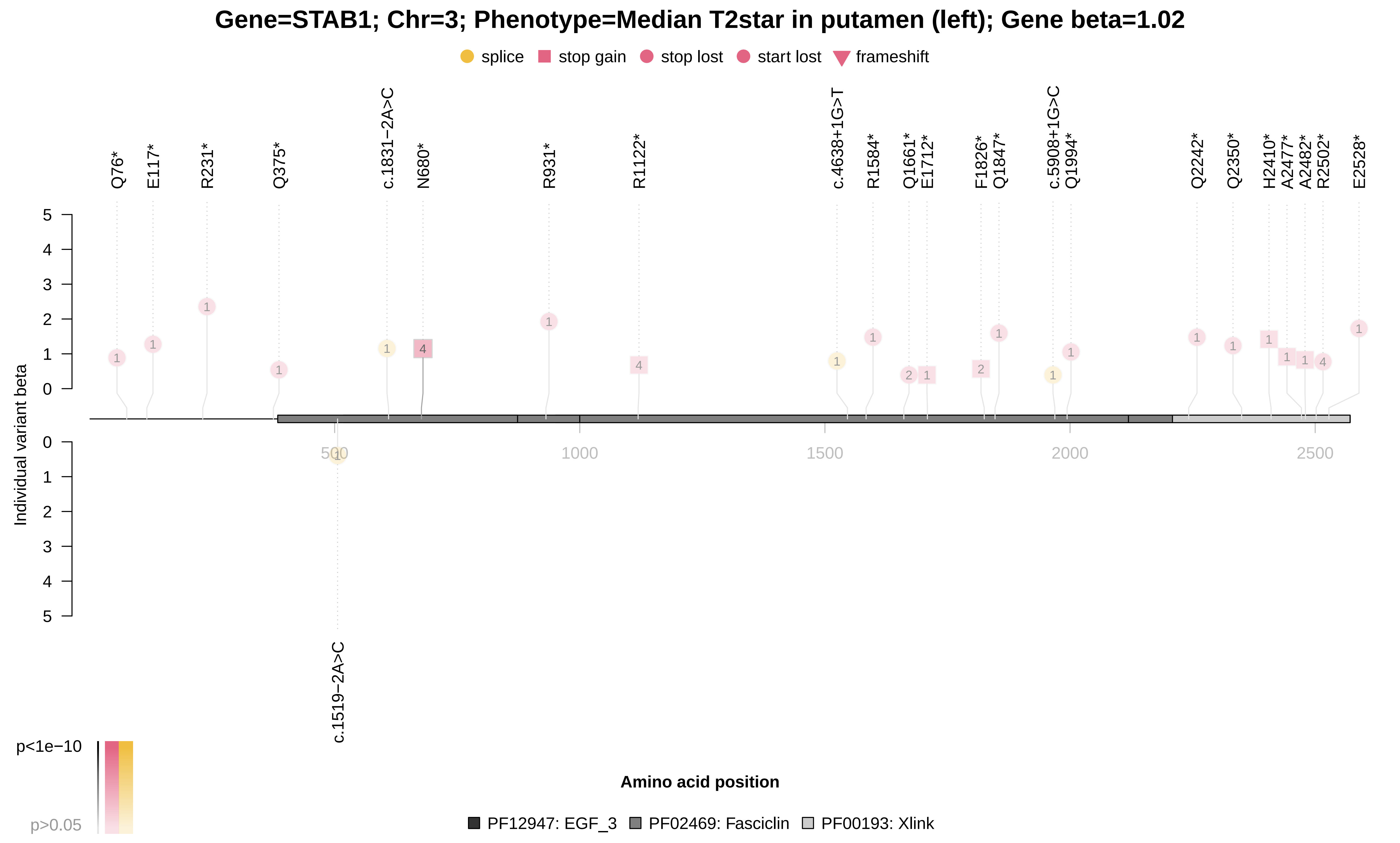

### STAB1_x25031.pdf

##### Gene=STAB1; Chr=3; Phenotype=Median T2star in putamen (right); Gene beta=0.44

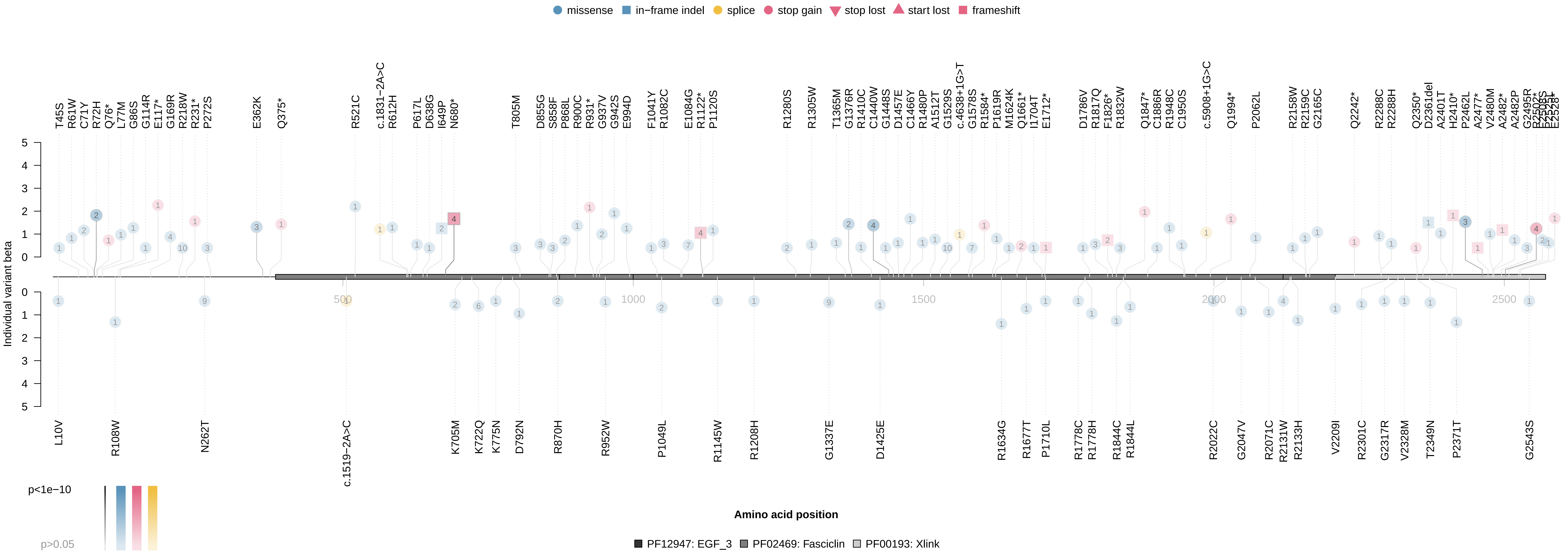

### STAB1_x25031.pdf

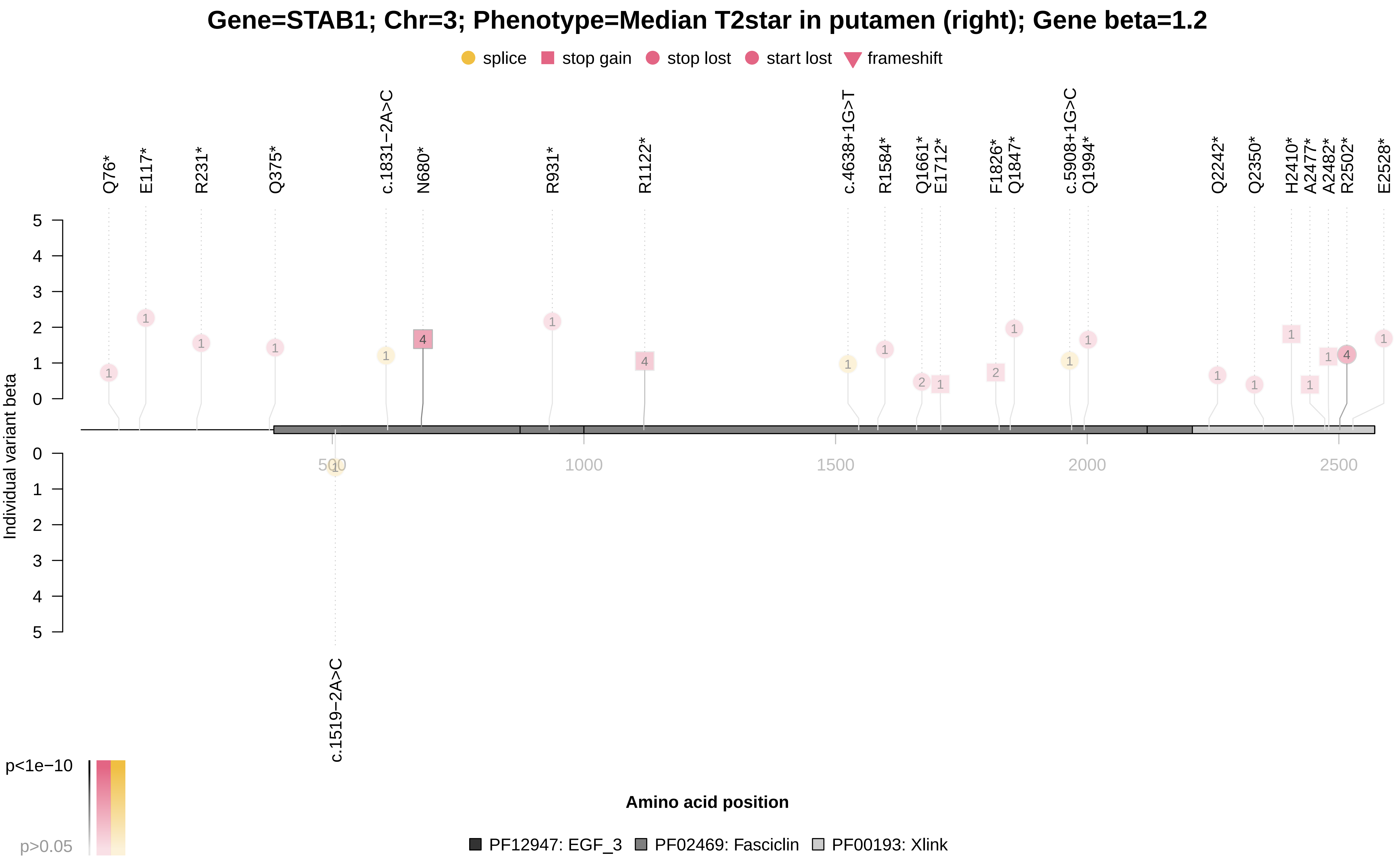

### TTN_I48.pdf

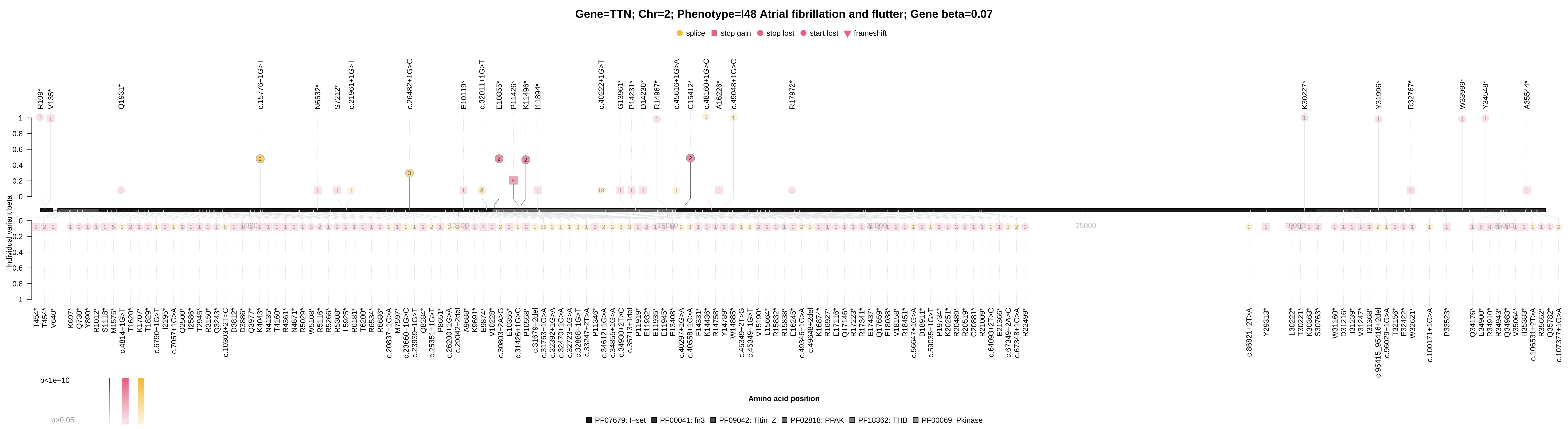

### TUBB1_x30080.pdf

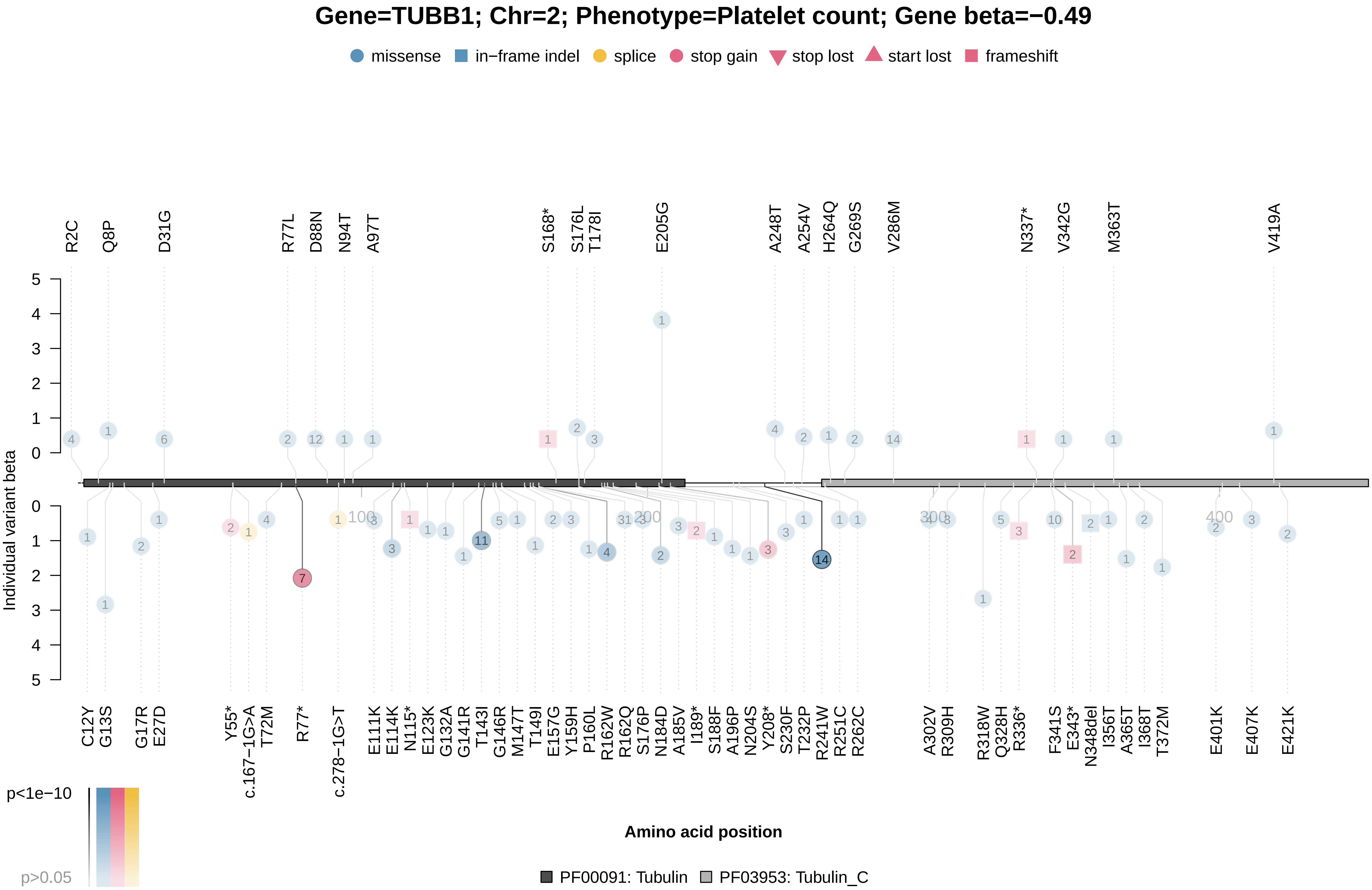

### TUBB1_x30080.pdf

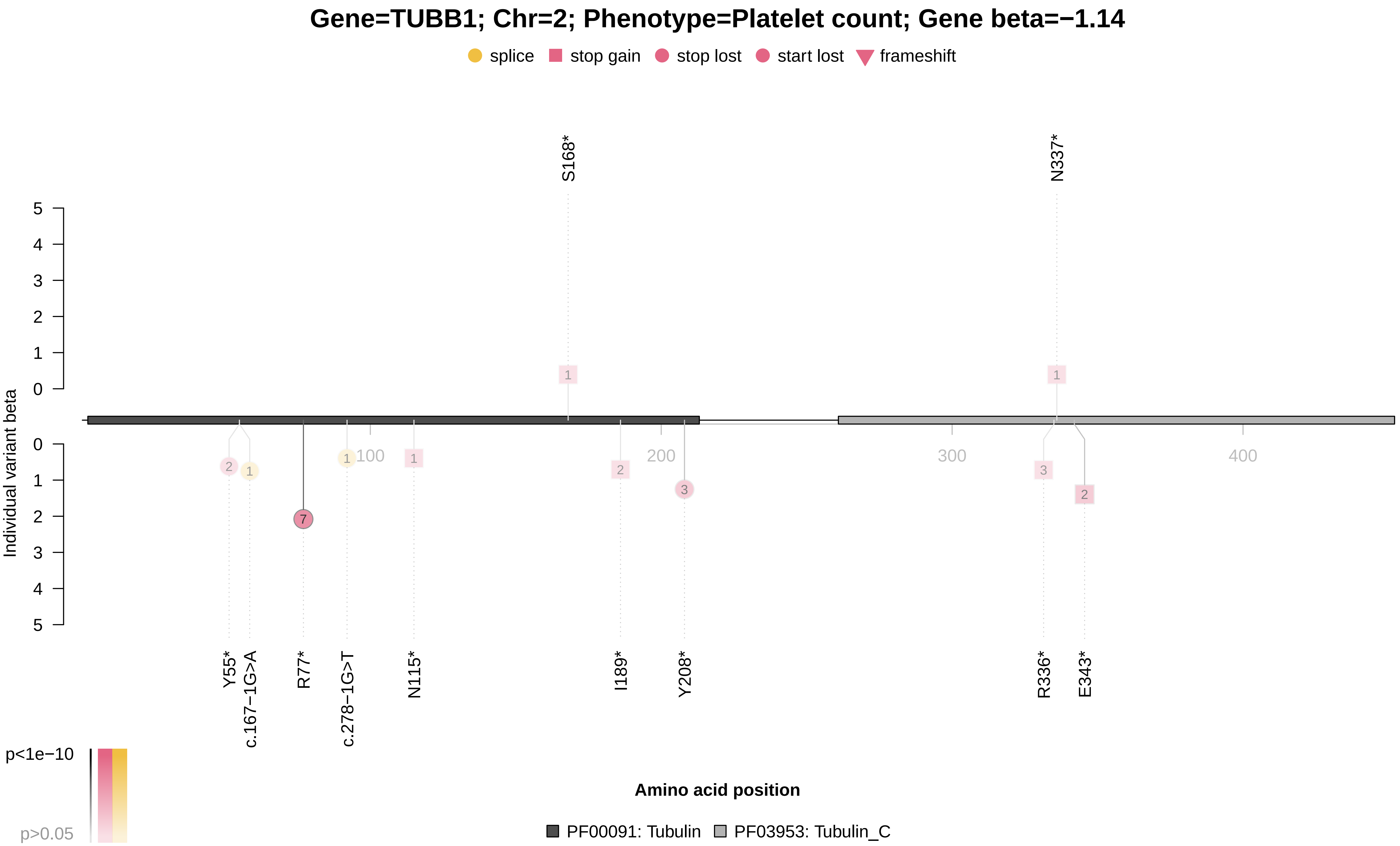

### TUBB1_x30090.pdf

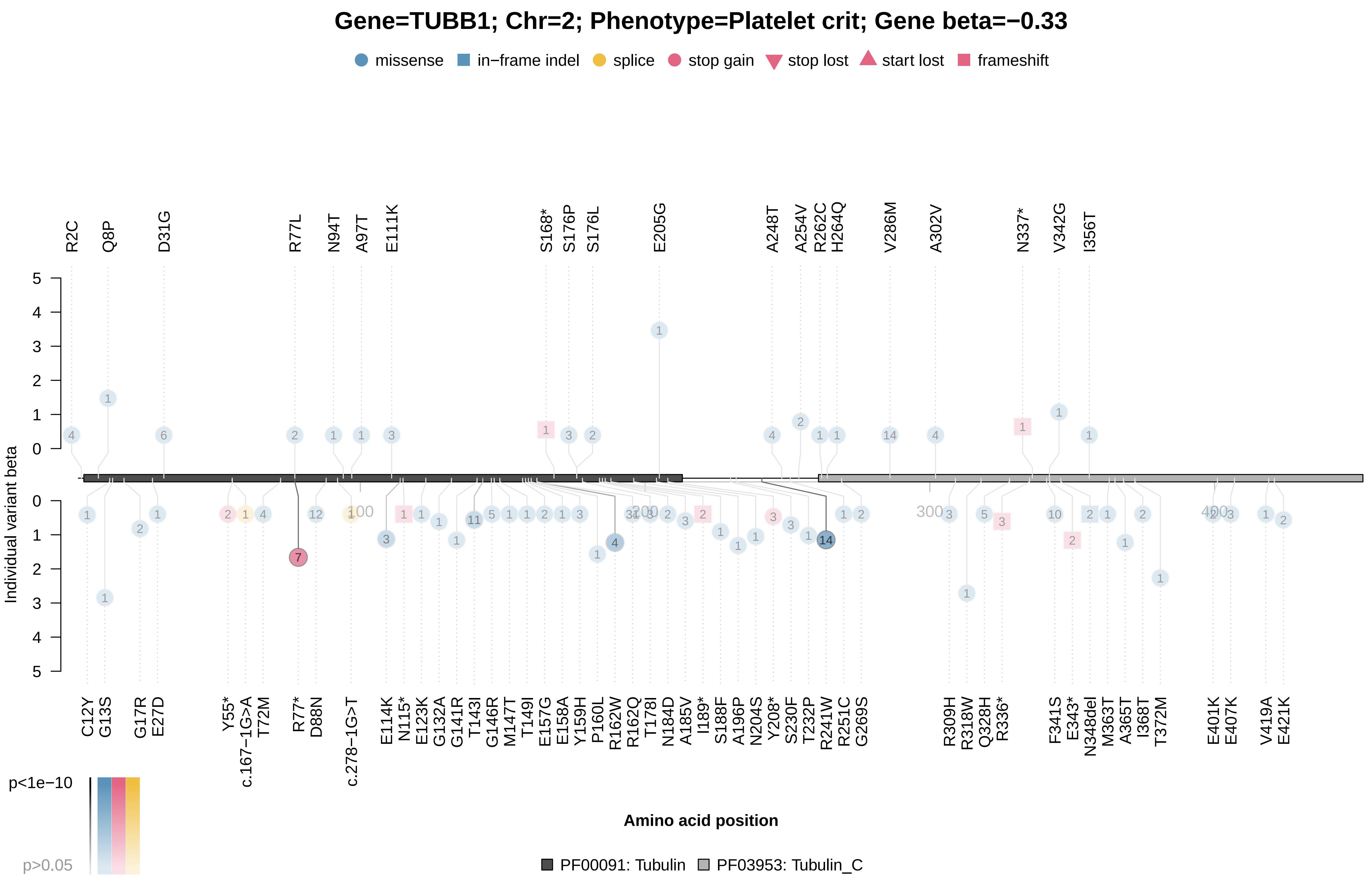

### TUBB1_x30100.pdf

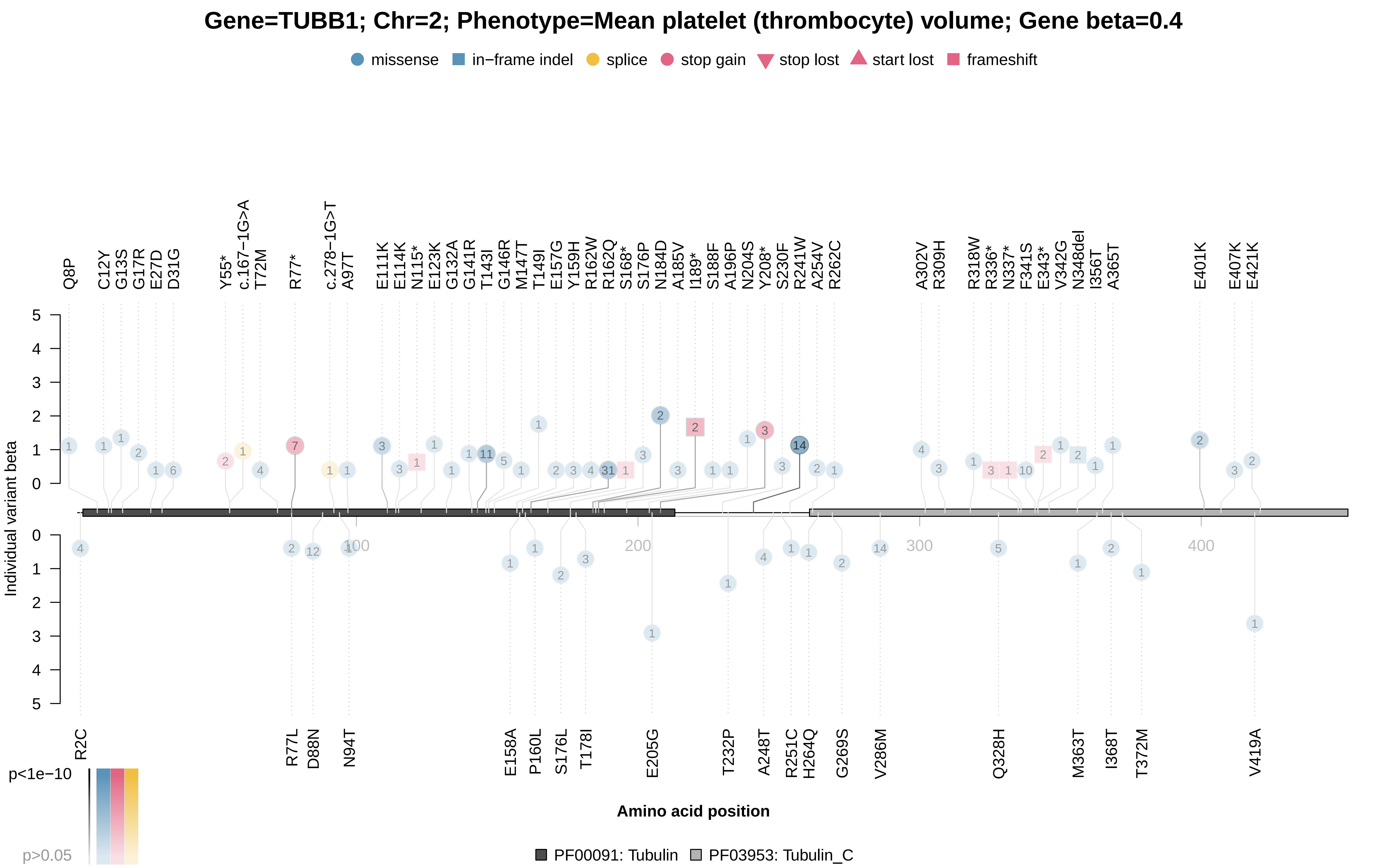

### TUBB1_x30110.pdf

##### Gene=TUBB1; Chr=2; Phenotype=Platelet distribution width; Gene beta=0.85

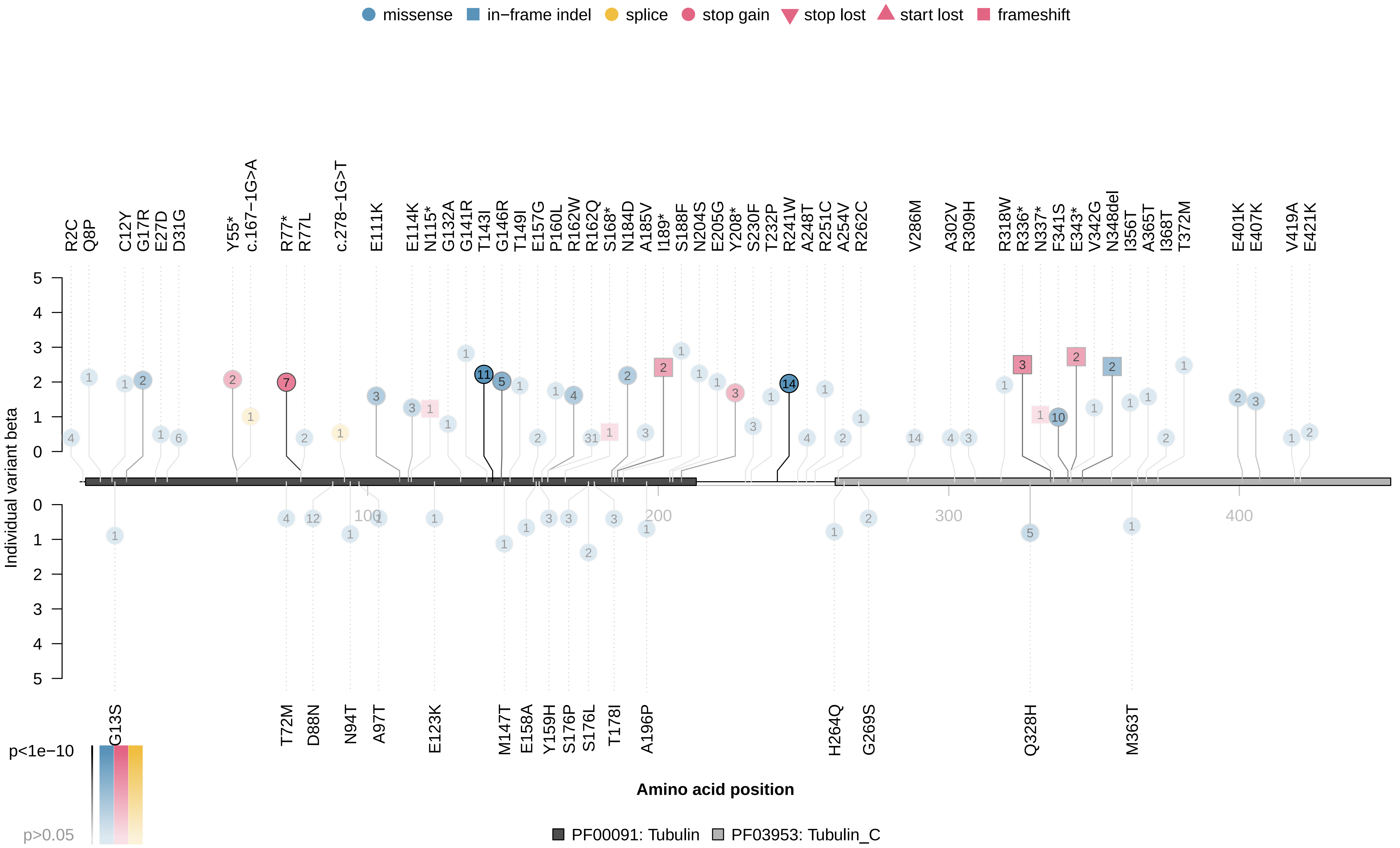

### TUBB1_x30110.pdf

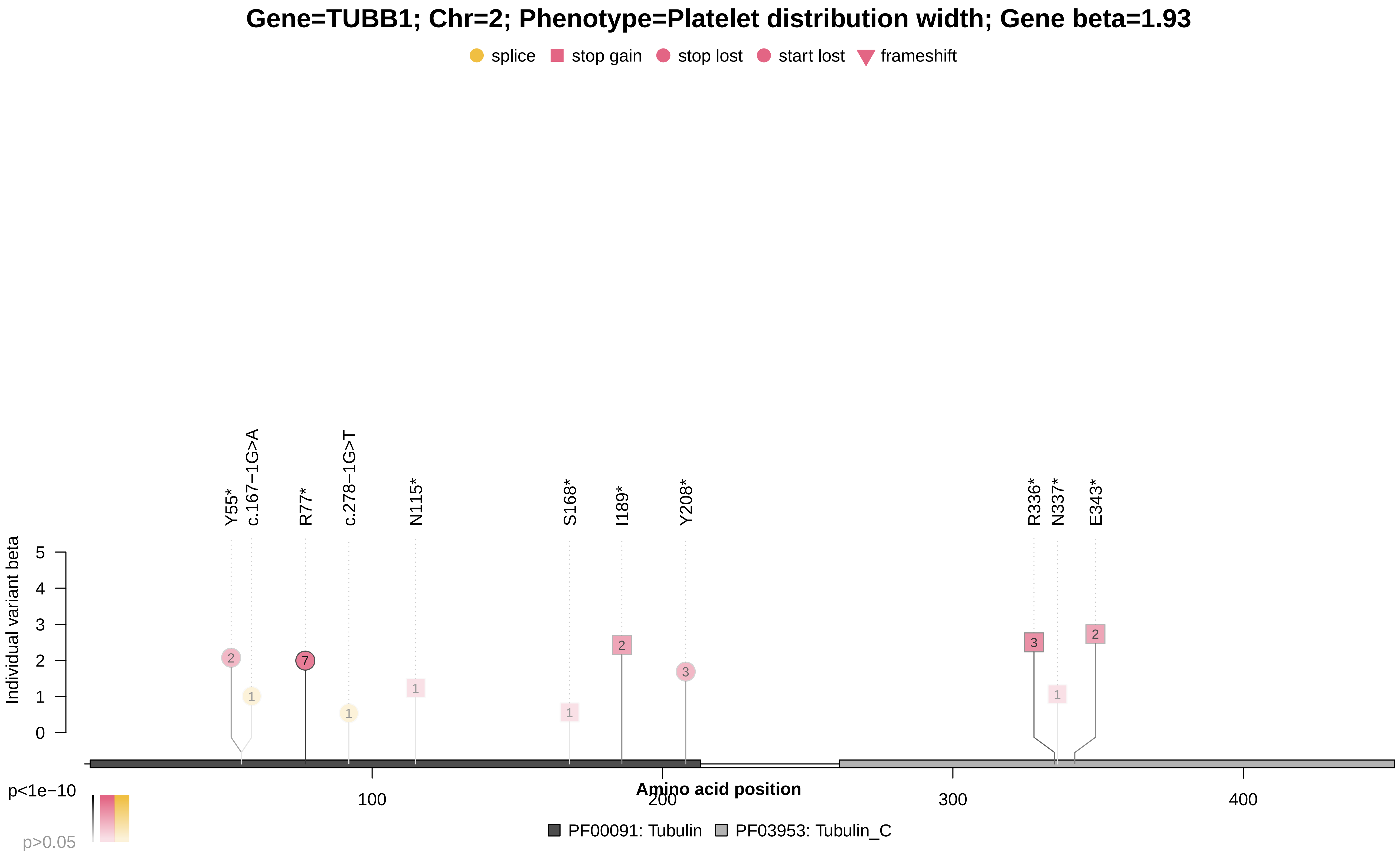

### TYRP1_x1747_1.pdf

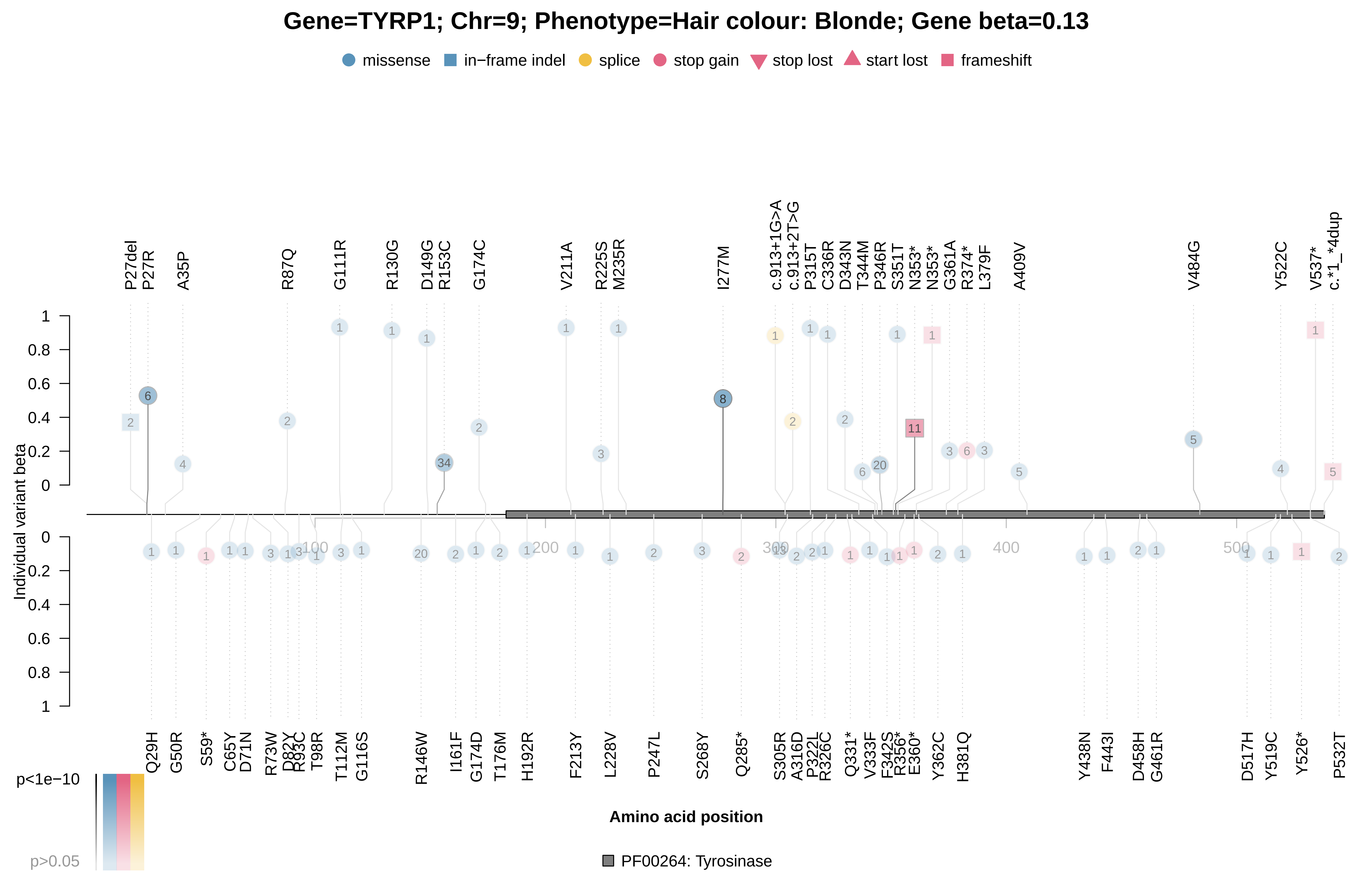
